# Do questionnaires reflect their purported cognitive functions?

**DOI:** 10.1101/583690

**Authors:** Ian A. Clark, Eleanor A. Maguire

## Abstract

Questionnaires are used widely across psychology and permit valuable insights into a person’s thoughts and beliefs, which are difficult to derive from task performance measures alone. Given their importance and widespread use, it is vital that questionnaires map onto the cognitive functions they purport to reflect. However, where performance on naturalistic tasks such as imagination, autobiographical memory, future thinking and navigation is concerned, there is a dearth of knowledge about the relationships between task performance and questionnaire measures. Questionnaires are also typically designed to probe a specific aspect of cognition, when instead researchers sometimes want to obtain a broad profile of a participant. To the best of our knowledge, no questionnaire exists that asks simple single questions about a wide range of cognitive functions. To address these gaps in the literature, we recruited a large sample of participants (n = 217), all of whom completed a battery of widely used questionnaires and performed naturalistic tasks involving imagination, autobiographical memory, future thinking and navigation. We also devised a questionnaire that comprised simple single questions about the cognitive functions of interest. There were four main findings. First, imagination and navigation questionnaires reflected performance on their related tasks. Second, memory questionnaires were associated with autobiographical memory vividness and not internal (episodic) details. Third, imagery questionnaires were more associated with autobiographical memory vividness and future thinking than the questionnaires purporting to reflect these functions. Finally, initial exploratory analyses suggested that a broad profile of information can be obtained efficiently using a small number of simple single questions, and these modelled task performance comparably to established questionnaires in young, healthy adults. Overall, while some questionnaires can act as proxies for behaviour, the relationships between memory and future thinking tasks and questionnaires are more complex and require further elucidation.

## 1. Introduction

Questionnaires are used widely across psychology. They offer the opportunity to gain valuable insights into a person’s subjective thoughts, beliefs and meta-cognition which are not easy to derive from task performance measures alone. Even where performance on cognitive tasks or measures of brain activity are the main focus of investigation, questionnaires nevertheless often also play a role (Arnold, Iaria, & Ekstrom, 2016; Auger, Zeidman, & Maguire, 2017; Belardinelli et al., 2009; Monaghan et al., 2015; Sheldon, Farb, Palombo, & Levine, 2016; Silani et al., 2008). Moreover, in some branches of psychology, questionnaire studies outnumber those involving the direct measurement of cognition and behaviour (e.g. Baumeister, Vohs, & Funder, 2007).

Given the importance of capturing subjective data and the widespread use of questionnaires, it is vital that such instruments map onto the cognitive functions they purport to reflect. The construction and validation of an effective questionnaire often includes consideration of the questions themselves via factor analyses and principal component loadings, examination of internal validity, test-retest reliability and, where possible, confirmatory analyses against existing measures, be they from relevant cognitive tests or from previously-validated questionnaires (Andrade, May, Deeprose, Baugh, & Ganis, 2014; Blazhenkova & Kozhevnikov, 2009; Hegarty, Richardson, Montello, Lovelace, & Subbiah, 2002).

However, when the focus is on naturalistic aspects of cognition, it can be significantly challenging, and not always possible, to test a questionnaire against performance measures. For example, our interest here is in visual imagination (the process of forming and visualising images in the absence of visual input), autobiographical memory (the recall of past events from one’s life), future thinking (imagining future experiences) and spatial navigation (the process of ascertaining one’s position, and planning and following a route). These cognitive functions can be time consuming and resource-heavy to examine and score (e.g. Hassabis, Kumaran, Vann, & Maguire, 2007; Levine, Svoboda, Hay, Winocur, & Moscovitch, 2002; Woollett & Maguire, 2010). This is especially true when large sample sizes are involved. Aspects of these cognitive functions can be assessed quickly and easily in the laboratory using simplified proxy tests, such as memory for a word list in place of autobiographical memory. However, the exact relationship between laboratory-based and naturalistic tasks is often unknown and, in the case of memory, for example, the two task types engage distinct brain networks (McDermott, Szpunar, & Christ, 2009). Thus, if the relationship between the tasks themselves is not clear, then generalising from questionnaires based on proxy tasks to more complex real-world tasks could be problematic.

Given the nature of mental imagery and imagination, questionnaires have been a mainstay of investigation. Across the literature, the classic measure of mental imagery ability is that of vividness. A particularly popular questionnaire is the Vividness of Visual Imagery Questionnaire (VVIQ; Marks, 1973; Marks, 1995), which has been correlated with many measures including neural activity in visual areas of the brain during imagination (Cui, Jeter, Yang, Montague, & Eagleman, 2007; D’Argembeau & Van der Linden, 2006; Finke & Kosslyn, 1980; Hanggi, 1989; Miller et al., 1987). An updated and extended version of the VVIQ is the Plymouth Sensory Imagery Questionnaire (PSIQ; Andrade et al., 2014). As well as measuring the vividness of visual imagery, the PSIQ also examines the vividness of imagery outside of the visual domain. The PSIQ has high reliability and is strongly correlated with the VVIQ (Andrade et al., 2014).

In addition to vividness, an individual’s general use of visual imagery is also often of interest, and can be captured using the Spontaneous Use of Imagery Scale (SUIS; Reisberg, Pearson, & Kosslyn, 2003). The SUIS has good reliability and convergent validity against other imagery questionnaires (e.g. Nelis, Holmes, Griffith, & Raes, 2014), and has predominately been used to study the relationship between imagery and psychopathology (Pearson, Deeprose, Wallace-Hadrill, Heyes, & Holmes, 2013). It has also been argued that there are multiple types of imagery, and this is believed to be captured in the Object-Spatial Imagery and Verbal Questionnaire (OSIVQ; Blazhenkova & Kozhevnikov, 2009). Indeed, when associated with laboratory visual imagery tasks, the subscales of the OSIVQ show different patterns of associations (Blazhenkova & Kozhevnikov, 2009; Sheldon, Amaral, & Levine, 2017; Vannucci, Mazzoni, Chiorri, & Cioli, 2008).

There are, therefore, multiple visual imagery questionnaires, which correlate highly with each other and with laboratory measures of visual imagery ability. However, to the best of our knowledge, for the widely used questionnaires highlighted above, it is currently unknown if they are associated with performance measures of real world imagination (see also McAvinue & Robertson, 2007).

Considering next autobiographical memory, multiple questionnaires have been developed in an attempt to capture memory ability. However, despite their widespread use, the utility of memory questionnaires for assessing memory performance has been questioned. A review of 14 memory questionnaires (one of which we also use here – the Subjective Memory Questionnaire; Bennett-Levy & Powell, 1980) found that while responses to the questionnaires were reliable and stable, there were limited relationships between any of the memory questionnaires and laboratory measures of memory performance (Herrmann, 1982).

No investigations, however, were performed using naturalistic tests of autobiographical memory, which may show different associations, given their recruitment of distinct brain regions (McDermott et al., 2009).

Since the Herrmann (1982) review, new questionnaires have been developed which may better reflect memory ability. The Memory Experience Questionnaire (Sutin & Robins, 2007) focuses on how memories can differ on a number of phenomenological dimensions. These dimensions have been used to examine, for example, how visual perspective relates to other memory phenomenology (Sutin & Robins, 2010), and to assess correlates of memory and positive self-esteem (D’Argembeau & Van der Linden, 2008). However, to the best of our knowledge, there has been no study examining how the dimensions of this questionnaire relate to autobiographical memory recall.

In contrast, the Survey of Autobiographical Memory (SAM; Palombo, Williams, Abdi, & Levine, 2013), contains two subscales relating to memory recall that purport to reflect autobiographical episodic and semantic memory. In support of these subscale definitions, the SAM Episodic subscale (and not the Semantic) was found to correlate with the recollection of scene pictures, and mean scores on this subscale were lower in participants with a history of depression, in line with the literature. However, of key relevance to the current study, in a sub-analysis of 52 participants reported by Palombo et al. (2013), the SAM Episodic subscale was not associated with the number of episodic (‘internal’) details generated during the Autobiographical Interview, one of the most commonly used measures of real-world autobiographical memory recall (Levine et al., 2002). How well the SAM Episodic subscale reflects autobiographical memory recall is, therefore, unclear. Overall, the link between memory questionnaires and autobiographical memory recall is questionable, and requires additional exploration.

While imagery and memory questionnaires are common across the literature, to the best of our knowledge, there are no questionnaires designed specifically to examine future thinking. Instead, questionnaires have, for example, looked at whether individuals are more biased towards thinking about the past, present or future (Zimbardo & Boyd, 1999) or the consequences of future actions (Strathman, Gleicher, Boninger, & Edwards, 1994). However, the SAM questionnaire mentioned above (Palombo et al., 2013) also contains a “Future subscale”, with items believed to represent future thinking ability and phenomenology. However, while these items were separated from that of the other subscales in the SAM by a multiple correspondence analysis (a form of factor analysis), the Future subscale is yet to be associated with any measure of future thinking.

In contrast to imagination, autobiographical memory and future thinking, navigation questionnaires have been well validated against real world navigation tasks. One of the most popular is the Santa Barbara Sense of Direction Scale (Hegarty et al., 2002). This questionnaire was initially validated in a large sample of over 200 participants against various measures of their sense of direction in environments learned from direct exposure, virtual environments on a computer screen, and videotaped real world environments (Hegarty, Montello, Richardson, Ishikawa, & Lovelace, 2006; Hegarty et al., 2002). A strong relationship between the Santa Barbara Sense of Direction Scale and navigation ability tested in real and virtual worlds has also been replicated across multiple other studies, albeit often in smaller samples (e.g. Hund & Padgitt, 2010; Ishikawa & Montello, 2006; Weisberg, Schinazi, Newcombe, Shipley, & Epstein, 2014).

The SAM questionnaire, mentioned above (Palombo et al., 2013), also has a fourth subscale designed to represent spatial ability. In an online study of over 300 participants familiar with downtown Toronto, scores on the SAM Spatial subscale significantly correlated with the ability to judge distances and spatial relationships between downtown Toronto landmarks, while the Episodic, Future and Semantic subscales did not (Selarka, Rosenbaum, Lapp, & Levine, 2019). Both the Santa Barbara Sense of Direction Scale and the Spatial subscale of the SAM seem, therefore, to reflect their purported cognitive function of navigation.

In summary, when considering imagination, autobiographical memory and future thinking, there is a surprising lack of validation of questionnaire data against naturalistic tasks. This is despite the questionnaires being widely used in the literature as a proxy for real-world behaviour. By contrast, navigation questionnaires, in particular the Santa Barbara Sense of Direction Scale, have been widely validated against real-world navigation, showing a strong relationship.

The focus of the current study is the relationship between performance on naturalistic cognitive tasks and questionnaire measures purporting to reflect the cognitive functions being assessed in those tasks. In a recent study, involving the same participants that we report on here, we examined the relationships between imagination, autobiographical memory, future thinking and navigation in terms of task performance (Clark et al., in press). We found that scores across the four tasks were related, although navigation was more closely aligned with other (laboratory-based) spatial tasks. In that study we also tested whether or not there was a common cognitive process underpinning performance on these tasks. It has been suggested that autobiographical memory provides the building blocks for thinking about the future and imagining fictitious atemporal events (e.g. Moscovitch, Cabeza, Winocur, & Nadel, 2016; Schacter et al., 2012). An alternative proposition is that imagination, recalling the past, thinking about the future and navigation require the mental construction of scene imagery (Hassabis & Maguire, 2007; Maguire & Mullally, 2013). In this context, a scene is a naturalistic three dimensional spatially coherent representation of the world typically populated by objects and viewed from an egocentric perspective. Using mediation analyses, we identified that scene construction, and not autobiographical memory, was a key process linking performance on the four tasks (Clark et al., in press). This association was stronger between imagination, autobiographical memory and future thinking than with navigation. However, the sole focus of Clark et al. (in press) was on the task performance data. The question arises, therefore, as to whether a similar influence of visual (scene) imagery would also be evident in questionnaires purporting to reflect the cognitive functions underpinning these tasks.

Several previous studies have alluded to a crossover between imagery questionnaires and autobiographical memory, future thinking and navigation. Comparing high and low imagery-reporting participants, Vannucci et al. (2016) found that high imagery participants generated a greater number of autobiographical memories, and scored these memories more highly on subjective ratings of vividness and richness of detail. Additionally, D’Argembeau and Van der Linden (2006) observed that self-reported imagery correlated with subjective reports of visual and sensory details when thinking about both past and future events. Interestingly, two participants who claimed to have a complete lack of visual imagery also reported a low sense of reliving when recalling autobiographical memories (Greenberg & Knowlton, 2014). For navigation, a more complex picture has emerged, where benefits have been apparent for participants reporting higher visual over verbal abilities, but also for those with the opposite pattern (Kraemer et al., 2017; Pazzaglia & Moè, 2013).

In summary, how questionnaires designed for one of our cognitive functions of interest relate to the other cognitive functions is unknown, and further investigation is required to identify whether there are, hitherto hidden, links between them.

A separate potential issue with questionnaires is their scope. Researchers often want to obtain a broad profile of a participant. However, questionnaires are typically designed to probe a specific aspect of cognition (e.g. imagery vividness: Andrade et al., 2014; navigation: Hegarty et al., 2002). As such, questionnaires ask multiple questions about the same cognitive function or behaviour. A standard questionnaire typically takes around 5-10 minutes to complete. Therefore, if seeking a wide-ranging profile over multiple aspects of cognition (and thus multiple questionnaires), a battery of questionnaires can easily take an hour. Yet, if a general overview is what is required, then in-depth information on each topic is not necessary. Instead, could a set of single questions on each topic (e.g. “Please rate your ability to construct a mental image”; “Please rate your navigational ability”) provide sufficient information to broadly model behaviour? If so, an individual profile could be collected using a few questions, taking only minutes. To the best of our knowledge, no such questionnaire exists.

### 1.1 The current study

The current study asked three main questions. **First**, do questionnaires reflect their purported cognitive functions? **Second**, is it the case that a visual imagery or autobiographical memory process is actually what is being assessed in questionnaires despite their purporting to be related to distinct cognitive functions? **Third**, how does a single question probing a cognitive function perform relative to a whole questionnaire? To address these questions we collected questionnaire and naturalistic task performance measures of imagination, autobiographical memory, future thinking and navigation from a large sample of participants (n = 217). The questionnaires were well established and provided detailed information about each cognitive domain of interest. We also included a bespoke, short, One Sentence Questionnaire designed for the current study. This questionnaire contained just 15 questions covering imagination, autobiographical memory, future thinking and navigation. For the task performance measures we used established, published tasks designed to assess each cognitive function in a naturalistic manner.

We considered the four cognitive functions in turn – imagination, autobiographical memory, future thinking, and navigation. For each, we started with the questionnaires most directly relevant to a cognitive function (i.e. the questionnaires on imagery for the imagination task; the questionnaires on memory for the autobiographical memory task, and so on) and examined how they related to task performance. We then investigated whether questionnaires not associated with a task (e.g. the questionnaires on imagery for the autobiographical memory task) were correlated with task performance. This allowed us to investigate whether an imagery or autobiographical memory process might be what is being tapped by the questionnaires rather than them being specific to a particular cognitive function. Finally, we assessed the ability of the questions of the One Sentence Questionnaire to predict performance on each task and how this compared to the established questionnaires, as before looking first at the questions most directly relevant, and then at those supposedly not associated with a task.

Considering our first question, we expected questionnaires to be related to their purported cognitive functions that would be evidenced by significant correlations between questionnaire and task performance data. For the second question, in our previous study (Clark et al., in press) we reported that the construction of scene imagery explained the relationships between the performance data from tasks assessing imagination, autobiographical memory, and future thinking, with navigation being partly related. Consequently, we hypothesised that the imagery questionnaires would be significantly correlated with imagination, autobiographical memory and future thinking performance, and perhaps partly with navigation performance. Furthermore, we hypothesised that the imagery questionnaires might even correlate with the task performance data to a greater extent than the questionnaires that actually purport to closely model a particular cognitive function. In relation to the novel One Sentence Questionnaire, as this was an exploratory aspect of the experiment, we had an open mind about whether or not it would reflect the task performance data as effectively as the longer questionnaires.

## 2. Methods

### 2.1. Participants

Two hundred and seventeen people took part in the study. They were aged between 20 and 41 years, had English as their first language and reported no psychological, psychiatric, neurological or behavioural health conditions. The age range was restricted to 20-41 to limit the possible effects of ageing. The mean age of the sample was 29.0 years old (95% CI; 20, 38) and included 109 females and 108 males. Participants were reimbursed £10 per hour for taking part which was paid at study completion. All participants gave written informed consent and the study was approved by the University College London Research Ethics Committee.

The guidelines described by Cohen (1992) were used to establish sample size during study design. A sample of 216 participants was determined to be robust to employing different statistical approaches when answering multiple questions of interest. For example, it allowed for sufficient power to identify medium effect sizes of correlations at alpha levels of 0.01, medium effect sizes when comparing two correlations at alpha levels of 0.05 and medium effect sizes with eight variables in multiple regressions at alpha levels of 0.01 – the statistical tests used in the current paper. In addition, although not relevant here, the sample size was large enough to identify medium effect sizes across multiple groups using ANOVAs at alpha levels of 0.01 and to conduct mediation analyses and structural equation modelling (Anderson & Gerbing, 1988) as used in Clark et al. (in press). A final sample of 217 was obtained due to over recruitment.

### 2.2. Procedure

Participants completed the questionnaires before any of the cognitive tasks. The tasks were conducted over multiple visits. The order of the tasks within each visit was the same for all participants (see Clark et al., in press). Task order was arranged so as to avoid task interference, for example, not having a verbal test followed by another verbal test, and to provide sessions of approximately equal length (∼3-3.5 hours, including breaks). The scene construction, future thinking and autobiographical memory tasks were completed during the same visit, while the navigation tasks were performed on a separate visit. All participants completed all parts of the study.

### 2.3. Questionnaires

Participants filled out a range of well-established, published questionnaires that purported to relate to one or more of the cognitive functions of interest in the current study. We briefly describe each questionnaire in alphabetical order and, where relevant, highlight particular subscales of interest.

#### 2.3.1. Memory Experiences Questionnaire

(MEQ; Sutin & Robins, 2007). This assesses the phenomenology of autobiographical memory across different dimensions. The original questionnaire asks participants to focus on a specific past event. For our purposes, this was adapted to concern the recall of autobiographical memories in general.

The full questionnaire examines ten dimensions (Accessibility, Coherence, Distancing, Emotional Intensity, Sharing, Sensory (non-visual) details, Time Perspective, Valence, Visual Perspective and Vividness). Here, we focussed the analyses on the Accessibility, Coherence, Sharing and Vividness subscales as they were most relevant for the current study. These subscales consist of varying numbers of statements which participants rate on a 5 point scale from 1 (strongly disagree) to 5 (strongly agree). Scores are calculated for each subscale by totalling the responses to each statement within the subscale. Questions from all subscales are intermixed throughout the questionnaire.

##### MEQ Accessibility

This subscale consists of five statements. Two are positively scored (“Memories are easy for me to recall”) and three are reverse scored (“It is difficult for me to think of past events”). The total score is out of 25.

##### MEQ Coherence

This subscale comprises eight statements. Four are positively scored (“I recognize the setting in which my memories take place”) and four are reverse scored (“I have a difficult time remembering events in a coherent manner”). The total score is out of 40.

##### MEQ Sharing

This subscale consists of six statements. Three are positively scored (“I frequently think about or talk about past event with others”) and three are reverse scored (“I rarely tell others about my memories”). The total score is out of 30.

##### MEQ Vividness

This subscale consists of six statements. Three items are positively scored (“My memory for events is very vivid”), and three are reverse scored (“My memory for events is dim”). The total score is out of 30.

#### 2.3.2. Object-Spatial Imagery and Verbal Questionnaire

(OSIVQ; Blazhenkova & Kozhevnikov, 2009). This questionnaire is designed to distinguish between different types of imagery users and has three subscales, two related to imagery and one to verbal processing. The Object subscale measures the ability to imagine vivid and detailed images of objects and scenes. The Spatial subscale measures the ability to process locations, movement and transformations, often represented by more technical and schematic imagery. The Verbal subscale measures the use of verbal strategies.

Here, for complete clarity, we renamed the Object subscale “Object-Scene” in order to better represent what the scale is designed to measure. Object imagery typically suggests an image of an object devoid of any background. However, as stated by Blazenkova and Kozhevnikov (2009), the Object subscale is not limited to individual objects but can also refer to imagery of patterns and scenes, characterising their colour, vividness, shape and details.

Each of the three subscales of the OSIVQ contains 15 statements. The participant is asked to rate each statement on a five point scale from 1 (totally disagree) to 5 (totally agree). The final score on each subscale is the average of the responses over the 15 items.

##### OSIVQ Object-Scene

For this subscale, statements include: “When reading fiction, I usually form a clear and detailed mental picture of a scene or room that has been described” and “I can close my eyes and easily picture a scene that I have experienced”. No statements are reverse scored.

##### OSIVQ Spatial

For this subscale, statements include: “My images are more like schematic representations for things and events rather than detailed pictures” and “I can easily sketch a blueprint for a building that I am familiar with”. One statement in the subscale is reverse scored (“I find it difficult to imagine how a three-dimensional geometric figure would exactly look like when rotated”).

##### OSVIQ Verbal

For this subscale, statements include: “When remembering a scene, I use verbal description rather than mental pictures” and “I am always aware of sentence structure”. Three statements in the subscale are reverse scored (e.g. “I have difficulty expressing myself in writing”).

#### 2.3.3. Plymouth Sensory Imagery Questionnaire, Appearance subscale

(PSIQ; Andrade et al., 2014). The PSIQ is a modernised version of the commonly used Vividness of Visual Imagery Questionnaire (VVIQ; Marks, 1973, 1995), with high correlations between the two questionnaires (Total score on the PISQ with VVIQ, r = 0.66, p < 0.001; PSIQ Appearance subscale with the VVIQ, r = 0.51, p < 0.001; Andrade et al., 2014). The PSIQ has an advantage over the VVIQ in also measuring imagery across multiple sensory modalities. Here, however, we focus solely on the Appearance subscale as this pertains specifically to the vividness of visual imagery – our area of interest. The subscale requires participants to imagine three scenarios: a bonfire, a sunset and a cat climbing a tree. They then rate the visual image they generated on an 11 point scale from 0 (no image at all) to 10 (vivid as real life). Scores on the three scenarios are summed to create a total score out of 30.

#### 2.3.4. Santa Barbara Sense of Direction Scale

(Hegarty et al., 2002). This questionnaire assesses spatial and navigational abilities, preferences and experiences. Fifteen statements are presented, with participants indicating their level of agreement with each statement. Ratings are made on a 7 point scale from 1 (strongly agree) to 7 (strongly disagree). Seven statements are positively coded (“I am very good at giving directions”) and eight are reverse scored (“I don’t have a very good “mental map” of my environment”). Scores are summed across the 15 statements and are then reversed to create a total score out of 105 (so that a high score reflects good navigation ability).

#### 2.3.5. Spontaneous Use of Imagery Scale

(SUIS; Reisberg et al., 2003). This questionnaire, consisting of 12 statements, measures how frequently an individual uses visual imagery. Participants read each statement and indicate the degree to which the statement is appropriate to them. Each statement is rated on a 5 point scale from 1 (never) to 5 (always). Example statements include: “If I am looking for new furniture in a store, I always visualize what the furniture would look like in particular places in my home” and “When I hear a radio announcer or DJ I’ve never actually seen, I usually find myself picturing what they might look like.” Scores are summed across the 12 items to give a final score out of 60.

#### 2.3.6. Subjective Memory Questionnaire

(SMQ; Bennett-Levy & Powell, 1980). This questionnaire probes memory for things people often try to remember. It is split into two sections. First, is a list of 36 items (“telephone numbers”; “jokes”; “birthdays”) which participants rate in response to the question “How good is your memory for…?” Answers are given on a 5 point scale from 1 (very bad) to 5 (very good). The second section asks the question “How often do you…?” in relation to seven experiences (e.g. “Set off to do something, then can’t remember what”; “Forget whether or not you have locked up the house”). Answers are provided on a 5 point scale from 1 (very rarely) to 5 (often). The total score is the sum of all responses (out of 215).

#### 2.3.7. Survey of Autobiographical Memory

(SAM; Palombo et al., 2013). This questionnaire assesses episodic and semantic aspects of autobiographical memory, as well as future thinking and spatial memory. There are 26 items in total. Participants respond on a five point scale regarding the extent to which they agree with each statement, from 1 (completely disagree) to 5 (completely agree). Scoring is determined via a weighting system. Responses are weighted and calculated together to provide an average score that centres around 100, like an IQ. Full details of the weighting and scoring procedure are available from the SAM authors.

##### SAM Episodic

This subscale assesses autobiographical memory recall. It contains 8 statements (“I am highly confident in my ability to remember past events”), two of which are reversed scored (“Specific events are difficult for me to recall”).

##### SAM Future

This subscale examines a participant’s ability to imagine future events. It contains 6 statements (“When I imagine an event in the future, the event generates vivid mental images that are specific in time and place”), one of which is reverse scored (“I have a difficult time imagining specific events in the future”).

##### SAM Spatial

This subscale assesses navigation ability. It contains 6 statements (“In general, my ability to navigate is better than most of my family/friends”), two of which are reverse scored (“I get lost easily, even in familiar areas”).

##### SAM Semantic

This subscale probes the ability to recall facts and information. It contains 6 statements (“I can learn and repeat facts easily, even if I don’t remember where I learned them”), two of which are reverse scored (“I have a hard time remembering information I have learned at school or work”).

#### 2.3.8. Visualizer –Verbalizer

(Kirby, Moore, & Schofield, 1988). This questionnaire assesses an individual’s preference for visual or verbal learning styles. Twenty statements are provided, 10 corresponding to visual items and 10 to verbal items. The participant indicates for each item whether, for them, the statement is true or false. Half of the statements are phrased positively, in that an answer of “true” reflects a visual or verbal learning preference (visual learning style: “The old saying ‘A picture is worth a thousand words’ is certainly true for me”; verbal learning style: “I have better than average fluency in using words”). The other half are phrased negatively where an answer of “false” reflects a visual or verbal learning preference (visual learning style: “I seldom use diagrams to explain things”; verbal learning style: “I dislike word games like crossword puzzles”). The two scales are scored separately. The final score (out of 10 for each scale) is the number of responses reflecting the stated learning style.

### 2.4. Questionnaire groups

We divided the established questionnaires into five groups, four of which related to our areas of interest: imagination, autobiographical memory, future thinking and navigation. The fifth group was composed of questionnaires relating to verbal and semantic processing. This was included as a control to ensure that questionnaires (regardless of subject matter) were not associated with the tasks for spurious reasons. The questionnaires in each group are shown in Table 1.

**Table 1.** The questionnaires in each group.

| Questionnaire Group | Questionnaires |
| --- | --- |
| Imagination | OSIVQ; Object-Scene<br>OSIVQ; Spatial<br>PSIQ; Appearance<br>SUIS<br>Visualizer |
| Autobiographical Memory | Memory Experiences Questionnaire (all subscales)<br>Subjective Memory Questionnaire<br>SAM; Episodic |
| Future Thinking | SAM; Future |
| Navigation | SAM; Spatial<br>Santa Barbara Sense of Direction Scale |
| Control | OSIVQ; Verbal<br>SAM; Semantic<br>Verbalizer |

Note. OSIVQ = Object-Spatial Imagery Verbal Questionnaire; PSIQ = Plymouth Sensory Imagery Questionnaire; SUIS = Spontaneous Use of Imagery Scale; SAM = Survey of Autobiographical Memory.

### 2.5. Experimental ‘One Sentence Questionnaire’

This questionnaire is new, and in developing and testing it we aimed to gain a broad profile of an individual in a short time frame. It consists of 15 questions covering the four areas of cognition that were of interest in this study – imagination, autobiographical memory, future thinking and navigation (see the Supplementary Materials for the questionnaire).

#### Imagery Ability

“Please rate your ability to construct a mental image”. Answers are on a 7 point scale from 1 (very high) to 7 (very low). This is reverse scored so that a high ability is a high score.

#### Imagery Use

“In everyday life, how much do you think in images (e.g. thinking in pictures in your mind)?” Answers are on a 7 point scale from 1 (not at all) to 7 (all the time).

#### Imagery as a Scene

“If you think in images, to what extent does this involve spatially coherent scenes (e.g. scenes that you could step into or operate within) compared to single objects?” Answers are on a 7 point scale from 1 (single objects) to 7 (coherent scenes).

#### Memory Ability

“Please rate your ability to remember your personal past”. Answers are on a 7 point scale from 1 (very high) to 7 (very low). This is reverse scored so that a high ability is a high score.

#### Memory in Imagery

“When recalling the past, to what extent do you think in images?” Answers are on a 7 point scale from 1 (not at all) to 7 (all the time).

#### Memory in Scene Imagery

“If you think in images when recalling the past, to what extent do you evoke spatially coherent scenes in your mind’s eye, compared to imagining single objects?” Answers are on a 7 point scale from 1 (single objects) to 7 (coherent scenes).

#### Memory in Words

“When recalling the past, how much do you think verbally (e.g. thinking in words and sentences)?” Answers are on a 7 point scale from 1 (not at all) to 7 (all the time).

#### Future Thinking Ability

“Please rate your ability to imagine future events”. Answers are on a 7 point scale from 1 (very high) to 7 (very low). This is reverse scored so that a high ability is a high score.

#### Future Thinking in Imagery

“When imagining the future, to what extent do you think in images?” Answers are on a 7 point scale from 1 (not at all) to 7 (all the time).

#### Future Thinking in Scene Imagery

**“**If you think in images when imagining the future, to what extent do you evoke spatially coherent scenes in your mind’s eye, compared to imagining single objects?” Answers are on a 7 point scale from 1 (single objects) to 7 (coherent scenes).

#### Future Thinking in Words

“When imagining the future, how much do you think verbally (e.g. thinking in words and sentences)?” Answers are on a 7 point scale from 1 (not at all) to 7 (all the time).

#### Navigation Ability

“Please rate your navigational ability”. Answers are on a 7 point scale from 1 (very high) to 7 (very low). This is reverse scored so that a high ability is a high score.

#### Navigation in Imagery

“When you navigate, to what extent do you think in images?” Answers are on a 7 point scale from 1 (not at all) to 7 (all the time).

#### Navigation in Scene Imagery

“If you think in images when navigating, to what extent do you evoke spatially coherent scenes in your mind’s eye, compared to imagining single objects?” Answers are on a 7 point scale from 1 (single objects) to 7 (coherent scenes).

#### Navigation in Words

“When navigating, how much do you think verbally (e.g. thinking in words and sentences)?” Answers are on a 7 point scale from 1 (not at all) to 7 (all the time).

### 2.6. Cognitive tasks

#### 2.6.1. Imagination – the scene construction test

(Hassabis et al., 2007). This test measures a participant’s ability to mentally construct an atemporal visual scene, meaning that the scene is not grounded in the past or the future. Participants construct seven different scenes of commonplace settings. For each scene, a short cue is provided (“Imagine lying on a beach in a beautiful tropical bay”), which the participant is asked to imagine and then describe out loud in as much detail as possible. All descriptions are audio recorded and transcribed for scoring. Participants are explicitly told not to describe a memory, but to create a new atemporal scene that they have never experienced before.

The main overall outcome measure is an “experiential index” which is calculated for each scene and then averaged. In brief, it is composed of four elements: numerical scoring of the content, participant ratings of their sense of presence (how much they felt like they were really there) and perceived vividness, participant ratings of the spatial coherence of the scene, and an experimenter rating of overall quality of the scene (see Supplementary Materials for full details). Double scoring was performed on 20% of the data. We took the most stringent approach to identifying across-experimenter agreement. Inter-class correlation coefficients, with a two-way random effects model looking for absolute agreement, indicated excellent agreement among the experimenter ratings (minimum score of 0.90; see Supplementary Materials, Table S1). For reference, a score of 0.8 or above is considered excellent agreement beyond chance.

#### 2.6.2. Autobiographical memory – the Autobiographical Interview

(AI; Levine et al., 2002). The AI asks participants to recall and describe autobiographical memories from a specific time and place over four time periods – early childhood (up to age 11), teenage years (from 11-17 years of age), adulthood (from aged 18 up to 12 months prior to the interview; two memories were requested) and the last year (a memory from the last 12 months).

Memories provided from the AI are scored to collect “internal” and “external” details of the event (see Supplementary Materials for details). Internal details are those describing the event in question (i.e., episodic details). External details describe semantic information concerning the event, or non-event information. The main outcome measure is the number of internal details as a percentage of the total number of utterances (i.e., combined internal and external details). This provides a measure of episodic memory, while taking into account a participant’s verbosity. Overall AI scores were obtained by averaging performance across the five autobiographical memories. Double scoring found excellent agreement across the experimenters (minimum score of 0.81; see Supplementary Materials, Table S2).

As a secondary measure, we also examined the vividness of the memories generated. This was a participant rating performed for each memory. Participants answered the question “How clearly can you visualize this event?” on a 6 point scale from 1 (vague memory, no recollection) to 6 (extremely clear as if it’s happening now). Responses were averaged across the five memories.

#### 2.6.3. Future thinking

(Hassabis et al., 2007). The future thinking task follows the same procedure as the scene construction task, but requires participants to imagine three plausible future scenes involving themselves (an event at the weekend; next Christmas; the next time they meet a friend). There are two main differences between the future thinking task and the scene construction task. First, unlike scenes in the scene construction task, scenes in the future thinking task involve ‘mental time travel’ to the future, so they have a clear temporal dimension. Second, the cues for the scene construction tasks are somewhat more specific than those employed in the future thinking task (see Hurley, Maguire, & Vargha-Khadem, 2011 for more on this issue). Scoring procedures are the same as for the scene construction task. Double scoring identified excellent agreement across the experimenters (minimum score of 0.88; see Supplementary Materials, Table S3).

#### 2.6.4. Navigation

(Woollett & Maguire, 2010). In this test, navigation ability is assessed using movies of navigation through an unfamiliar town. Movie clips of two overlapping routes through this real town (Blackrock, in Dublin, Ireland) are shown to a participant four times.

Five tasks are used to assess navigational ability. First, following each viewing of the route movies, participants are shown four short clips – two from the actual routes and two distractors. Participants indicate whether they had seen that clip or not. Second, after all four route viewings are completed, recognition memory for landmarks is tested. A third test involves assessing knowledge of the spatial relationships between landmarks in the town. Fourth, route knowledge is examined by having participants place sets of photographs from the routes in the correct order as if travelling through the town. Finally, participants draw a sketch map of the two routes including as many landmarks as they can remember. Sketch maps are scored in terms of the number of road segments, road junctions, correct landmarks, landmark positions, the orientation of the routes and an overall map quality score from the experimenters (see Supplementary Materials for full details). Double scoring was performed on 20% of the sketch maps finding excellent agreement (minimum of 0.89; see Supplementary Materials, Table S4). An overall navigation score was calculated by combining the scores from the five tasks.

### 2.7. Statistical analyses

Questionnaire and task data were first summarised using means and 95% confidence intervals, calculated in SPSS v22. Bivariate Person correlations were performed between the questionnaires and tasks in SPSS v22. Correlations were compared using the technique described by Meng, Rosenthal and Rubin (1992), which extends the Fisher z transformation, allowing for more accurate testing and comparison of two related correlations. The analysis was performed using the R cocor package, v1.1.3 (Diedenhofen & Musch, 2015) in R v3.4 (R Core Team, 2017). Multiple regressions were performed in SPSS v22. Effect sizes are reported as R^2^ values. There were no missing data in any of the analyses.

As we performed 32 correlations for each cognitive task (17 correlations pertaining to the established questionnaires and 15 correlations for the exploratory One Sentence Questionnaire), the alpha level for the initial correlations was set at p < 0.001 to avoid false positives. We note that this is slightly more stringent than using Bonferroni correction which would suggest p < 0.0016, but we felt it was prudent to adopt a more cautious approach given our reasonably large sample size. For the analyses following the correlations (i.e. the comparisons of the correlations and the multiple regressions) we only included those variables already determined to be significantly correlated at p < 0.001. Therefore, when comparing the correlations and identifying significant variables in the regressions, as we had already been selective, the alpha level was set at p < 0.05.

## 3. Results

Performance data from the cognitive tasks have been presented in a previous paper that addressed a completely different research question to that under consideration here (Clark et al., in press). As detailed in the introduction, the previous research question involved assessing the relationships between imagination, autobiographical memory, future thinking and navigation in terms of task performance. By contrast, in the current study we investigated how questionnaire data were related to task performance on each of the aforementioned tasks. None of the questionnaire data has been reported previously. A summary of the outcome measures for the questionnaires and cognitive tasks is presented in Table 2. A wide range of scores was obtained for all variables.

**Table 2.** Means and 95% confidence internals (CI) for the questionnaires and cognitive tasks.

|  | Mean | Lower<br>CI | Upper<br>CI |
| --- | --- | --- | --- |
| **Questionnaires** |  |  |  |
| Memory Experience Questionnaire; Accessibility (/25) | 17.29 | 11.0 | 22.0 |
| Memory Experience Questionnaire; Coherence (/40) | 28.87 | 22.0 | 36.0 |
| Memory Experience Questionnaire; Sharing (/30) | 21.18 | 13.0 | 29.0 |
| Memory Experience Questionnaire; Vividness (/30) | 21.06 | 13.0 | 27.10 |
| Object Spatial Imagery and Verbal Questionnaire; Object-Scene (/5) | 3.20 | 2.13 | 4.13 |
| Object Spatial Imagery and Verbal Questionnaire; Spatial (/5) | 2.87 | 1.80 | 4.0 |
| Object Spatial Imagery and Verbal Questionnaire; Verbal (/5) | 3.11 | 2.13 | 3.87 |
| Plymouth Sensory Imagery Questionnaire; Appearance (/30) | 21.89 | 10.0 | 30.0 |
| Santa Barbara Sense of Direction Scale (/105) | 55.17 | 29.0 | 79.0 |
| Spontaneous Use of Imagery Scale (/60) | 39.64 | 24.90 | 51.0 |
| Subjective Memory Questionnaire (/215) | 146.88 | 117.90 | 173.20 |
| Survey of Autobiographical Memory; Episodic | 98.63 | 79.18 | 123.60 |
| Survey of Autobiographical Memory; Future | 90.70 | 78.86 | 120.84 |
| Survey of Autobiographical Memory; Spatial | 97.81 | 69.37 | 119.88 |
| Survey of Autobiographical Memory; Semantic | 95.76 | 79.05 | 116.98 |
| Visualizer (/10) | 8.12 | 5.0 | 10.0 |
| Verbalizer (/10) | 6.94 | 3.0 | 10.0 |
| One Sentence: Imagery Ability (/7) | 5.27 | 3.0 | 7.0 |
| One Sentence: Imagery Use (/7) | 4.69 | 2.0 | 7.0 |
| One Sentence: Imagery as a Scene (/7) | 4.67 | 2.0 | 7.0 |
| One Sentence: Memory Ability (/7) | 4.83 | 3.0 | 7.0 |
| One Sentence: Memory in Imagery (/7) | 5.51 | 3.0 | 7.0 |
| One Sentence: Memory in Scene Imagery (/7) | 5.43 | 3.0 | 7.0 |
| One Sentence: Memory in Words (/7) | 3.84 | 2.0 | 6.0 |
| One Sentence: Future Thinking Ability (/7) | 5.53 | 4.0 | 7.0 |
| One Sentence: Future Thinking in Imagery (/7) | 5.14 | 2.0 | 7.0 |
| One Sentence: Future Thinking in Scene Imagery (/7) | 5.13 | 2.0 | 7.0 |
| One Sentence: Future Thinking in Words (/7) | 4.38 | 2.0 | 6.0 |
| One Sentence: Navigation Ability (/7) | 5.04 | 3.0 | 7.0 |
| One Sentence: Navigation in Imagery (/7) | 5.19 | 2.90 | 7.0 |
| One Sentence: Navigation in Scene Imagery (/7) | 4.98 | 2.0 | 7.0 |
| One Sentence: Navigation in Words (/7) | 3.89 | 1.90 | 6.0 |
| **Tasks** |  |  |  |
| Scene Construction Experiential Index (/60) | 40.50 | 29.50 | 50.13 |
| Autobiographical Interview Number of Internal Details | 23.95 | 13.80 | 37.41 |
| Autobiographical Interview Vividness (/6) | 4.62 | 3.38 | 5.80 |
| Future Thinking Experiential Index (/60) | 39.12 | 25.00 | 49.99 |
| Navigation (/250) | 143.46 | 88.90 | 201.50 |

The results are reported in the following manner:

**(1)** For each cognitive function of interest, the scores on the associated cognitive task were correlated with the scores from the questionnaires in the relevant group (see Table 1), i.e. those that purport to be closely related to this cognitive function.
**(2)** For those results that were significant (at p < 0.001) at the first step, we next examined whether there were any differences in correlations between the questionnaires, to see which questionnaire(s) best represented the cognitive task in question.
**(3)** We then conducted a multiple regression analysis to examine whether the questionnaires tapped into the same cognitive construct, or if the different questionnaires were representing distinct aspects of the cognitive function.
**(4)** We next sought to ascertain if questionnaires which were related to other cognitive functions (i.e. the other questionnaire groups in Table 1), were associated with performance on the cognitive task in question (at p < 0.001).
**(5)** The next step was to investigate whether the questionnaires associated with the cognitive task outperformed those from the other questionnaire groups, or whether all the significantly correlated questionnaires correlated with the cognitive task to a similar extent.
**(6)** We then performed multiple regression to see if the questionnaires were correlating with the cognitive task via the same cognitive construct, or if there were multiple distinct contributions from the different types of questionnaires.
**(7)** We next moved to our exploratory One Sentence Questionnaire, to test whether the relevant questions from this questionnaire correlated with task performance (e.g. did the imagination questions correlate with scene construction performance).
**(8)** We then compared the correlations between the relevant questions of the One Sentence Questionnaire with the significantly correlating established questionnaires.
**(9)** Our final analyses then examined whether the questions from the One Sentence Questionnaire that were relevant to other cognitive functions were associated with performance on the task in question, and how this compared to the relevant questions of the One Sentence Questionnaire.
**(10)** Finally, we relate the questionnaire results for a cognitive function back to the three research questions.

Of note, some of the steps above were not performed for a cognitive function if the main correlations between questionnaires and the cognitive task (steps 1, 4, 7) were not significant at p < 0.001.

### 3.1 Imagination

#### 3.1.1. Questionnaires from the imagery group

We first assessed whether or not the imagery questionnaires were correlated with performance on a test of imagination - the scene construction test. As is evident in Table 3, this was predominantly the case. Of the established questionnaires, the PSIQ, OSIVQ Object-Scene subscale and SUIS all correlated with performance on the scene construction test. On the other hand, the Visualizer questionnaire and the OSIVQ Spatial subscale were not correlated with scene construction, suggesting a possible divide in the imagery type being measured.

**Table 3.**
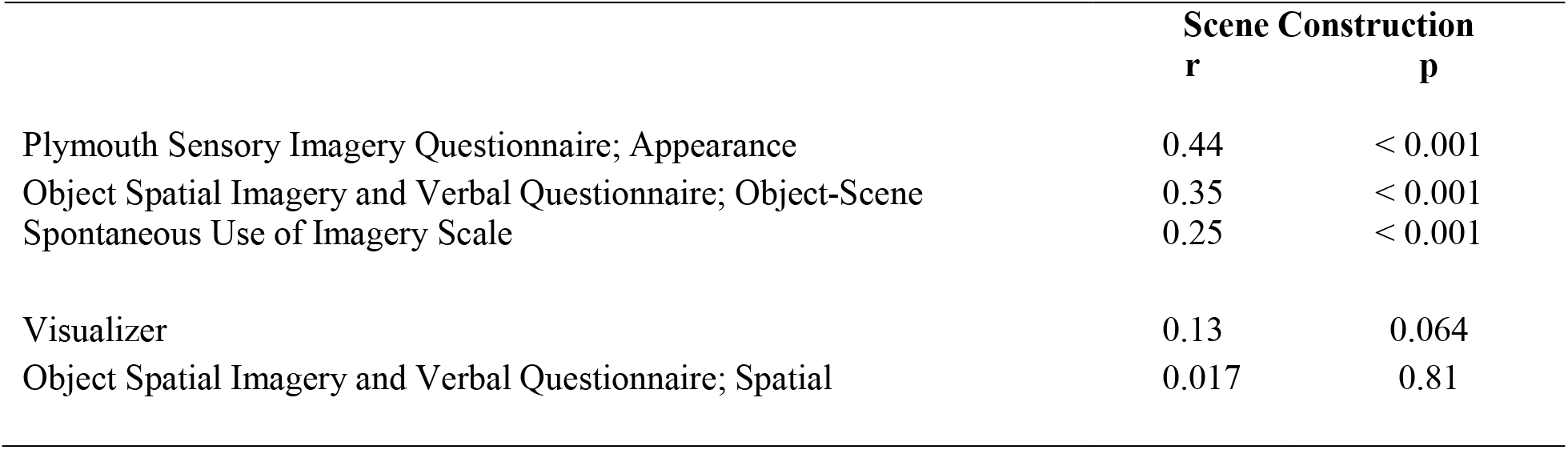
Correlation coefficients of the imagery questionnaires with scene construction performance.

Next, we wanted to know if there were any differences in correlations between the questionnaires. To investigate this, we compared the correlation coefficients of the questionnaires that were significantly associated with scene construction, and the results are reported in Table 4. The only significant difference was that the PSIQ had a greater correlation with scene construction performance than the SUIS, while the OSIVQ Object-Scene subscale had a correlational value in between the PSIQ and SUIS, but was not significantly different to either.

**Table 4.**
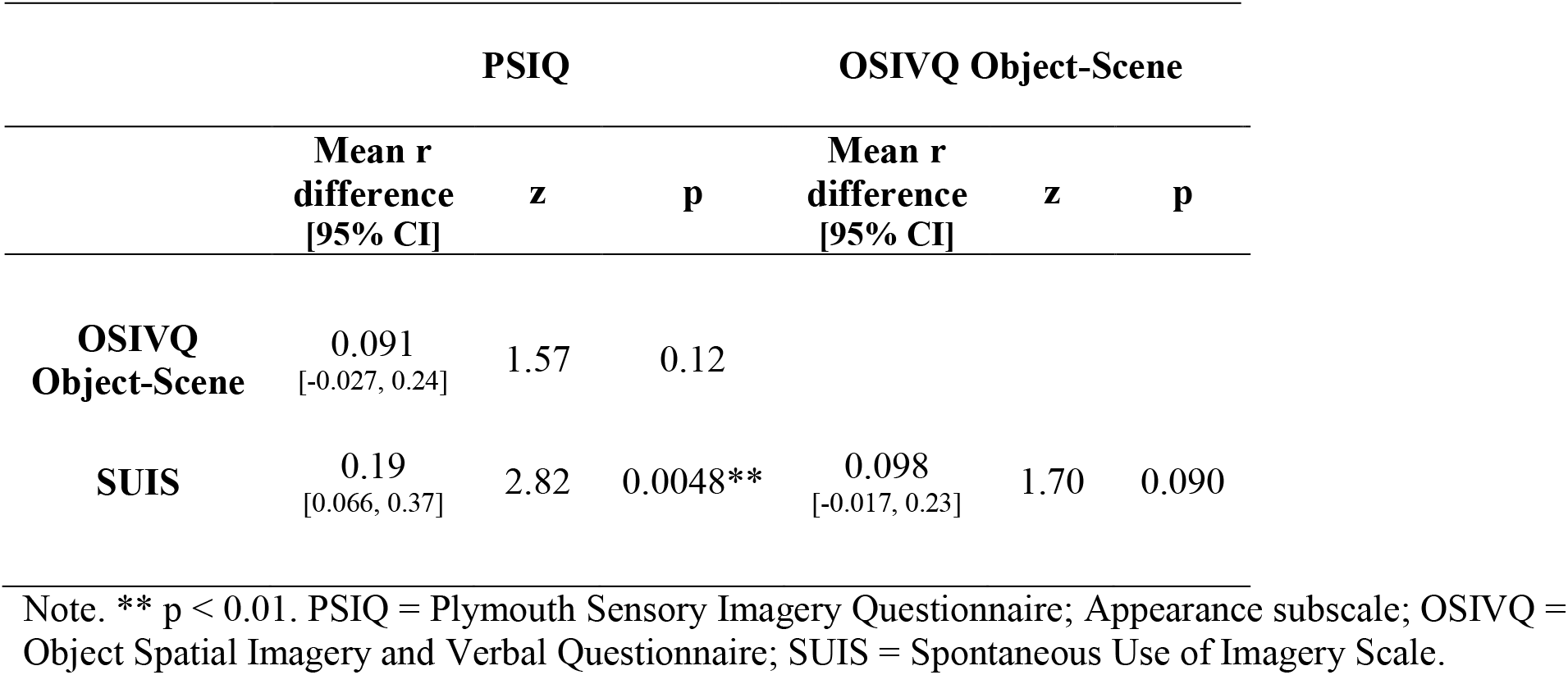
Comparisons between the significant correlation coefficients of the imagery questionnaires with scene construction performance. The difference in correlation is the column questionnaires less the row questionnaires.

Next, we wanted to examine whether the questionnaires tapped into the same cognitive construct, or if the different questionnaires are representing distinct aspects of imagery. To do this, we performed a multiple regression analysis using the imagery questionnaires that were significantly correlated with scene construction performance (at the p < 0.001 level). The results are shown in Table 5. Only the PSIQ was associated with scene construction performance (regression model statistics: F(3,213 = 18.62, p < 0.001, R^2^ = 0.21, Adj. R^2^ = 0.20).

**Table 5.** Multiple regression of the imagery questionnaires with scene construction performance.

|  | **Beta**<br>**(95% CI)** | **Standardised**<br>**Beta** | **t** | **p** |
| --- | --- | --- | --- | --- |
| PSIQ | 0.38 (0.22, 0.54) | 0.35 | 4.77 | < 0.001 |
| OSIVQ Object-Scene | 1.45 (-0.18, 3.09) | 0.15 | 1.76 | 0.081 |
| SUIS | 0.007 (-0.11, 0.13) | 0.009 | 0.11 | 0.91 |

Note. PSIQ = Plymouth Sensory Imagery Questionnaire; Appearance subscale; OSIVQ = Object Spatial Imagery and Verbal Questionnaire; SUIS = Spontaneous Use of Imagery Scale.

Why might the PSIQ be key? The PSIQ is the questionnaire most similar to the scene construction test. The PSIQ asks participants to rate how vividly they can imagine three specific scenes (a bonfire, a sunset, a cat climbing a tree). The scene construction test asks participants to imagine and describe specific scenes (e.g. a busy fishing harbour). It, therefore, follows that a participant’s subjective belief about their ability to perform this task correlated particularly well with their score on essentially the same task.

#### 3.1.2. Questionnaires from the other groups

We next sought to ascertain if questionnaires which are typically associated with the other cognitive functions (namely, autobiographical memory, future thinking and navigation), were associated with performance on the scene construction task. To check that questionnaires were not simply associated with task performance for spurious reasons (e.g. the product of a reasonably large sample size and performing multiple correlations), we also included questionnaires which examined less related cognitive constructs (verbal processing and semantic memory) to act as controls.

As can be seen in Table 6, the majority of questionnaires pertaining to autobiographical memory and future thinking were also significantly (p < 0.001) correlated with performance on the scene construction test. This is in direct contrast to the control questionnaires (assessing verbal and semantic processing) that were not correlated with scene construction performance. Of note, the significant correlations seemed to occur when the questionnaire also assessed visual imagery, for example, when asking about the vividness of memory recall (the MEQ Vividness subscale). The questionnaires assessing navigation, however, told a different story, with neither questionnaire being correlated with scene construction performance (at the p < 0.001 level). Navigation questionnaires may, therefore, be assessing a different cognitive construct than the imagery questionnaires.

**Table 6.** Correlation coefficients of the questionnaires from other groups with scene construction performance.

|  | Scene Construction |  |
| --- | --- | --- |
|  | r | p |
| **Memory Questionnaires** |  |  |
| Memory Experience Questionnaire; Vividness | 0.31 | < 0.001 |
| Memory Experience Questionnaire; Coherence | 0.28 | < 0.001 |
| Subjective Memory Questionnaire | 0.27 | < 0.001 |
| Survey of Autobiographical Memory; Episodic | 0.25 | < 0.001 |
| Memory Experience Questionnaire; Accessibility | 0.22 | 0.001 |
| Memory Experience Questionnaire; Sharing | 0.20 | 0.003 |
| **Future Thinking Questionnaire** |  |  |
| Survey of Autobiographical Memory; Future | 0.30 | < 0.001 |
| **Navigation Questionnaires** |  |  |
| Survey of Autobiographical Memory; Spatial | 0.19 | 0.004 |
| Santa Barbara Sense of Direction Scale | 0.17 | 0.011 |
| **Verbal and Semantic Memory Questionnaires** |  |  |
| Survey of Autobiographical Memory; Semantic | 0.12 | 0.082 |
| Verbalizer | 0.10 | 0.13 |
| Object Spatial Imagery and Verbal Questionnaire; Verbal | 0.089 | 0.19 |

The next step was to investigate whether the imagery questionnaires were outperforming the questionnaires from the other groups, or whether all the significantly correlated questionnaires were correlating with scene construction performance to the same extent. To examine this, we compared the correlations of the significantly associated questionnaires from the other groups with the significantly correlated imagery questionnaires.

As can be seen in Table 7, the PSIQ had a higher correlation with scene construction performance than questionnaires from the other groups. As such, directly asking about imagery led to a greater correlation than asking about indirectly-related cognitive constructs. On the other hand, the OSIVQ Object-Scene subscale and the SUIS were similarly correlated with scene construction as the indirectly-related questionnaires. This suggests that the memory and future thinking questionnaires, although designed to examine different cognitive constructs, may in fact have a substantial overlap with imagery.

**Table 7.** Comparisons between the correlation coefficients of the imagery questionnaires and the other group questionnaires significantly correlated with scene construction performance. The difference in correlation is the imagery questionnaires less the other group questionnaires (a negative value shows higher correlation for the row questionnaire).

|  | PSIQ |  |  | OSIVQ Object-Scene |  |  | SUIS |  |  |
| --- | --- | --- | --- | --- | --- | --- | --- | --- | --- |
|  | Mean r<br>difference<br>[95% CI] | z | p | Mean r<br>difference<br>[95% CI] | z | p | Mean r<br>difference<br>[95% CI] | z | p |
| **MEQ**<br>**Vividness** | 0.13<br>[0.001, 0.30] | 1.97 | 0.049* | 0.039<br>[-0.096, 0.18] | 0.61 | 0.54 | -0.059<br>[-0.23, 0.10] | -0.76 | 0.45 |
| **MEQ**<br>**Coherence** | 0.16<br>[0.012, 0.36] | 2.09 | 0.036* | 0.069<br>[-0.093, 0.25] | 0.88 | 0.38 | -0.029<br>[-0.22, 0.16] | -0.33 | 0.74 |
| **SMQ** | 0.17<br>[0.033, 0.36] | 2.35 | 0.019* | 0.082<br>[-0.049, 0.23] | 1.27 | 0.20 | -0.016<br>[-0.20, 0.16] | -0.19 | 0.85 |
| **SAM Episodic** | 0.19<br>[0.057, 0.38] | 2.65 | 0.0081** | 0.10<br>[-0.015, 0.24] | 1.73 | 0.084 | 0.004<br>[-0.16, 0.17] | 0.050 | 0.96 |
| **SAM Future** | 0.14<br>[0.0019, 0.33] | 1.98 | 0.047* | 0.05<br>[-0.083, 0.19] | 0.79 | 0.43 | 0.048<br>[-0.21, 0.11] | -0.65 | 0.52 |

Note. \* = $p < 0.05$ ; \*\* = $p < 0.01$ . PSIQ = Plymouth Sensory Imagery Questionnaire; Appearance subscale; OSIVQ = Object Spatial Imagery and Verbal Questionnaire; SUI = Spontaneous Use of Imagery Scale; MEQ = Memory Experience Questionnaire; SMQ = Subjective Memory Questionnaire; SAM = Survey of Autobiographical Memory.

Given the correlations of questionnaires from the other groups (in addition to the imagery questionnaires) with scene construction performance, we next performed a multiple regression to ascertain if the questionnaires were correlating with scene construction via the same construct, or if there were multiple distinct contributions from the different types of questionnaires.

The results of the multiple regression are presented in Table 8. Two questionnaires were found to contribute to scene construction performance – the PSIQ and the MEQ Coherence subscale [regression model statistics: F(8,208) = 8.57, p < 0.001, R^2^ = 0.25, Adj. R^2^ = 0.22], with the standardised Betas suggesting that the PSIQ had twice the weighting of the MEQ Coherence.

**Table 8.**
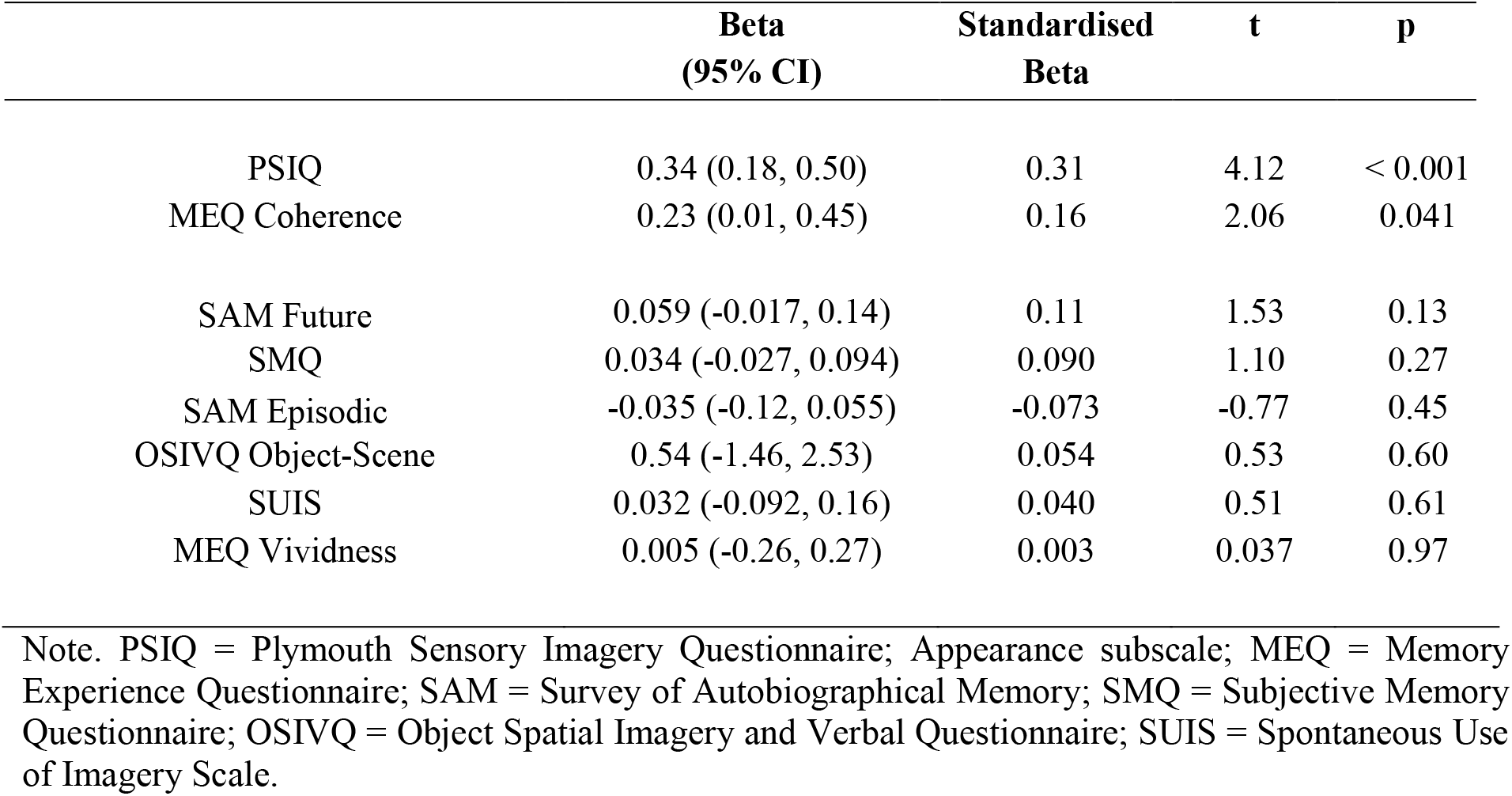
Multiple regression of all the questionnaires significantly correlated with scene construction performance.

|  | **Beta**<br>**(95% CI)** | **Standardised**<br>**Beta** | **t** | **p** |
| --- | --- | --- | --- | --- |
| PSIQ | 0.34 (0.18, 0.50) | 0.31 | 4.12 | < 0.001 |
| MEQ Coherence | 0.23 (0.01, 0.45) | 0.16 | 2.06 | 0.041 |
| SAM Future | 0.059 (-0.017, 0.14) | 0.11 | 1.53 | 0.13 |
| SMQ | 0.034 (-0.027, 0.094) | 0.090 | 1.10 | 0.27 |
| SAM Episodic | -0.035 (-0.12, 0.055) | -0.073 | -0.77 | 0.45 |
| OSIVQ Object-Scene | 0.54 (-1.46, 2.53) | 0.054 | 0.53 | 0.60 |
| SUIS | 0.032 (-0.092, 0.16) | 0.040 | 0.51 | 0.61 |
| MEQ Vividness | 0.005 (-0.26, 0.27) | 0.003 | 0.037 | 0.97 |

Note. PSIQ = Plymouth Sensory Imagery Questionnaire; Appearance subscale; MEQ = Memory Experience Questionnaire; SAM = Survey of Autobiographical Memory; SMQ = Subjective Memory Questionnaire; OSIVQ = Object Spatial Imagery and Verbal Questionnaire; SUIS = Spontaneous Use of Imagery Scale.

The multiple regression, therefore, suggests that the majority of correlations observed between the memory and future thinking questionnaires and scene construction ability could be explained by visual imagery. If the memory or future thinking questionnaires represented different constructs outside of imagery, then they would have been identified as separate variables by the multiple regression – as was the case with the MEQ Coherence subscale. So what does the MEQ Coherence subscale represent? This subscale probes how well a memory is placed together in a logical fashion. Hence it seems to add information to the PSIQ, which is concerned only with the vividness of an image. Overall, therefore, while many questionnaires probing imagery, memory and future thinking correlated with scene construction performance, all of these questionnaires seem to be, in some way, tapping into the same constructs as those measured by the PSIQ and MEQ Coherence.

#### 3.1.3. One Sentence Questionnaire

Our final area of interest in relation to scene construction performance was the exploratory One Sentence Questionnaire. First, we assessed whether or not the imagery questions from the One Sentence Questionnaire correlated with scene construction performance, finding that they all did (One Sentence Imagery Use: r = 0.32, p < 0.001; One Sentence Imagery as a Scene: r = 0.32, p < 0.001; One Sentence Imagery Ability: r = 0.31, p < 0.001).

We then tested how the One Sentence Questionnaire imagery questions performed in comparison to the established imagery questionnaires that also correlated with scene construction performance (the PSIQ, OSIVQ Object-Scene subscale and the SUIS). The results are shown in Table 9. The only significant differences were between the PSIQ and One Sentence: Imagery as a Scene and One Sentence: Imagery Ability. That is, the questions from the One Sentence Questionnaire were correlated with scene construction to a similar degree as most of the longer, and more time consuming, established questionnaires.

**Table 9.** Comparisons between the correlation coefficients of the established imagery questionnaires that correlated with scene construction performance and the One Sentence Questionnaire imagery questions. The difference in correlation is the established questionnaires less the One Sentence Questionnaire (a negative value shows higher correlation for the row questionnaire).

|  | PSIQ |  |  | OSIVQ Object-Scene |  |  | SUIS |  |  |
| --- | --- | --- | --- | --- | --- | --- | --- | --- | --- |
|  | Mean r<br>difference<br>[95% CI] | z | p | Mean r<br>difference<br>[95% CI] | z | p | Mean r<br>difference<br>[95% CI] | z | p |
| **Imagery Use** | 0.12<br>[-0.0078, 0.30] | 1.86 | 0.063 | 0.032<br>[-0.096, 0.17] | 0.53 | 0.59 | -0.066<br>[-0.21, 0.071] | -0.98 | 0.33 |
| **Imagery as a Scene** | 0.12<br>[0.0009, 0.28] | 1.97 | 0.049* | 0.029<br>[-0.11, 0.18] | 0.45 | 0.66 | -0.069<br>[-0.24, 0.085] | -0.92 | 0.36 |
| **Imagery Ability** | 0.13<br>[0.012, 0.29] | 2.13 | 0.033* | 0.037<br>[-0.087, 0.17] | 0.63 | 0.53 | -0.061<br>[-0.22, 0.089] | -0.83 | 0.40 |

Note. \* = $p < 0.05$ . PSIQ = Plymouth Sensory Imagery Questionnaire; Appearance subscale; OSIVQ = Object Spatial Imagery and Verbal Questionnaire; SUIS = Spontaneous Use of Imagery Scale.

Next, we sought to ascertain if, like the established questionnaires, the One Sentence Questionnaire questions designed to look at other cognitive functions (namely, autobiographical memory, future thinking and navigation), were also associated with performance on the scene construction task. The results can be seen in Table 10. Mirroring the established questionnaires, significant correlations seemed to occur when the questions also involved visual imagery.

**Table 10.** Correlation coefficients of the One Sentence Questionnaire questions from other groups with scene construction performance.

|  | **Scene Construction** |  |
| --- | --- | --- |
|  | **r** | **p** |
| **Memory Questions** |  |  |
| One Sentence: Memory in Imagery | 0.29 | < 0.001 |
| One Sentence: Memory in Scene Imagery | 0.26 | < 0.001 |
| One Sentence: Memory Ability | 0.17 | 0.013 |
| One Sentence: Memory in Words | -0.18 | 0.007 |
| **Future Thinking Questions** |  |  |
| One Sentence: Future Thinking in Imagery | 0.26 | < 0.001 |
| One Sentence: Future Thinking in Scene Imagery | 0.26 | < 0.001 |
| One Sentence: Future Thinking Ability | 0.22 | 0.001 |
| One Sentence: Future Thinking in Words | -0.030 | 0.67 |
| **Navigation Questions** |  |  |
| One Sentence: Navigation in Imagery | 0.24 | < 0.001 |
| One Sentence: Navigation in Scene Imagery | 0.20 | 0.003 |
| One Sentence: Navigation Ability | 0.17 | 0.011 |
| One Sentence: Navigation in Words | 0.067 | 0.33 |

Finally, we investigated whether the imagery questions from the One Sentence Questionnaire outperformed the One Sentence Questionnaire questions from the other groups, or whether all the significantly correlated questions were correlating with scene construction performance to the same extent. To examine this, we compared the correlations of the significantly associated One Sentence Questionnaire questions from the other groups with the One Sentence Questionnaire imagery questions.

As can be seen in Table 11, the imagery based One Sentence Questionnaire questions were similarly correlated with scene construction as the other significantly correlated One Sentence Questionnaire questions. This suggests two potential conclusions. First, it could be that the specificity of the One Sentence Questionnaire questions is poor. Or, second, and in line with the established questionnaires, it may be that, while the other questions are designed to examine different cognitive constructs, due to the inclusion of visual imagery in these questions, associations with scene construction performance remain.

**Table 11.** Comparisons between the correlation coefficients of imagery questions from the One Sentence Questionnaire and the other questions from the One Sentence Questionnaire that were significantly correlated with scene construction performance. The difference in correlation is the imagery questions less the other group questions.

|  | **One Sentence:**<br>**Imagery Everyday** |  |  | **One Sentence:**<br>**Imagery as a Scene** |  |  | **One Sentence:**<br>**Imagery Ability** |  |  |
| --- | --- | --- | --- | --- | --- | --- | --- | --- | --- |
|  | **Mean r**<br>**difference**<br>**[95% CI]** | **z** | **p** | **Mean r**<br>**difference**<br>**[95% CI]** | **z** | **p** | **Mean r**<br>**difference**<br>**[95% CI]** | **z** | **p** |
| **Memory in**<br>**Imagery** | 0.025<br>[-0.12, 0.17] | 0.37 | 0.71 | 0.028<br>[-0.11, 0.18] | 0.42 | 0.67 | 0.02<br>[-0.11, 0.16] | 0.32 | 0.75 |
| **Memory in**<br>**Scene Imagery** | 0.052<br>[-0.096, 0.21] | 0.73 | 0.46 | 0.055<br>[-0.076, 0.20] | 0.86 | 0.39 | 0.047<br>[-0.096, 0.20] | 0.68 | 0.49 |
| **Navigation in**<br>**Imagery** | 0.078<br>[-0.064, 0.23] | 1.11 | 0.26 | 0.081<br>[-0.084, 0.26] | 1.0 | 0.32 | 0.073<br>[-0.080, 0.24] | 0.97 | 0.33 |
| **Future in**<br>**Imagery** | 0.055<br>[-0.073, 0.19] | 0.88 | 0.38 | 0.058<br>[-0.059, 0.19] | 1.02 | 0.31 | 0.05<br>[-0.087, 0.20] | 0.76 | 0.45 |
| **Future in**<br>**Scene Imagery** | 0.055<br>[-0.083, 0.20] | 0.82 | 0.41 | 0.058<br>[-0.058, 0.18] | 1.03 | 0.30 | 0.05<br>[-0.091, 0.20] | 0.73 | 0.46 |

#### 3.1.4. Imagination summary

The main associations between the questionnaires and performance on the scene construction test can be seen in Figure 1.

**Figure 1.**
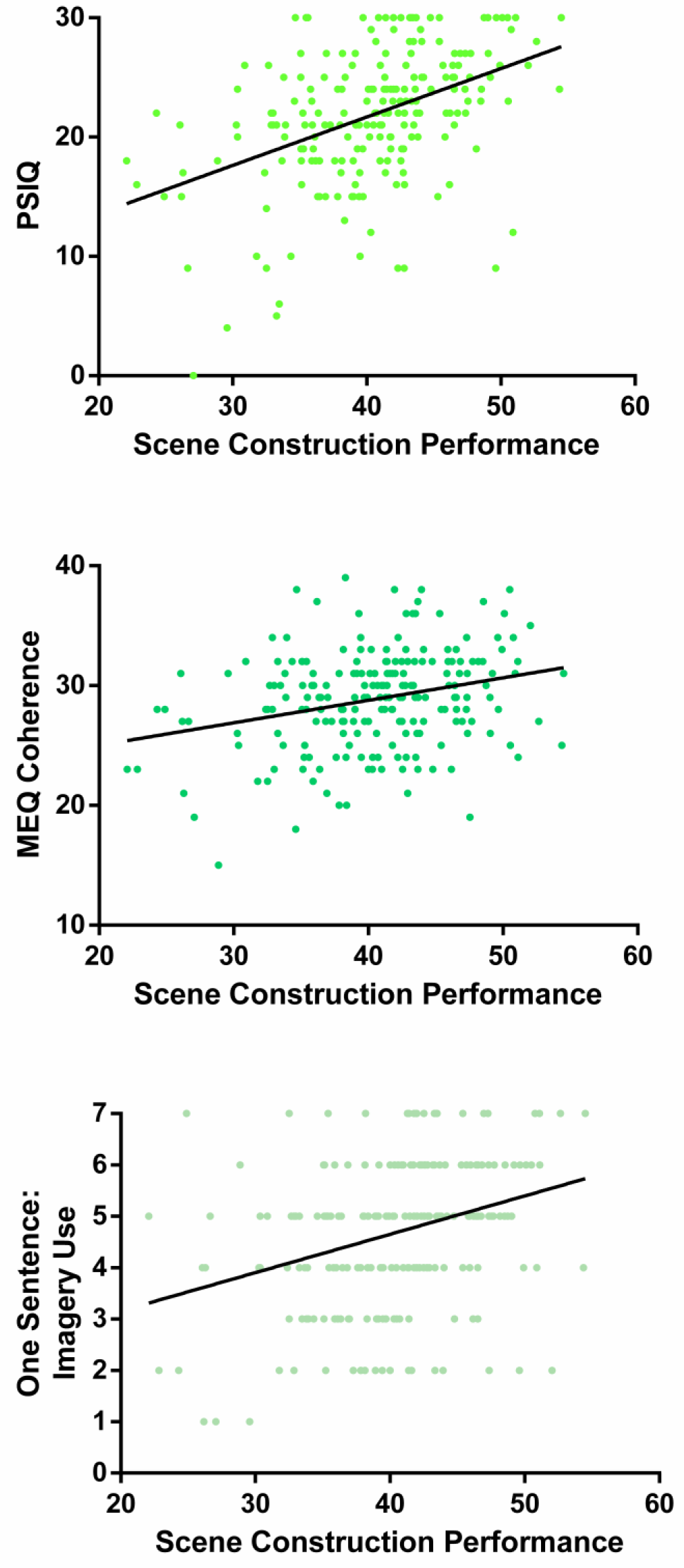
Correlations of the PSIQ, MEQ Coherence subscale and the One Sentence: Imagery Use question with performance on the scene construction task.

Considering our three research questions, first, imagery questionnaires that focus on concrete visual imagery measure the construct they are designed to reflect, namely imagery ability. Second, for the questionnaires from the other groups, overlap between imagination, autobiographical memory and future thinking questionnaires (but not navigation) was clearly evident with regard to scene construction performance. Notably, this relationship seemed to be based around the use of visual imagery. Third, the imagery questions from the One Sentence Questionnaire performed at the same level as the established, and more time consuming to complete, questionnaires. The only exception to this being the PSIQ, the questionnaire which also produced higher correlations with scene construction performance when compared to the established imagery questionnaires.

### 3.2. Autobiographical memory

#### 3.2.1. Questionnaires from the autobiographical memory group

Considering next the memory questionnaires, the task performance data here were the number of AI internal (episodic) details as a percentage of total utterances (internal and external details combined), to provide a measure of autobiographical memory ability with verbosity taken into account. The expectation based on the literature was that responses on the memory questionnaires would be correlated with AI internal details. However, as is shown in Table 12, none of the memory questionnaires were correlated with AI internal details at the p < 0.001 level. Reducing this threshold to p < 0.05 identified a weak negative correlation with MEQ Coherence. The established memory questionnaires, therefore, seemed not to correlate with memory performance as measured by internal details. Given that internal details is generally taken as the ‘gold standard’ for measuring autobiographical memory ability, and the memory questionnaires employed here are widely used, this result is surprising.

**Table 12.** Correlation coefficients of the memory questionnaires with AI internal details as a percentage of total utterances.

|  | AI Internal Details |  |
| --- | --- | --- |
|  | r | p |
| Memory Experience Questionnaire; Coherence | -0.14 | 0.041 |
| Memory Experience Questionnaire; Accessibility | 0.080 | 0.24 |
| Survey of Autobiographical Memory; Episodic | -0.073 | 0.29 |
| Memory Experience Questionnaire; Sharing | 0.038 | 0.58 |
| Memory Experience Questionnaire; Vividness | -0.01 | 0.89 |
| Subjective Memory Questionnaire | -0.002 | 0.98 |

#### 3.2.2. Questionnaires from the other groups

As the memory questionnaires showed limited correlations with AI internal details, we next examined whether any of the questionnaires from the other groups were correlated with AI internal details, with the results shown in Table 13. In short, none of the questionnaires were correlated with AI internal details at the p < 0.001 level. Even reducing this threshold to p < 0.05 identified only a weak negative correlation with the Visualizer questionnaire (r = −0.14). This is in stark contrast to the correlations of the other group questionnaires with the scene construction imagination task noted previously (Table 6), which showed multiple significant correlations.

**Table 13.**
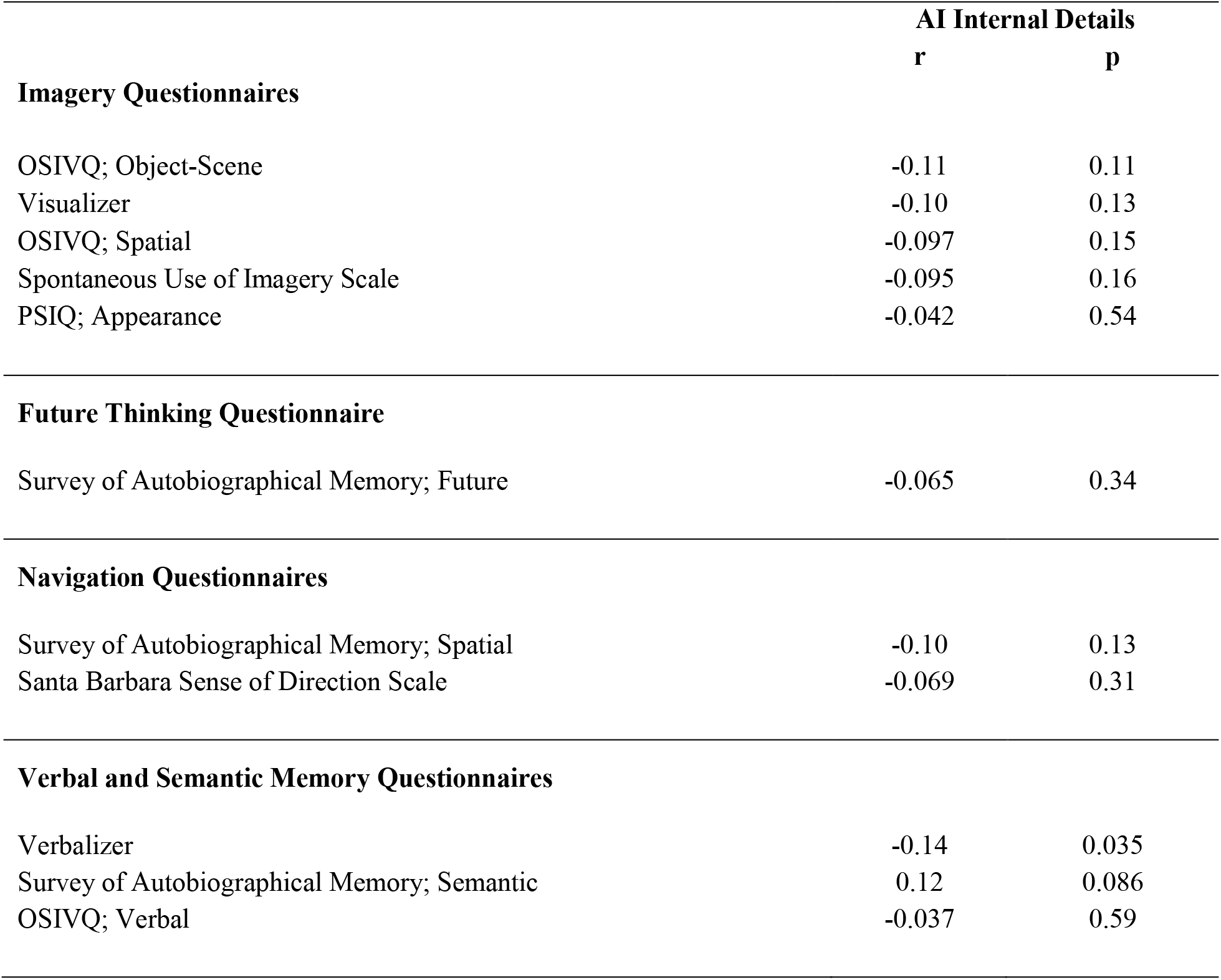
Correlation coefficients of the other group questionnaires with AI internal details as a percentage of total utterances.

#### 3.2.3. One Sentence Questionnaire

In line with the established questionnaires, none of the questions on the One Sentence Questionnaire were significantly related to AI internal details (Table 14).

**Table 14.** Correlation coefficients of the One Sentence Questionnaire questions with AI internal details as a percentage of total utterances.

|  | AI Internal Details |  |
| --- | --- | --- |
|  | r | p |
| **Imagery Questions** |  |  |
| One Sentence: Imagery Ability | 0.069 | 0.31 |
| One Sentence: Imagery as a Scene | 0.046 | 0.50 |
| One Sentence: Imagery Use | 0.042 | 0.53 |
| **Memory Questions** |  |  |
| One Sentence: Memory in Imagery | 0.12 | 0.081 |
| One Sentence: Memory Ability | 0.11 | 0.11 |
| One Sentence: Memory in Scene Imagery | 0.053 | 0.44 |
| One Sentence: Memory in Words | -0.057 | 0.41 |
| **Future Thinking Questions** |  |  |
| Future Thinking in Imagery | 0.059 | 0.39 |
| Future Thinking Ability | 0.022 | 0.74 |
| Future Thinking in Scene Imagery | 0.011 | 0.88 |
| Future Thinking in Words | 0.004 | 0.95 |
| **Navigation Questions** |  |  |
| One Sentence: Navigation in Words | 0.090 | 0.19 |
| One Sentence: Navigation in Scene Imagery | 0.089 | 0.19 |
| One Sentence: Navigation in Imagery | 0.026 | 0.71 |
| One Sentence: Navigation Ability | 0.008 | 0.91 |

#### 3.2.4. Additional investigations into AI internal details

The analyses reported above were performed with AI internal details averaged across all of the memory ages (childhood, teenage, adulthood, last year), and combining all of the sub-categories of internal details (events, place, time, perceptual, emotion). It could be argued that use of this combined data introduced additional noise, blurring potential associations.

We, therefore, performed all of the correlations again, but separately for each of the four time points (childhood, teenage, adulthood, last year) as well as combining the two older memory categories (childhood, teenage) to create an approximation of “remote” memory and the two more recent memory categories (adulthood, last year) to create an approximation of “recent” memory. In none of these additional 192 correlations was any relationship between questionnaires and AI internal details found (at our p < 0.001 statistical threshold; see full details in Supplementary Materials, Tables S5-7).

We also separately examined the five sub-categories of internal details (events, place, time, perceptual, emotion, each averaged across all of the time points, and represented as a percentage of total utterances). From among these additional 160 correlations, only one significant relationship was identified - a negative correlation between the internal events sub-category and the Verbalizer questionnaire (r = −0.24, Supplementary Materials, Tables S8-10).

Moreover, we conducted all of these analyses again, but using only the raw number of internal details (instead of the number of internal details as a percentage of total utterances). Doing so identified only four significant relationships at our p < 0.001 statistical threshold, all for the MEQ Sharing subscale (Supplementary Materials, Tables S11-16). This subscale was positively associated with the total number of internal details when averaged across all the time points (r = 0.26), and was specifically associated with the adulthood memory category (r = 0.25) and the “recent” memory category (r = 0.26). Investigation into the internal details sub-categories found that this positive correlation was specific to the internal events sub-category (r = 0.25). Given that the MEQ Sharing subscale asks about how often an individual talks about their memories, it is likely that these relationships are a reflection of participants’ verbosity.

It seems, therefore, that combining across time points and internal detail sub-categories did not adversely affect the sensitivity of the results.

#### 3.2.5. Autobiographical memory summary

Neither the established questionnaires nor the questions from our experimental One Sentence Questionnaire correlated with autobiographical memory internal details.

### 3.3. Autobiographical memory vividness

#### 3.3.1. Questionnaires from the autobiographical memory group

While the memory questionnaires (and indeed questionnaires in general) were not correlated with the main outcome measure of the AI, in contrast, the memory questionnaires were correlated with performance on the scene construction imagination task (see Table 6). As noted above, this could be because the memory questionnaires actually relate to the imagery experience of memories, rather than the number of details generated. Another measure collected from the AI is a participant rating of the vividness of memory recall. A correlation analysis revealed that memory vividness was measuring a different cognitive construct than the number of internal details generated, as there was no correlation between these two measures (r = 0.067, p = 0.33). Consequently, we examined whether the memory questionnaires were instead correlated with the vividness of memory recall.

As can be seen in Table 15, nearly all of the memory questionnaires significantly correlated with AI vividness at p < 0.001.

**Table 15.** Correlation coefficients of the memory questionnaires with AI vividness.

|  | AI Vividness |  |
| --- | --- | --- |
|  | r | p |
| Memory Experience Questionnaire; Vividness | 0.38 | < 0.001 |
| Subjective Memory Questionnaire | 0.37 | < 0.001 |
| Survey of Autobiographical Memory; Episodic | 0.37 | < 0.001 |
| Memory Experience Questionnaire; Accessibility | 0.33 | < 0.001 |
| Memory Experience Questionnaire; Coherence | 0.28 | < 0.001 |
| Memory Experience Questionnaire; Sharing | 0.23 | 0.001 |

Given that nearly all of the memory questionnaires were correlated with AI vividness, we next wanted to establish if there were any differences between these correlations. No differences were found between any of the correlations (Table 16).

**Table 16.**
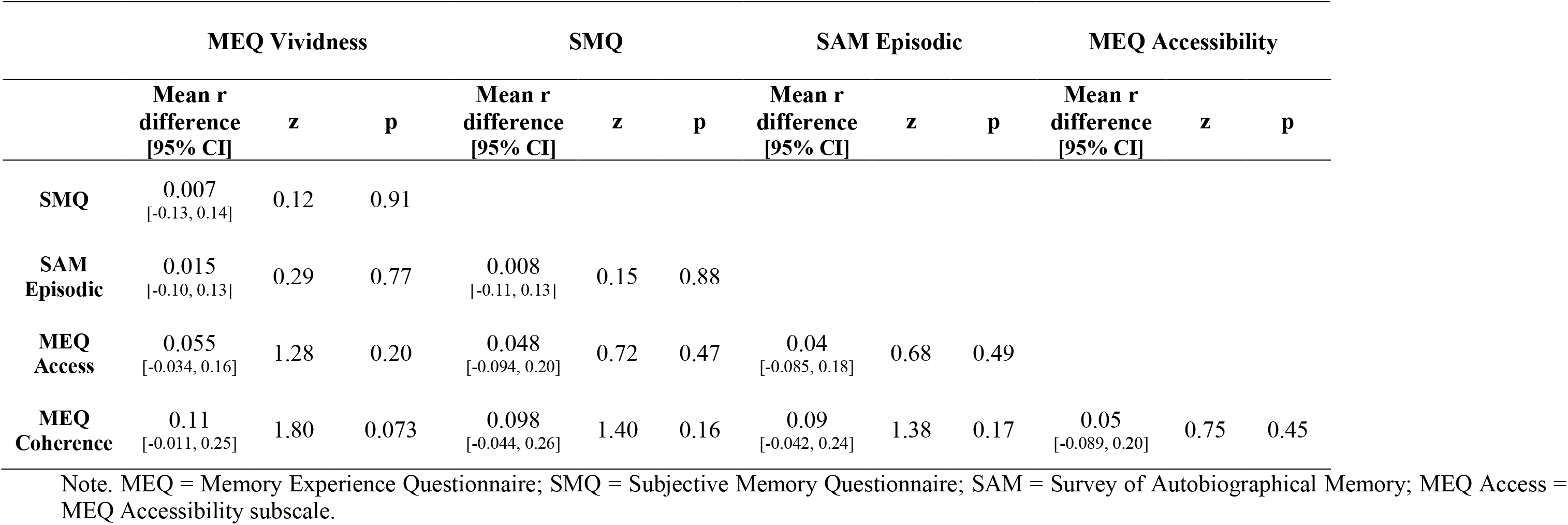
Comparisons between the correlation coefficients of the memory questionnaires with AI memory vividness. The difference in correlation is the questionnaire in the column less the questionnaires in the row.

|  | MEQ Vividness |  |  | SMQ |  |  | SAM Episodic |  |  | MEQ Accessibility |  |  |
| --- | --- | --- | --- | --- | --- | --- | --- | --- | --- | --- | --- | --- |
|  | Mean r<br>difference<br>[95% CI] | z | p | Mean r<br>difference<br>[95% CI] | z | p | Mean r<br>difference<br>[95% CI] | z | p | Mean r<br>difference<br>[95% CI] | z | p |
| SMQ | 0.007<br>[-0.13, 0.14] | 0.12 | 0.91 |  |  |  |  |  |  |  |  |  |
| SAM<br>Episodic | 0.015<br>[-0.10, 0.13] | 0.29 | 0.77 | 0.008<br>[-0.11, 0.13] | 0.15 | 0.88 |  |  |  |  |  |  |
| MEQ<br>Access | 0.055<br>[-0.034, 0.16] | 1.28 | 0.20 | 0.048<br>[-0.094, 0.20] | 0.72 | 0.47 | 0.04<br>[-0.085, 0.18] | 0.68 | 0.49 |  |  |  |
| MEQ<br>Coherence | 0.11<br>[-0.011, 0.25] | 1.80 | 0.073 | 0.098<br>[-0.044, 0.26] | 1.40 | 0.16 | 0.09<br>[-0.042, 0.24] | 1.38 | 0.17 | 0.05<br>[-0.089, 0.20] | 0.75 | 0.45 |

Note. MEQ = Memory Experience Questionnaire; SMQ = Subjective Memory Questionnaire; SAM = Survey of Autobiographical Memory; MEQ Access = MEQ Accessibility subscale.

Next, we sought to ascertain whether the memory questionnaires were all tapping into the same construct, or if different questionnaires were measuring distinct aspects of AI vividness. To do this, we performed a multiple regression using the memory questionnaires that were significantly correlated with AI vividness (at p < 0.001). The results are shown in Table 17. While the full regression model was significant [F(6,210) = 8.71,p < 0.001, R^2^ = 0.20, Adj. R^2^ = 0.18], only the SMQ questionnaire was significantly related to AI vividness.

**Table 17.** Multiple regression of all the memory questionnaires significantly correlated with AI vividness.

|  | **Beta**<br>**(95% CI)** | **Standardised**<br>**Beta** | **t** | **p** |
| --- | --- | --- | --- | --- |
| SMQ | 0.009 (0.002, 0.017) | 0.21 | 2.54 | 0.012 |
| MEQ Sharing | 0.014 (-0.008, 0.037) | 0.089 | 1.26 | 0.21 |
| MEQ Vividness | 0.021 (-0.019, 0.061) | 0.12 | 1.03 | 0.30 |
| SAM Episodic | 0.005 (-0.005, 0.016) | 0.091 | 0.98 | 0.33 |
| MEQ Coherence | 0.008 (-0.019, 0.035) | 0.045 | 0.59 | 0.56 |
| MEQ Accessibility | 0.007 (-0.036, 0.050) | 0.030 | 0.30 | 0.76 |

Note. SMQ = Subjective Memory Questionnaire; MEQ = Memory Experience Questionnaire; SAM = Survey of Autobiographical Memory

#### 3.3.2. Questionnaires from the other groups

Memory vividness is potentially more of an imagery construct rather than being specific to memory. Therefore, we wanted to know if the questionnaires from the other groups (namely, imagination, future thinking and navigation), were also correlated with AI vividness, with a particular interest in the imagery questionnaires. To check that responses to questionnaires in general were not correlating with AI vividness, the control questionnaires were again included.

As is evident in Table 18, the majority of questionnaires from the other groups correlated with AI vividness. Importantly, the control questionnaires did not correlate with AI vividness. This, therefore, supports the idea that the significant correlations are not simply due to artificial constructs, but arise from aspects of the questionnaires. Of additional note, two of the imagery questionnaires - the Visualizer and OSIVQ Spatial subscale also did not correlate with AI vividness. This is particularly interesting because these are the two imagery questionnaires that also did not correlate with the scene construction imagination task. Overall, therefore, the correlations of the memory questionnaires with AI vividness seemed to extend beyond memory, with instead correlations being apparent whenever there was a visual imagery element to the questionnaire.

**Table 18.** Correlation coefficients of the other group questionnaires with AI vividness.

|  | AI Vividness |  |
| --- | --- | --- |
|  | r | p |
| **Imagery Questionnaires** |  |  |
| Object Spatial Imagery and Verbal Questionnaire; Object-Scene | 0.51 | < 0.001 |
| Plymouth Sensory Imagery Questionnaire; Appearance | 0.49 | < 0.001 |
| Spontaneous Use of Imagery Scale | 0.28 | < 0.001 |
| Visualizer | 0.082 | 0.23 |
| Object Spatial Imagery and Verbal Questionnaire; Spatial | 0.044 | 0.52 |
| **Future Thinking Questionnaire** |  |  |
| Survey of Autobiographical Memory; Future | 0.28 | < 0.001 |
| **Navigation Questionnaires** |  |  |
| Santa Barbara Navigation Questionnaire | 0.26 | < 0.001 |
| Survey of Autobiographical Memory; Spatial | 0.25 | < 0.001 |
| **Verbal and Semantic Memory Questionnaires** |  |  |
| Survey of Autobiographical Memory; Semantic | 0.18 | 0.009 |
| Object Spatial Imagery and Verbal Questionnaire; Verbal | 0.009 | 0.89 |
| Verbalizer | 0.020 | 0.77 |

On the one hand, this may be expected – the vividness of an autobiographical memory is not a pure memory measure, and shares substantial overlap with that of imagination/imagery. Nevertheless, our findings suggest that widely used memory questionnaires may not only be reflecting an individual’s memory vividness, but also acting as an indirect measure of imagery ability.

To examine this in more detail, we next compared the significant correlations of the questionnaires from the other groups and AI vividness with the significantly correlated memory questionnaires. This allowed us to assess whether either type of questionnaire was outperforming the other. As shown in Table 19, the OSIVQ Object-Scene subscale and the PSIQ had higher correlations than all the established memory questionnaires (significant for all the comparisons with the OSIVQ Object-Scene subscale, but only for the MEQ Accessibility and MEQ Coherence subscales for the PSIQ). On the other hand, all the other questionnaires from the other groups performed at around the same level as the main memory questionnaires. That is, none of the memory questionnaires had greater correlations with AI vividness than any of the other questionnaires. Thus, the specificity of the memory questionnaires seems to be in question.

**Table 19.**
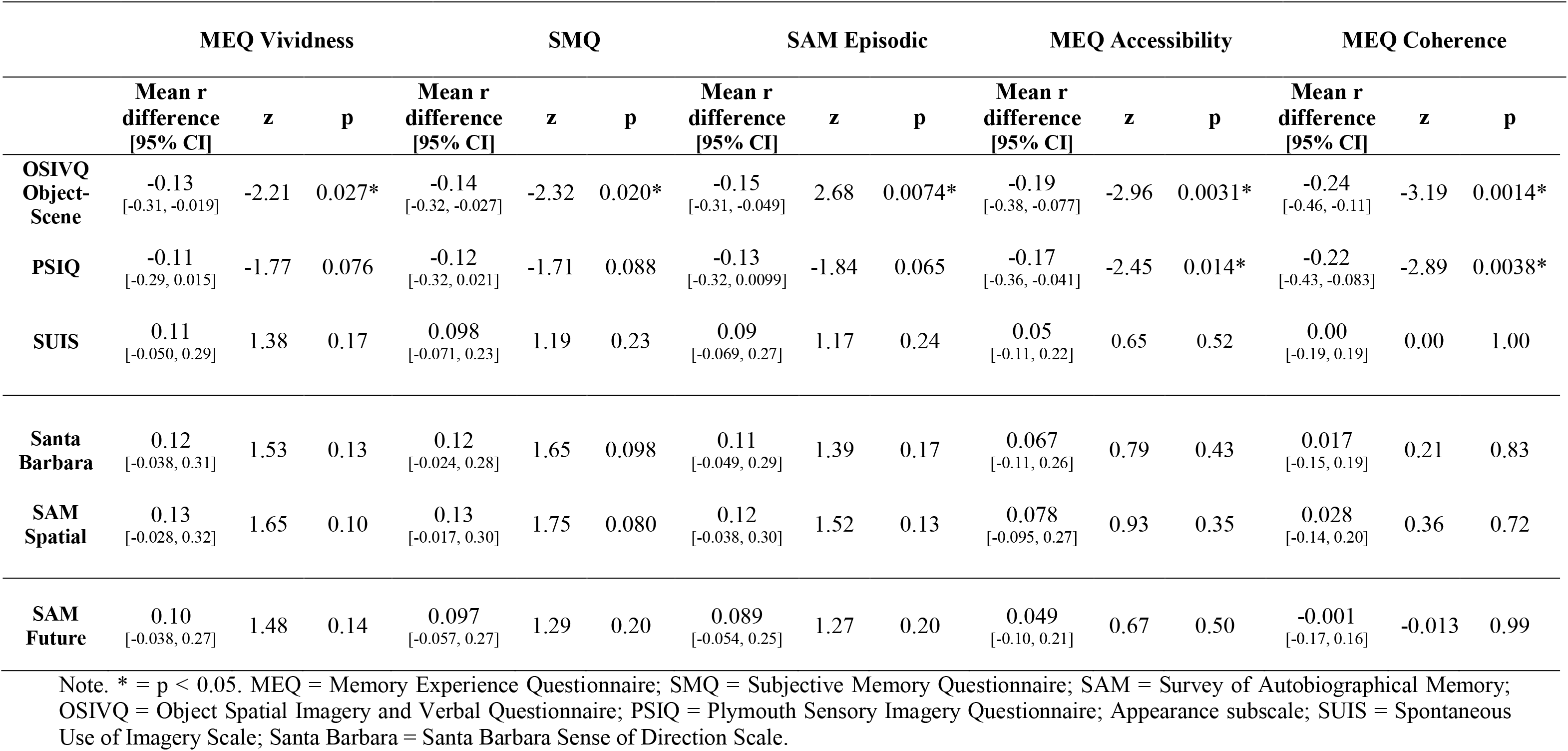
Comparisons between the correlation coefficients of the significantly correlating memory questionnaires with the questionnaires from the other groups which correlated with AI vividness (at p < 0.001). The difference in correlation is the memory questionnaires less the questionnaires from the other groups (a negative value shows higher correlation for the non-memory questionnaire).

Given the high levels of correlations of the questionnaires in the other groups (in addition to the memory questionnaires) with AI vividness, we next performed a multiple regression to ascertain if the questionnaires were correlating with AI vividness via the same construct or if there were distinct contributions from the different types of questionnaires.

The results of the multiple regression can be seen in Table 20. The full regression model was found to be significant [F(11,205) = 10.39, p < 0.001, R^2^ = 0.36, Adj. R^2^ = 0.32], with two questionnaires contributing to AI vividness; the PSIQ and the OSIVQ Object-Scene subscale. This suggests that for AI vividness, the PSIQ and the OSIVQ Object-Scene subscale were tapping into different constructs.

**Table 20.** Multiple regression of all the questionnaires significantly correlated with AI vividness.

|  | **Beta**<br>**(95% CI)** | **Standardised**<br>**Beta** | **t** | **p** |
| --- | --- | --- | --- | --- |
| PSIQ | 0.036 (0.017, 0.054) | 0.27 | 3.82 | < 0.001 |
| OSIVQ Object-Scene | 0.39 (0.16, 0.61) | 0.32 | 3.36 | 0.001 |
| MEQ Coherence | 0.014 (-0.011, 0.039) | 0.078 | 1.08 | 0.28 |
| SMQ | 0.003 (-0.004, 0.010) | 0.072 | 0.91 | 0.37 |
| Spontaneous Use Of Imagery Scale | -0.005 (-0.019, 0.009) | -0.048 | -0.66 | 0.51 |
| Santa Barbara | 0.003 (-0.006, 0.012) | 0.054 | 0.56 | 0.58 |
| SAM; Spatial | 0.002 (-0.007, 0.011) | 0.039 | 0.41 | 0.68 |
| SAM; Future | -0.002 (-0.01, 0.007) | -0.026 | -0.39 | 0.70 |
| MEQ Vividness | 0.004 (-0.032, 0.040) | 0.024 | 0.22 | 0.82 |
| SAM Episodic | -0.001 (-0.011, 0.009) | -0.02 | -0.22 | 0.83 |
| MEQ Accessibility | 0.004 (-0.035, 0.043) | 0.018 | 0.20 | 0.84 |

Note. PSIQ = Plymouth Sensory Imagery Questionnaire; Appearance subscale; OSIVQ = Object Spatial Imagery and Verbal Questionnaire; MEQ = Memory Experience Questionnaire; SMQ = Subjective Memory Questionnaire; Santa Barbara = Santa Barbara Sense of Direction Scale; SAM = Survey of Autobiographical Memory.

The multiple regression, therefore, suggests that if one wants to examine AI vividness then memory questionnaires are not required. Instead, imagery questionnaires have the best association with AI vividness. Overall, the main cognitive construct associated with AI vividness seemed to be subjective reports of visual imagery ability.

#### 3.3.3. One Sentence Questionnaire

We next considered AI vividness in relation to our exploratory One Sentence Questionnaire. First, we assessed whether or not the memory questions from the One Sentence Questionnaire correlated with AI vividness, finding that they did, with the exception of the Memory in Words question (One Sentence Memory in Imagery: r = 0.36, p < 0.001; One Sentence Memory in Scene Imagery: r = 0.36, p < 0.001; One Sentence Memory Ability: r = 0.28, p < 0.001; One Sentence Memory in Words: r = −0.062, p = 0.36).

We then tested how the significantly associated One Sentence Questionnaire memory questions performed in comparison to the established memory questionnaires that correlated with AI vividness. The results are shown in Table 21. No differences in correlations were found. That is, the memory questions from the One Sentence Questionnaire were correlated with AI vividness to a similar degree as the longer, and more time consuming, established questionnaires.

**Table 21.** Comparisons between the correlation coefficients of the established memory questionnaires and the memory questions from the One Sentence Questionnaire that were associated with AI memory vividness. The difference in correlation is the questionnaire in the column less the questionnaires in the row (a negative value shows higher correlation for the questions of the One Sentence Questionnaire).

|  | MEQ Vividness |  |  | SMQ |  |  | SAM Episodic |  |  | MEQ Accessibility |  |  | MEQ Coherence |  |  |
| --- | --- | --- | --- | --- | --- | --- | --- | --- | --- | --- | --- | --- | --- | --- | --- |
|  | Mean r<br>difference<br>[95% CI] | z | p | Mean r<br>difference<br>[95% CI] | z | p | Mean r<br>difference<br>[95% CI] | z | p | Mean r<br>difference<br>[95% CI] | z | p | Mean r<br>difference<br>[95% CI] | z | p |
| Memory in<br>Imagery | 0.018<br>[-0.14, 0.19] | 0.25 | 0.80 | 0.011<br>[-0.16, 0.18] | 0.15 | 0.88 | 0.003<br>[-0.17, 0.17] | 0.040 | 0.97 | -0.037<br>[-0.21, 0.12] | -0.50 | 0.62 | -0.087<br>[-0.27, 0.080] | -1.07 | 0.28 |
| Memory in<br>Scene<br>Imagery | 0.026<br>[-0.14, 0.20] | 0.35 | 0.73 | 0.019<br>[-0.15, 0.19] | 0.26 | 0.80 | 0.011<br>[-0.16, 0.19] | 0.14 | 0.89 | -0.029<br>[-0.20, 0.14] | -0.38 | 0.70 | -0.079<br>[-0.26, 0.087] | -0.98 | 0.33 |
| Memory<br>Ability | 0.097<br>[-0.0062, 0.22] | 1.86 | 0.064 | 0.09<br>[-0.043, 0.25] | 1.37 | 0.17 | 0.082<br>[-0.028, 0.21] | 1.51 | 0.13 | 0.042<br>[-0.077, 0.17] | 0.74 | 0.46 | -0.008<br>[-0.16, 0.14] | -0.12 | 0.91 |

Note. MEQ = Memory Experience Questionnaire; SMQ = Subjective Memory Questionnaire; SAM = Survey of Autobiographical Memory.

Next, we sought to ascertain if, like the established questionnaires, the One Sentence Questionnaire questions designed to look at other cognitive functions (namely, imagination, future thinking and navigation), were also associated with AI vividness. The results can be seen in Table 22. Nearly all the questions correlated with AI vividness. The only ones that did not were those pertaining to verbal processing.

**Table 22.** Correlation coefficients of the other group questions from the One Sentence Questionnaire with AI vividness.

|  | AI Vividness |  |
| --- | --- | --- |
|  | r | p |
| **Imagery Questions** |  |  |
| One Sentence: Imagery Ability | 0.39 | < 0.001 |
| One Sentence: Imagery as a Scene | 0.31 | < 0.001 |
| One Sentence: Imagery Use | 0.29 | < 0.001 |
| **Future Thinking Questions** |  |  |
| One Sentence: Future Thinking in Imagery | 0.37 | < 0.001 |
| One Sentence: Future Thinking in Scene Imagery | 0.32 | < 0.001 |
| One Sentence: Future Thinking Ability | 0.31 | < 0.001 |
| One Sentence: Future Thinking in Words | 0.063 | 0.36 |
| **Navigation Questions** |  |  |
| One Sentence: Navigation in Scene Imagery | 0.29 | < 0.001 |
| One Sentence: Navigation in Imagery | 0.28 | < 0.001 |
| One Sentence: Navigation Ability | 0.24 | < 0.001 |
| One Sentence: Navigation in Words | 0.019 | 0.79 |

Finally, we investigated whether the One Sentence Questionnaire questions from the other groups were correlated with AI vividness to the same extent as the memory questions from the One Sentence Questionnaire. No differences were identified (Table 23). In other words, unlike the established imagery questionnaires, the One Sentence Questionnaire imagery questions did not predict AI vividness better than the memory questions, but they were no worse than the memory questions.

**Table 23.** Comparisons between the correlation coefficients of the One Sentence Questionnaire memory questions and the One Sentence Questionnaire questions from the other groups which correlated with AI vividness (at p < 0.001). The difference in correlation is the memory questions less the questions from the other groups (a negative value shows higher correlation for the non-memory questions).

|  | One Sentence: Memory in Imagery |  |  | One Sentence: Memory in Scene Imagery |  |  | One Sentence: Memory Ability |  |  |
| --- | --- | --- | --- | --- | --- | --- | --- | --- | --- |
|  | Mean r difference<br>[95% CI] | z | p | Mean r difference<br>[95% CI] | z | p | Mean r difference<br>[95% CI] | z | p |
| Imagery Ability | -0.028<br>[-0.17, 0.11] | -0.46 | 0.65 | -0.036<br>[-0.19, 0.11] | -0.55 | 0.58 | -0.011<br>[-0.28, 0.037] | -1.50 | 0.13 |
| Imagery as a Scene | 0.056<br>[-0.082, 0.21] | 0.85 | 0.39 | 0.048<br>[-0.084, 0.19] | 0.77 | 0.44 | -0.023<br>[-0.18, 0.13] | -0.31 | 0.75 |
| Imagery Use | 0.078<br>[-0.059, 0.23] | 1.17 | 0.24 | 0.07<br>[-0.075, 0.23] | 1.0 | 0.32 | -0.001<br>[-0.16, 0.16] | -0.014 | 0.99 |
| Navigation in Scene Imagery | 0.077<br>[-0.070, 0.24] | 1.08 | 0.28 | 0.069<br>[-0.075, 0.23] | 0.99 | 0.32 | -0.002<br>[-0.18, 0.17] | -0.024 | 0.98 |
| Navigation in Imagery | 0.079<br>[-0.061, 0.24] | 1.16 | 0.25 | 0.071<br>[-0.084, 0.24] | 0.95 | 0.34 | 0.0<br>[-0.18, 0.18] | 0.0 | 1.0 |
| Navigation Ability | 0.13<br>[-0.031, 0.31] | 1.61 | 0.11 | 0.12<br>[-0.049, 0.31] | 1.43 | 0.15 | 0.049<br>[-0.13, 0.23] | 0.58 | 0.56 |
| Future in Imagery | -0.011<br>[-0.15, 0.12] | -0.19 | 0.85 | -0.019<br>[-0.16, 0.12] | -0.31 | 0.76 | -0.09<br>[-0.26, 0.056] | -1.26 | 0.21 |
| Future in Scene Imagery | 0.046<br>[-0.094, 0.20] | 0.70 | 0.49 | 0.038<br>[-0.083, 0.17] | 0.67 | 0.50 | -0.033<br>[-0.21, 0.13] | -0.42 | 0.68 |
| Future Ability | 0.052<br>[-0.097, 0.21] | 0.74 | 0.46 | 0.044<br>[-0.11, 0.21] | 0.60 | 0.55 | -0.027<br>[-0.20, 0.14] | -0.34 | 0.73 |

#### 3.3.4. Autobiographical memory vividness summary

The main associations between the questionnaires and AI vividness can be seen in Figure 2.

**Figure 2.**
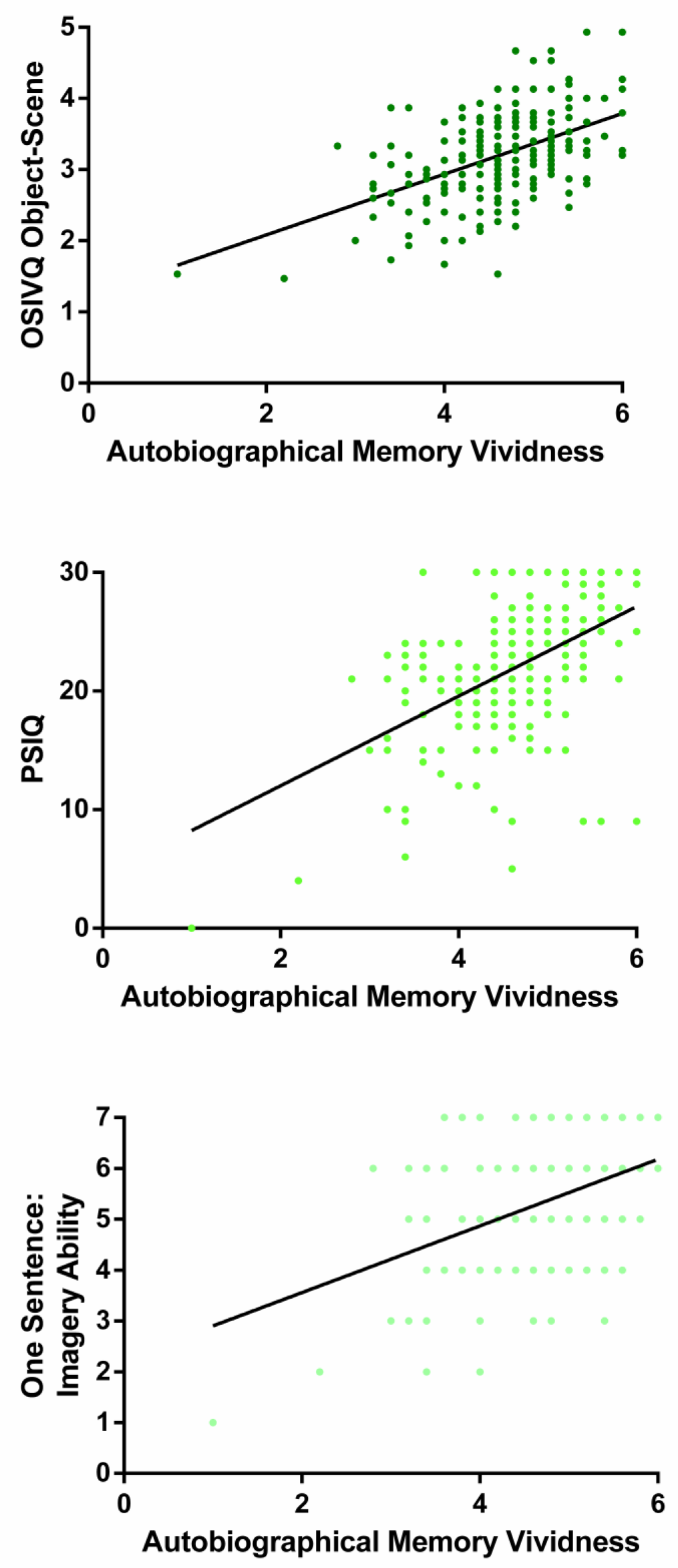
Correlations of the PSIQ, OSIVQ Object-Scene and One Sentence: Imagery Ability question with AI vividness.

Considering our three research questions, first, memory questionnaires seem to relate to autobiographical memory vividness, rather than internal details. Second, imagination, future thinking and navigation questionnaires were also associated with autobiographical memory vividness. Furthermore, the imagery questionnaires had greater associations with autobiographical memory vividness than the other questionnaires. Imagery seems, therefore, to underlie memory vividness, and it may be that memory questionnaires are in fact only indirect measures of imagery ability; an important point to note when deploying questionnaires in future memory research. Third, the memory questions from our One Sentence Questionnaire preformed similarly to the established, and more time consuming, memory questionnaires. They also performed comparably with the other (imagery based) questions pertaining to other cognitive functions on the One Sentence Questionnaire.

### 3.4. Future thinking

#### 3.4.1. Future thinking questionnaire

We next looked at the relationship between the future thinking questionnaire and the future thinking task. As a brief reminder, we have already noted that the future thinking questionnaire correlated with both the scene construction imagination task and AI vividness. Of note, as future thinking has become a topic of heightened interest more recently than imagery and memory, we have only one questionnaire available with which to directly assess it.

We first examined whether or not the future thinking questionnaire was correlated with performance on the future thinking task, finding only a weak correlation (r = 0.21, p = 0.002) that was not significant at our p < 0.001 level.

#### 3.4.2. Questionnaires from the other groups

Given the previous overlaps in questionnaire and task correlations, we next wanted to see if the questionnaires from the other groups (namely, imagery, memory and navigation), were correlated with future thinking performance. In particular this allowed us to test whether an imagery and/or an autobiographical memory process was associated with future thinking ability. As before, to check that questionnaires in general were not simply correlating with future thinking, the control questionnaires were included.

As shown in Table 24, the PSIQ, the OSIVQ Object-Scene subscale and the MEQ Coherence subscale all correlated with future thinking performance at p < 0.001. These questionnaires were also associated with scene construction performance and AI vividness, in particular the PSIQ and OSIVQ Object-Scene subscale. In addition, a number of the other imagery-based questionnaires were also correlated with future thinking at a weaker level, similar to that of the future thinking questionnaire itself. Navigation questionnaires, on the other hand, were not correlated with performance on the future thinking task. Overall, therefore, as with AI vividness, the imagery questionnaires may be particularly associated with future thinking performance.

**Table 24.** Correlation coefficients of the questionnaires from the other groups with future thinking performance.

|  | Future Thinking |  |
| --- | --- | --- |
|  | r | p |
| **Imagery Questionnaires** |  |  |
| Plymouth Sensory Imagery Questionnaire; Appearance | 0.30 | < 0.001 |
| Object Spatial Imagery and Verbal Questionnaire; Object-Scene | 0.24 | < 0.001 |
| Spontaneous Use of Imagery Scale | 0.17 | 0.014 |
| Visualizer | 0.068 | 0.32 |
| Object Spatial Imagery and Verbal Questionnaire; Spatial | -0.017 | 0.80 |
| **Memory Questionnaires** |  |  |
| Memory Experience Questionnaire; Coherence | 0.24 | < 0.001 |
| Survey of Autobiographical Memory; Episodic | 0.23 | 0.001 |
| Memory Experience Questionnaire; Vividness | 0.22 | 0.001 |
| Subjective Memory Questionnaire | 0.17 | 0.013 |
| Memory Experience Questionnaire; Sharing | 0.16 | 0.017 |
| Memory Experience Questionnaire; Accessibility | 0.16 | 0.021 |
| **Navigation Questionnaires** |  |  |
| Santa Barbara Sense of Direction Scale | 0.10 | 0.13 |
| Survey of Autobiographical Memory; Spatial | 0.092 | 0.18 |
| **Verbal and Semantic Memory Questionnaires** |  |  |
| Object Spatial Imagery and Verbal Questionnaire; Verbal | 0.13 | 0.049 |
| Survey of Autobiographical Memory; Semantic | 0.088 | 0.20 |
| Verbalizer | 0.079 | 0.25 |

Given the multiple significant correlations, we performed a multiple regression to test whether the different questionnaires were tapping into the same cognitive construct or if they represented different aspects of future thinking. As can be seen in Table 25, two questionnaires were identified as being associated with future thinking [regression model statistics: F(3,213) = 10.26, p < 0.001, R^2^ = 0.12, Adj. R^2^ = 0.11]; the PSIQ and the MEQ Coherence subscale. Overall, therefore, the multiple regression showed two constructs associated with future thinking; visual imagery vividness and coherence. Of note, the PSIQ was also strongly associated with both the scene construction task and AI vividness.

**Table 25.**
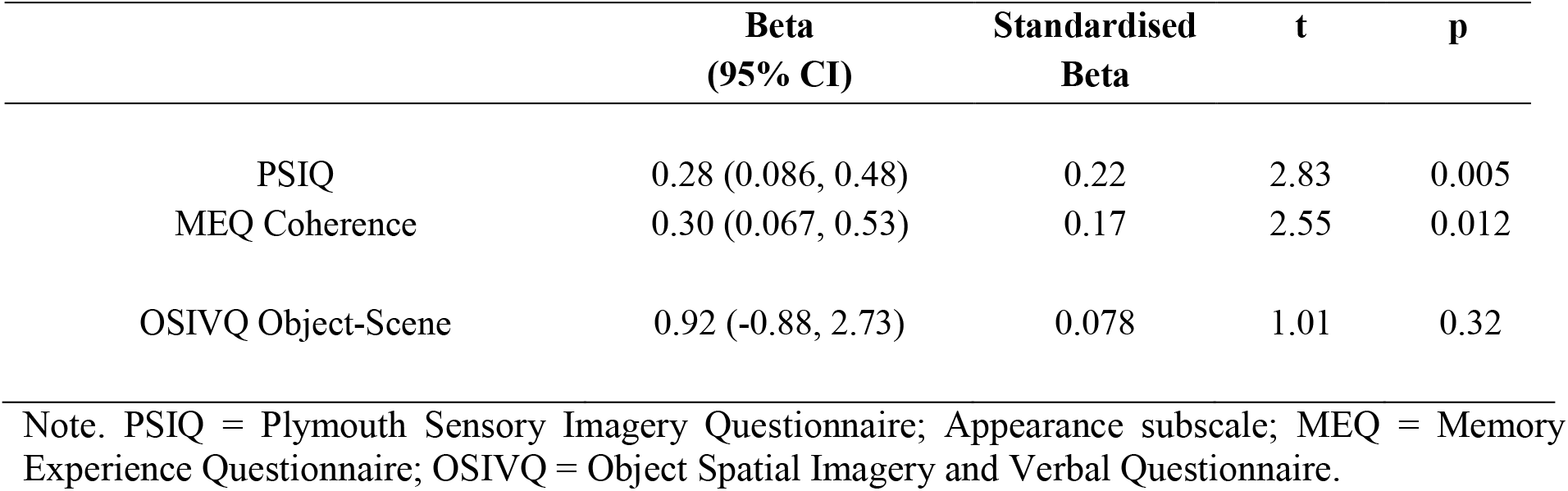
Multiple regression of all the questionnaires significantly correlated with future thinking performance.

|  | **Beta**<br>**(95% CI)** | **Standardised**<br>**Beta** | **t** | **p** |
| --- | --- | --- | --- | --- |
| PSIQ | 0.28 (0.086, 0.48) | 0.22 | 2.83 | 0.005 |
| MEQ Coherence | 0.30 (0.067, 0.53) | 0.17 | 2.55 | 0.012 |
| OSIVQ Object-Scene | 0.92 (-0.88, 2.73) | 0.078 | 1.01 | 0.32 |

Note. PSIQ = Plymouth Sensory Imagery Questionnaire; Appearance subscale; MEQ = Memory Experience Questionnaire; OSIVQ = Object Spatial Imagery and Verbal Questionnaire.

#### 3.4.3. One Sentence Questionnaire

We first assessed whether the future thinking questions from the One Sentence Questionnaire were correlated with future thinking performance. Mirroring the established SAM Future subscale, none of the future thinking One Sentence Questionnaire questions correlated with future thinking performance (One Sentence Future Thinking in Scene Imagery: r = 0.17, p = 0.012; One Sentence Future Thinking in Imagery: r = 0.17, p = 0.013; One Sentence Future Thinking Ability: r = 0.16, p = 0.017; One Sentence Future Thinking in Words: r = 0.026, p = 0.70).

Next, we sought to ascertain if, like the established questionnaires, the One Sentence Questionnaire questions designed to look at other cognitive functions were associated with performance on the future thinking task. The results can be seen in Table 26. The only significant correlation (at p < 0.001) was one of the imagery questions – One Sentence Imagery as a Scene. This is in line with the established questionnaires, finding that the PSIQ (which specifically measures scene imagery) significantly correlated with future thinking performance.

**Table 26.** Correlation coefficients of the One Sentence Questionnaire questions from other groups with future thinking performance.

|  | Future Thinking |  |
| --- | --- | --- |
|  | r | p |
| **Imagery Questions** |  |  |
| One Sentence: Imagery as a Scene | 0.25 | < 0.001 |
| One Sentence: Imagery Ability | 0.22 | 0.001 |
| One Sentence: Imagery Use | 0.20 | 0.003 |
| **Memory Questions** |  |  |
| One Sentence: Memory in Imagery | 0.19 | 0.006 |
| One Sentence: Memory in Scene Imagery | 0.18 | 0.008 |
| One Sentence: Memory Ability | 0.15 | 0.026 |
| One Sentence: Memory in Words | -0.056 | 0.14 |
| **Navigation Questions** |  |  |
| One Sentence: Navigation in Scene Imagery | 0.15 | 0.032 |
| One Sentence: Navigation in Imagery | 0.13 | 0.05 |
| One Sentence: Navigation Ability | 0.10 | 0.13 |
| One Sentence: Navigation in Words | 0.076 | 0.27 |

#### 3.4.4. Future thinking summary

The main associations between the questionnaires and future thinking performance are shown in Figure 3.

**Figure 3.**
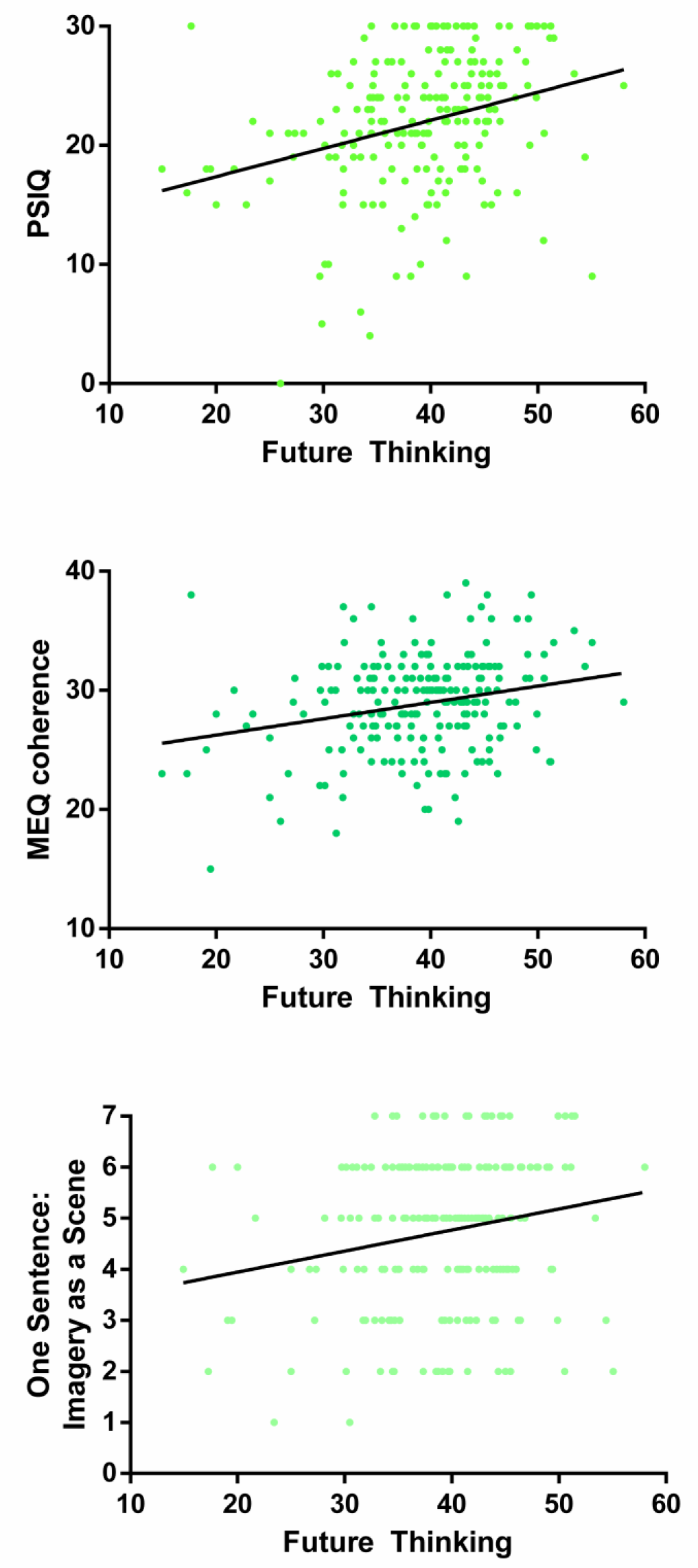
Graphs showing the correlations between the PSIQ, MEQ Coherence subscale and One Sentence: Imagery as a Scene question with future thinking performance.

Considering our three research questions, first, the established future thinking questionnaire was only weakly linked to future thinking performance. Second, questionnaires from the other groups and their relationship with future thinking illustrated the importance of imagery for reflecting future thinking performance. Overall, and once again, imagery questionnaires seem to be most strongly related to imagination, autobiographical memory and future thinking ability. Third, the future thinking questions from our One Sentence Questionnaire were also only weakly associated with future thinking performance. Instead, it was the scene imagery question from the One Sentence Questionnaire that was significantly associated with future thinking performance – mirroring the established questionnaires.

### 3.5. Navigation

#### 3.5.1. Questionnaires from the navigation group

Our final cognitive function of interest was navigation. Thus far, navigation questionnaires have been notable in not correlating with performance on scene construction, AI internal details or future thinking, although they did correlate with AI vividness. It may be, therefore, that navigation questionnaires are more specific in their line of questioning.

We first correlated the two established navigation questionnaires with performance on the navigation task. Both questionnaires correlated with navigation performance (SAM Spatial subscale: r = 0.33, p < 0.001; the Santa Barbara Sense of Direction Scale: r = 0.32, p < 0.001).

Second, we tested whether there was a difference in the correlations of the two questionnaires, finding no difference between them [mean r difference = 0.001 (95% CI = - 0.079, 0.10), z = 0.24, p =0.81].

Finally, we wanted to establish whether or not the two questionnaires were tapping into the same cognitive construct, or if they represented distinct aspects of navigation. We performed a multiple regression analysis using the two navigation questionnaires, finding that while the overall model was significant [F(2,214 = 13.92, p < 0.001, R^2^ = 0.12, Adj. R^2^ = 0.11], neither questionnaire was significantly associated with navigation performance [SAM Spatial: Beta value (95% CI) = 0.49 (−0.016, 1.0), Standardised beta value = 0.20, t = 1.91, p = 0.058; Santa Barbara Sense of Direction Scale: Beta value (95% CI) = 0.36 (−0.12, 0.84), Standardised beta value = 0.16, t = 1.47, p = 0.14]. This, therefore, suggests that both navigation questionnaires were tapping into the same cognitive construct.

#### 3.5.2. Questionnaires from the other groups

We next examined whether or not the other group questionnaires (namely, imagination, autobiographical memory and future thinking), were correlated with navigation performance. This was especially interesting given that navigation questionnaires were the least associated with these other tasks. As before, to check that questionnaires in general were not simply correlating with navigation performance, the control questionnaires were included.

The results are shown in Table 27. The only questionnaire from the other groups that correlated with navigation task performance was the OSIVQ Spatial subscale. These results are in line with the findings we described earlier, showing limited relationships between navigation questionnaires and imagination, autobiographical memory and future thinking. Navigation seems, therefore, to be a separate construct from the other three cognitive functions.

**Table 27.** Correlation coefficients of the other group questionnaires with navigation performance.

|  | Navigation |  |
| --- | --- | --- |
|  | r | p |
| **Imagery Questionnaires** |  |  |
| Object Spatial Imagery and Verbal Questionnaire; Spatial | 0.24 | < 0.001 |
| Visualizer | 0.17 | 0.01 |
| Plymouth Sensory Imagery Questionnaire; Appearance | 0.14 | 0.034 |
| Object Spatial Imagery and Verbal Questionnaire; Object-Scene | -0.044 | 0.52 |
| Spontaneous Use Of Imagery Scale | -0.026 | 0.71 |
| **Memory Questionnaires** |  |  |
| Memory Experience Questionnaire; Coherence | 0.12 | 0.072 |
| Subjective Memory Questionnaire | 0.056 | 0.41 |
| Survey of Autobiographical Memory; Episodic | -0.044 | 0.52 |
| Memory Experience Questionnaire; Vividness | 0.032 | 0.64 |
| Memory Experience Questionnaire; Accessibility | -0.028 | 0.68 |
| Memory Experience Questionnaire; Sharing | 0.009 | 0.89 |
| **Future Thinking Questionnaire** |  |  |
| Survey of Autobiographical Memory; Future | 0.016 | 0.81 |
| **Verbal and Semantic Memory Questionnaires** |  |  |
| Object Spatial Imagery and Verbal Questionnaire; Verbal | 0.061 | 0.37 |
| Survey of Autobiographical Memory; Semantic | 0.019 | 0.78 |
| Verbalizer | 0.005 | 0.95 |

Given the significant correlation of the OSIVQ Spatial subscale with navigation performance, we next wanted to know whether the navigation questionnaires outperformed the OSIVQ Spatial subscale, or whether they correlated with navigation task performance to the same extent. No differences between the questionnaires were found [OSIVQ Spatial vs SAM Spatial: mean r difference = −0.087 (95% CI = −0.24, 0.049), z = −1.29, p = 0.20; OSIVQ Spatial vs Santa Barbara Sense of Direction Scale: mean r difference = −0.077 (95% CI = - 0.23, 0.062), z = −1.13, p = 0.26].

Finally, we performed a multiple regression analysis to identify if the OSIVQ Spatial subscale was involved in navigation performance through the same or a different cognitive construct as the navigation questionnaires. As before, while the regression model was significant [F(3,213 = 9.94, p < 0.001, R^2^ = 0.12, Adj. R^2^ = 0.11], none of the questionnaires were associated with navigation performance [SAM Spatial: Beta value (95% CI) = 0.42 (−0.094, 0.94), Standardised beta value = 0.17, t = 1.61, p = 0.11; Santa Barbara Sense of Direction Scale: Beta value (95% CI) = 0.31 (−0.18, 0.80), Standardised beta value = 0.13, t = 1.25, p = 0.21; OSIVQ Spatial: Beta value (95% CI) = 5.32 (−2.36, 13.0), Standardised beta value = 0.10, t = 1.37, p = 0.18]. This, therefore, suggests that the OSIVQ Spatial subscale was tapping into the same construct as the navigation questionnaires.

#### 3.5.3. One Sentence Questionnaire

We next investigated the relationship between navigation performance and our exploratory One Sentence Questionnaire. First, we assessed whether or not the navigation questions from the One Sentence Questionnaire correlated with navigation task performance, finding that only the Navigation Ability question was significantly associated with navigation task performance at p < 0.001 (One Sentence: Navigation Ability: r = 0.30, p < 0.001; One Sentence: Navigation in Scene Imagery: r = 0.20, p = 0.003; One Sentence: Navigation in Imagery: r = 0.19, p = 0.004; One Sentence: Navigation in Words: r = −0.073, p = 0.28).

We then wanted to establish whether or not there were any differences in correlations between the established navigation questionnaires and the significant navigation question from the One Sentence Questionnaire. No differences between the correlations were identified [One Sentence Navigation Ability vs SAM Spatial: mean r difference (95% CI) = 0.027 (−0.065, 0.12), z = 0.62, p = 0.54; One Sentence Navigation Ability vs Santa Barbara Sense of Direction Scale: mean r difference (95% CI) = 0.017 (−0.071, 0.11), z = 0.41, p = 0.68].

Next, we sought to ascertain if the One Sentence Questionnaire questions designed to look at other cognitive functions were also associated with performance on the navigation task. As can be seen in Table 28, none of the other questions on the One Sentence Questionnaire were associated with navigation task performance.

**Table 28.** Correlation coefficients of the One Sentence Questionnaire questions from other groups with navigation task performance.

|  | Navigation |  |
| --- | --- | --- |
|  | r | p |
| **Imagery Questionnaires** |  |  |
| One Sentence: Imagery Ability | 0.038 | 0.58 |
| One Sentence: Imagery as a Scene | 0.018 | 0.79 |
| One Sentence: Imagery Use | -0.005 | 0.94 |
| **Memory Questionnaires** |  |  |
| One Sentence: Memory in Words | -0.19 | 0.004 |
| One Sentence: Memory in Imagery | 0.12 | 0.081 |
| One Sentence: Memory in Scene Imagery | 0.13 | 0.067 |
| One Sentence: Memory Ability | -0.085 | 0.21 |
| **Future Thinking Questionnaires** |  |  |
| One Sentence: Future Thinking Ability | 0.13 | 0.052 |
| One Sentence: Future Thinking in Words | -0.13 | 0.059 |
| One Sentence: Future Thinking in Scene Imagery | 0.055 | 0.42 |
| One Sentence: Future Thinking in Imagery | -0.044 | 0.52 |

#### 3.5.4. Navigation summary

The main associations between the questionnaires and navigation task performance are shown in Figure 4.

**Figure 4.**
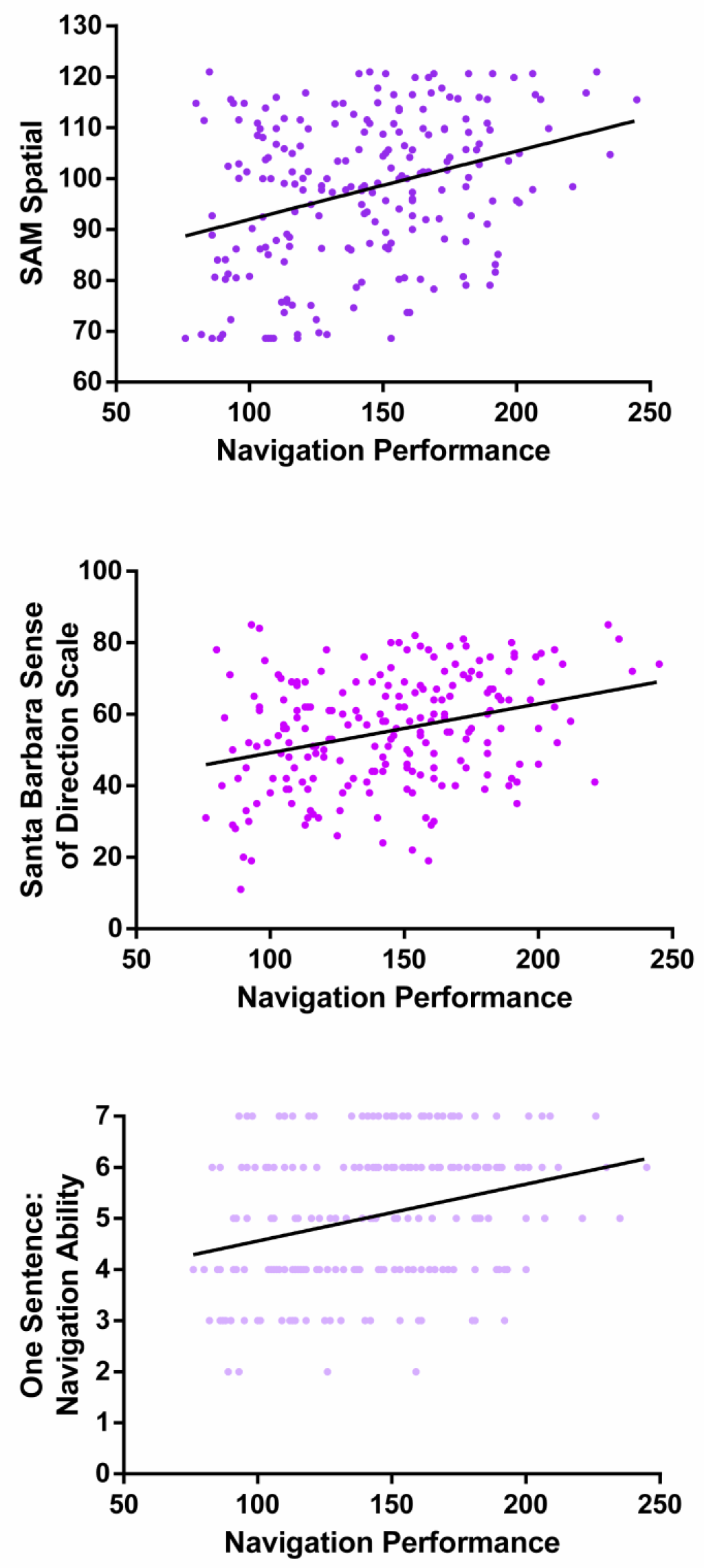
Graphs showing the correlations between the SAM Spatial subscale, the Santa Barbara Sense of Direction Scale and the One Sentence: Navigation Ability question with performance on the navigation task.

Considering our three research questions, first, navigation questionnaires reflect navigation performance. Second, associations with the navigation task seem to be specific to when individuals were asked about their navigation ability. This is in contrast to imagination, autobiographical memory vividness and future thinking, which seemed to be more related to the imagery questionnaires. Third, the One Sentence: Navigation Ability question performed at a comparable level as the established navigation questionnaires.

## 4. Discussion

Information about an individual’s thoughts and beliefs is a valuable resource in psychology, contributing unique data to help us understand cognition and behaviour. Questionnaires are one of the best methodologies for accessing this information. However, there is a surprising lack of empirical work examining whether questionnaires actually reflect their purported cognitive functions where performance on naturalistic tasks is concerned. Here, we showed that widely used visual imagery (imagination) and navigation questionnaires reflected their cognitive functions, but that autobiographical memory and future thinking questionnaires did not. Instead, we found that autobiographical memory questionnaires were associated with ratings of autobiographical memory vividness. Furthermore, we showed that as well as imagery questionnaires being most associated with the scene construction imagination task, they also had the greatest association with autobiographical memory vividness and future thinking. Finally, we highlighted the potential of our exploratory One Sentence Questionnaire for using single questions to capture a broad profile of a participant, with these questions performing comparably to the established in-depth questionnaires in this sample of healthy, young adults.

Three of the five questionnaires that purported to measure visual imagery were associated with performance on the scene construction task (our naturalistic measure of imagination): the PSIQ Appearance subscale (Andrade et al., 2014), the OSIVQ Object-Scene subscale (Blazhenkova & Kozhevnikov, 2009) and the SUIS (Reisberg et al., 2003). Of these, the PSIQ had the greatest association, likely due to its similarity to the imagination task in question. On the other hand, the Visualizer questionnaire (Kirby et al., 1988) and the OSIVQ Spatial subscale were not associated with scene construction task performance. Comparison of the questionnaires reveals why this might be the case. The PSIQ, OSIVQ Object-Scene subscale and SUIS all focus on imagery of pictures and scenes, while the Visualizer and OSIVQ Spatial subscale focus on more technical, abstract and schematic visual imagery representing locations, movement and transformations. Imagination performance, at least as measured by scene construction, seems, therefore, to only be associated with imagery questionnaires that focus on concrete visual imagery.

The two questionnaires designed to measure navigation ability, the Santa Barbara Sense of Direction Scale (Hegarty et al., 2002) and the SAM Spatial subscale (Palombo et al., 2013), were both associated with performance on the navigation task, in line with previous studies (Hegarty et al., 2006; Ishikawa & Montello, 2006; Selarka et al., 2019; Weisberg et al., 2014). In addition, these questionnaires were associated with navigation performance to the same extent, and both seemed to be tapping into the same construct. Overall, therefore, these results permit the conclusion that these widely used navigation questionnaires reflect their purported cognitive function.

By contrast, the memory questionnaires that we examined were not associated with autobiographical memory ability as measured by AI internal (episodic) details. Six commonly used memory questionnaires were included; four subscales of the MEQ (Sutin & Robins, 2007), the SMQ (Bennett-Levy & Powell, 1980), and the SAM Episodic subscale (Palombo et al., 2013). We are not the first to note the lack of correlation between memory questionnaires and performance measures of memory. A review of 14 memory questionnaires as far back as 1982 found limited relationships between any of the memory questionnaires and laboratory measures of memory performance (Herrmann, 1982). In addition, a more recent study also failed to find a relationship between the SAM Episodic subscale and AI internal details (Palombo et al., 2013).

However, memory questionnaires were not completely divorced from autobiographical memory measures. Instead, with the exception of the MEQ Sharing subscale, all of the memory questionnaires correlated with autobiographical memory vividness. It may, therefore, be that questionnaire measures of memory actually reflect an individual’s ability to visualise their memories, rather than the number of details they can explicitly produce.

Future thinking is a relatively new area of formal study, and we were only able to identify and include one future thinking questionnaire – the SAM Future subscale (Palombo et al., 2013). However, here, the SAM Future subscale was not associated with performance on the future thinking task. To the best of our knowledge, this is the first study to compare responses on the SAM Future subscale with a measure of future thinking. Further work is, therefore, required to understand the relationship between questionnaire measures of future thinking and future thinking task performance.

As well as examining whether questionnaires reflected their purported cognitive functions, we also investigated if a visual imagery or autobiographical memory process is actually what is being assessed by questionnaires, despite their purporting to be related to distinct cognitive functions. To do this, we looked at the associations between performance on each task and the questionnaires associated with the other cognitive function groups. This identified an overall role for imagery in imagination, autobiographical memory vividness and future thinking. The imagery questionnaires were associated with all three tasks, and best reflected performance, over and above that of the memory and future thinking questionnaires. This result aligns with previous findings of a link between subjective reports of imagery and elements of autobiographical memory and future thinking (D’Argembeau & Van der Linden, 2006; Vannucci et al., 2016). It also mirrors our finding of a key role for scene imagery in performance on imagination, autobiographical memory and future thinking tasks (Clark et al., in press).

By contrast, there was a different pattern of associations between questionnaires and navigation scores. Only one non-navigation questionnaire was associated with navigation task performance - the OSIVQ Spatial subscale. While nominally an imagery questionnaire, the OSIVQ Spatial subscale was not associated with performance on imagination, autobiographical memory (internal details or vividness) or future thinking tasks, and instead was only linked to navigation. In other words, the OSIVQ Spatial subscale followed the same pattern of results as the navigation questionnaires. Of note, scores on the OSIVQ Spatial subscale have previously been associated with performance on “spatial” laboratory tasks such as paper folding and mental rotation (Blazhenkova & Kozhevnikov, 2009), tasks which have also been linked to navigation performance (Clark et al., in press). In line with the aims of the OSIVQ, the Spatial subscale is, therefore, reflecting a different cognitive process to that of the OSIVQ Object-Scene subscale, with the OSVIQ Spatial subscale related to navigation, while the OSIVQ Object-Scene subscale reflects imagination, autobiographical memory vividness and future thinking.

The lack of an association between the memory questionnaires and AI internal details was a surprise, particularly as memory questionnaires and AI internal details are such widely used measures of autobiographical memory ability (and indeed are often used interchangeably), and so we might expect them to be related. Moving forward, it will be important to tease apart exactly what each of these test instruments is in fact measuring.

According to the scoring procedure of the AI, internal details are those that could reasonably reflect episodic re-experiencing (Levine et al., 2002; Palombo, Alain, Söderlund, Khuu, & Levine, 2015; but see also Strikwerda-Brown, Mothakunnel, Hodges, Piguet, & Irish, in press). The idea behind this is to remove any experimenter subjectivity when scoring, and so provide an unbiased estimate of the amount of episodic information in the memory as described by an individual. This methodology has been widely used and has been shown to be sensitive to individual differences in healthy compared to pathological aging (e.g. Grilli, Wank, Bercel, & Ryan, 2018; Murphy, Troyer, Levine, & Moscovitch, 2008) and damage to the medial temporal lobe, in particular the hippocampus (e.g. McCormick, Rosenthal, Miller, & Maguire, 2018; Rosenbaum et al., 2008; Steinvorth, Levine, & Corkin, 2005). In addition, in young, healthy participants, AI internal details has been associated with combined CA2/3 dentate gyrus volume in the left hippocampus (Palombo, Bacopulos, et al., 2018) and measures of fornix white matter (Hodgetts et al., 2017). However, while AI internal details may reflect some aspects of individual differences in autobiographical memory recall, the lack of an association between the memory questionnaires and AI internal details suggests that they might not reflect an individual’s subjective experience of memory recall.

Instead, the memory questionnaires were associated with AI vividness, suggesting that the subjective experience of memory is particularly related to the vividness with which a memory is recalled. This has an interesting parallel with findings in people with “severely deficient autobiographical memory”, where there was no difference compared to matched controls in terms of the number of internal details for autobiographical memories aged 1 week, 1 month, 1 year and 10 years old, but instead there were lower vividness ratings (Palombo et al., 2015; Palombo, Sheldon, & Levine, 2018). Furthermore, comparisons of true and false memories have found that the latter are less vivid (Heaps & Nash, 2001), whereas emotionally salient memories are typically highly vivid events (Rubin & Kozin, 1984; Talarico, LaBar, & Rubin, 2004). Vividness, therefore, seems to be central to the autobiographical memory recall experience.

Overall, memory questionnaires and AI internal details seem to be measuring different things. On the one hand, this may reflect the multifaceted nature of autobiographical memory recall. However, going forward, it will be important to no longer use memory questionnaires and AI internal detail measures interchangeably. Individual differences identified when using memory questionnaires (e.g. Sheldon et al., 2016) are unlikely to reflect “episodic memory” in the same manner as individual differences identified by AI internal details. Indeed, the results presented here suggest that memory questionnaire findings are related to memory vividness and, consequently, that when an individual is commenting upon their memory, it may be that they are actually referring to the extent to which they can visualise their memories, rather than the amount of detail they can recall. Which measure is most appropriate will likely depend upon the research question being asked.

Finally, we were also interested in whether a broad profile of a participant could be obtained by using a quick-to-administer set of single questions on each cognitive domain of interest. We compared the performance of our exploratory One Sentence Questionnaire with that of the established questionnaires. This identified a number of interesting parallels. For imagery, the three one sentence questions performed at a similar level to the OSIVQ Object-Scene and SUIS questionnaires. The PSIQ, however, outperformed the One Sentence Questionnaire, as it did the other established questionnaires. For autobiographical memory, in line with the established questionnaires, none of the memory questions from the One Sentence Questionnaire were associated with AI internal details. However, the imagery based memory questions were associated with autobiographical memory vividness, and this was to the same extent as the established memory questionnaires. For future thinking, as with the established questionnaire, the future thinking One Sentence Questionnaire questions did not correlate with performance. Finally, for navigation, the navigation ability question of the One Sentence Questionnaire correlated with navigation performance, and did so to the same extent as the established questionnaires. We observed, therefore, a parallel in the patterns of correlations between our exploratory One Sentence Questionnaire and the established questionnaires.

Overall, our exploratory One Sentence Questionnaire suggests that a broad profile of information about a person, that is comparable to established questionnaires, can be obtained quickly and easily using only single questions on a specific topic. Moreover, it seems that even our One Sentence Questionnaire can be further shortened, with only two questions required to predict performance on imagination, autobiographical memory (vividness), future thinking and navigation; one on imagery ability (reflecting imagination, autobiographical memory vividness and future thinking) and one on navigation ability. We emphasise, however, that this was an exploratory aspect of the study and additional work is needed in relation to this questionnaire, for example, replication in a separate sample of young, healthy participants and in other groups (e.g. clinical groups, older adults), and tests of validity (e.g., test-retest data) are also required. However, this initial exploration suggests that, while in-depth questionnaires certainly have their place, if a broad overview is required in a large sample of young, healthy volunteers, then asking a small number of single questions may be an efficient and accurate methodology in some circumstances.

In this study we were not able to include every published questionnaire relating to our cognitive functions of interest, nor every naturalistic cognitive task. However, those we examined here are widely used and, consequently, we believe they are representative of the extant literature.

In summary, the current study conveys four messages. First, imagination and navigation questionnaires reflect performance on naturalistic task measures of their purported cognitive functions. Second, memory questionnaires are associated with autobiographical memory vividness and not AI internal (episodic) details. Third, imagery questionnaires are better associated with autobiographical memory vividness and future thinking than the questionnaires purporting to reflect these functions. Finally, initial exploratory analyses (requiring replication) suggest that a broad profile of information may be obtained efficiently using a small number of simple single questions, and these model task performance comparably to the established questionnaires in the current sample of young, healthy adults. Overall, while some questionnaires can act as proxies for behaviour, the relationships between memory and future thinking tasks and questionnaires are more complex and require further elucidation.

## Acknowledgements

Thanks to Anna Monk, Victoria Hotchin, Gloria Pizzamiglio and Alice Liefgreen for assistance with data collection and scoring. Thanks also to Brian Levine for the materials and scoring details of the Survey of Autobiographical Memory questionnaire.

## Data availability

The data will be made freely available in the future once the construction of a dedicated data-sharing portal has been finalised. In the meantime, requests for the data can be sent to.

## Supplementary Materials

### Methods

#### Scene construction test

(Hassabis, Kumaran, Vann, & Maguire, 2007). This test measures a participant’s ability to mentally construct a visual scene. Participants construct seven different scenes of commonplace settings. For each scene, a short cue is provided (e.g. imagine lying on a beach in a beautiful tropical bay), and the participant is asked to imagine the scene that is evoked and then describe it out loud in as much detail as possible. Participants are explicitly told not to describe a memory, but to create a new scene that they have never experienced before. Participants give descriptions until they come to a natural end or cannot add any additional details. If required, a probing protocol is utilised to attempt to elicit more details (if a description is particularly poor); these are either very general probes (e.g. is there anything else you can tell me?) or based upon a theme described by the participant. The experimenter is never allowed to introduce new concepts or details that have not been mentioned by the participant. All descriptions are audio recorded and transcribed for scoring.

Following each scene description, participants rate their imagined scene in terms of their feeling of sense of presence (1 - ‘did not feel like I was there at all’…5 - ‘felt strongly like I was really there’) and perceived vividness (1 - ‘couldn’t really see anything’…5 - ‘extremely vivid’). They also rate the spatial coherence of the scene, choosing from twelve statements providing possible qualitative descriptions of the scene. Participants are instructed to indicate the statements they feel accurately described their construction. Eight of these statements describe a spatially coherent scene (e.g. ‘I could see the whole scene in my mind’s eye’) and 4 describe a fragmented scene (e.g. ‘It was a collection of separate images’). Participants can choose as many or as few options as they feel were relevant to the scene.

The overall outcome measure from the scene construction test is an “experiential index” which is calculated for each scene and then averaged. It is composed of four elements: numerical scoring of the content, participant ratings of their sense of presence (how much they felt like they were really there) and perceived vividness, participant ratings of the spatial coherence of the scene, and an experimenter rating of overall quality of the scene. To obtain the content score, the transcribed versions of each scene are split into statements. These are then classified by the experimenter as belonging to one of four categories: spatial references, entities present, sensory descriptions, or thoughts/emotions/actions. A maximum of seven details per category can be awarded (providing a maximum content score of 28). The original study reporting this test determined that seven details per category is enough to create a coherent scene without over-rewarding more verbose participants (Hassabis et al., 2007).

The participant ratings are the scores provided for sense of presence and perceived vividness, rescaled to 0-4, providing a maximum total of 8. Spatial coherence is from the qualitative statements that participants chose following each scene. One point is awarded for each coherent statement selected and one point taken away for each fragmented statement. This yields a score between –4 and +8 that is then normalised around zero to give a final spatial coherence index score ranging between –6 and +6. Only positive spatial coherence scores are included in the experiential index so as not to over penalise fragmented descriptions.

Finally, the experimenter assesses the overall quality of the scene on a scale between 0-10 (0 indicating a scene with no details and 10 a rich and vivid description). This was rescored to provide a rating from 0 to 18. The final experiential index score therefore ranged from 0 to 60; 28 points from the content, 8 for the participant ratings, 6 for the spatial coherence statements and 18 for the quality rating. An overall experiential index score was calculated by averaging across the 7 scenes.

Double scoring was performed on the data from 20% of the participants (i.e. 44 participants, which equates to 308 individual scenes). Scenes were scored by four experimenters, with double scoring performed on 20% of each experimenter’s scoring. We took the most stringent approach to identifying across-experimenter agreement. An inter-class correlation coefficient, with a two-way random effects model looking for absolute agreement, was calculated for each content score and for the quality ratings. This was performed both for individual scenes and as an average of all seven scenes across each participant. All inter-class correlation coefficients were above 0.9 (full details in Table S1). For reference, a score of 0.8 or above is considered excellent agreement beyond chance.

**Table S1.** Double scoring of the scene construction test.

|  |  | Rating |  |  |  |
| --- | --- | --- | --- | --- | --- |
|  | Spatial<br>References | Entities<br>present | Sensory<br>Descriptions | Thoughts/<br>Emotions/Actions | Quality<br>Ratings |
| **For each individual scene** |  |  |  |  |  |
| n = 308 | 0.90 | 0.96 | 0.94 | 0.90 | 0.90 |
| **For each individual participant (i.e. the score is averaged across the seven scenes)** |  |  |  |  |  |
| n = 44 | 0.91 | 0.99 | 0.97 | 0.91 | 0.93 |

#### Autobiographical Interview

(AI; Levine, Svoboda, Hay, Winocur, & Moscovitch, 2002). The AI asks participants to recall and describe autobiographical memories from a specific time and place over four time periods – early childhood (up to age 11), teenage years (from 11-17 years of age), adulthood (from aged 18 up to 12 months prior to the interview; two memories were requested) and the last year (a memory from the last 12 months). Participants are asked to avoid selecting a memory from the last 12 months in the adulthood category to ensure that only the last year category contained memories from the last 12 months. If a participant cannot spontaneously produce a memory, a cue card containing ∼100 typical life events is presented to aid recall.

Participants are asked to select events that they are comfortable to talk about. They are told that the event has to be one where they were personally involved and of which they have a clear recollection (they could not just have been told about the event by others). All memories have to be from a specific time and place – an event for example could not be a two week summer holiday, but a specific event on that holiday would be acceptable. Participants are first asked to simply describe and speak about the event selected. This occurs without interruption from the experimenter until they have reached a natural end point. On completion, the experimenter prompts the participant with a general probe (e.g. is there anything more you can tell me?) to ascertain if any additional details can be elicited. All memories are audio recorded and transcribed for analysis.

For scoring, each memory is divided into segments of information. Segments are defined as a specific occurrence, observation or thought. Segments are then defined as either “internal” or “external”. Internal details are those describing the event in question. Internal details are split into five sub-categories: events (a specific detail of the story), places, time, perceptual information and emotions/thoughts. The number of details from each sub-category is summed to create the overall “internal details” measure. External details describe semantic information concerning the event, or non-event information. The number of internal and external details was combined to provide a total number of utterances, allowing for the number of internal details to be represented as a percentage of the total number of utterances to control for verbosity.

Double scoring was performed on the data from 20% of the participants (i.e. 44 participants, which equates to 215 individual memories). Memories were scored by three experimenters, with double scoring performed on 20% of each experimenter’s scoring. An inter-class correlation coefficient, with a two-way random effects model looking for absolute agreement was calculated for each internal detail sub-category, and the overall internal and external details. This was performed both for individual memories and as an average of all five memories across each participant. The majority of inter-class correlation coefficients were above 0.9, with the lowest at 0.81 (full details in Table S2).

**Table S2.** Double scoring of the Autobiographical Interview.

|  | Rating |  |  |  |  |  |  |
| --- | --- | --- | --- | --- | --- | --- | --- |
|  | Internal<br>Event | Internal<br>Place | Internal<br>Time | Internal<br>Perceptual | Internal<br>Emotion | Internal<br>Details | External<br>Details |
| **For each individual memory** |  |  |  |  |  |  |  |
| n = 215 | .92 | .85 | .94 | .92 | .86 | **.94** | **.84** |
| **For each individual participant (i.e. score is averaged across the five memories)** |  |  |  |  |  |  |  |
| n = 43 | .95 | .88 | .96 | .94 | .81 | **.97** | **.87** |

#### Future thinking test

(Hassabis et al., 2007). Double scoring was performed on the data from 20% of the participants (i.e. 44 participants, which equates to 132 future scenes). Data were scored by four experimenters, with double scoring performed on 20% of each experimenter’s scoring – see Table S3.

**Table S3.** Double scoring of the future thinking test.

| Rating |  |  |  |  |  |
| --- | --- | --- | --- | --- | --- |
|  | Spatial<br>References | Entities<br>present | Sensory<br>Descriptions | Thoughts/<br>Emotions/Actions | Quality<br>ratings |
| **For each individual scene** |  |  |  |  |  |
| n = 132 | 0.90 | 0.94 | 0.93 | 0.88 | 0.90 |
| **For each individual participant (i.e. the score is averaged across the three future events)** |  |  |  |  |  |
| n = 44 | 0.94 | 0.95 | 0.96 | 0.88 | 0.92 |

#### Navigation tests

(Woollett & Maguire, 2010). Navigation ability was assessed using movies of navigation through an unfamiliar town. Movie clips of two overlapping routes through this real town (Blackrock, in Dublin, Ireland) are shown to the participant four times. Footage is unique to each route apart from one crossover point at a large road junction. The footage was shot at eye level and proceeds at an average walking pace, with the camera panning from side to side to simulate viewing and to allow for observation of the features and landmarks along the routes. At road junctions, the pace is slowed to allow for all elements of the junction to be observed before continuing. The movies are shown without sound. Participants are told to focus on salient landmarks to help them learn the town, ignoring cars, buses and people. Landmarks are defined as prominent buildings and distinctive elements of the route. Participants are explicitly told that the two routes, while shown separately, overlapped.

Five tasks are used to assess the participant’s ability to learn the new town, with an overall navigation score calculated by combining the scores from all the tasks. First, following each viewing of the route movies, participants are shown four short clips – two from the actual routes, and two distractors. Participants indicate whether they had seen that clip or not. The final score (/16) is the number of correctly identified clips.

Second, after all four route viewings are completed, recognition memory for scenes from the routes is tested. Participants are shown 32 photographs, 24 from the routes (12 from each route) and 8 similar distractors, randomly intermixed. Participants have to report whether they had seen that scene or not. The final score (/32) is the number of correctly identified scenes.

A third test involves assessing knowledge of the spatial relationships between landmarks from the routes. On each trial, three colour photographs of landmarks are presented and participants have to judge which of two of the landmarks was closer, as the crow flies, to the third picture (i.e. the target landmark). Ten trials are conducted, 6 where the landmarks are all from the same route (3 from each route) and 4 where the landmarks are from across the two routes. The final score is the number of correct judgements (/10).

Fourth, route knowledge is examined by having participants place photographs from the routes in the correct order as if travelling through the town. On each trial, participants are given eight photographs, one marked as the “start point” and another as the “end point”. They are then asked to place the other six photographs in the correct order to get from the start to the end. Four trials are performed, two that remain within one route and two that involve both routes. Correctly placed photographs are given a score of 1 (the maximum being 24).

Finally, participants draw a sketch map of the two routes including as many landmarks as they can remember (with it being made clear that drawing ability was not being assessed). Sketch maps are scored in terms of:

- The number of road segments (a segment being the section of road between road junctions, out of 16)
- The number of road junctions (out of 8)
- The number of correct landmarks (out of 34 identifiable landmarks)
- Landmark placement (up to 3 points per landmark; 1 for the correct side of the road, 1 for placement with regards to nearby road junctions and 1 for being in the correct sequence of nearby landmarks. Total maximum of 102)
- The orientation of the routes (an experimenter score assessing the orientation and layout of the map from 1-5. 1 being poor representation of the town, 5 being accurate orientation)
- An overall map categorisation score (an experimenter score representing map coherence from 1-6; 1: The two routes are merged. 2: Two routes are present, but drawn separately. 3: Routes are close together but not joined accurately. 4: Some elements of the routes are linked, but integration is mainly lacking. 5: The two routes are integrated, but with some inaccuracies. 6: Correct integration, easy to follow and use for navigating.

Sketch maps were scored by three experimenters, with double scoring performed on 20% of each experimenter’s scoring (n = 42 maps). All the inter-class correlation coefficients were above 0.89 (Table S4).

**Table S4.** Double scoring of the navigation sketch maps.

|  | Rating |  |  |  |  |  |
| --- | --- | --- | --- | --- | --- | --- |
|  | Road Segments | Road Junctions | Number of Landmarks | Landmark Placement | Map Orientation | Map Categorisation |
| n = 42 | 0.95 | 0.96 | 0.97 | 0.96 | 0.96 | 0.89 |

**Table S5.** Correlation coefficients of the memory questionnaires with AI internal details as a percentage of total utterances for each memory time point.

|  | Childhood |  | Teenage |  | “Remote” |  | Adult |  | Last Year |  | “Recent” |  |
| --- | --- | --- | --- | --- | --- | --- | --- | --- | --- | --- | --- | --- |
|  | r | p | r | p | r | p | r | p | r | p | r | p |
| Memory Experience Questionnaire; Accessibility | 0.015 | 0.82 | -0.053 | 0.44 | -0.023 | 0.74 | 0.13 | 0.048 | 0.063 | 0.36 | 0.12 | 0.074 |
| Memory Experience Questionnaire; Coherence | -0.030 | 0.66 | -0.21 | 0.002 | -0.14 | 0.039 | -0.14 | 0.042 | -0.094 | 0.17 | -0.14 | 0.043 |
| Memory Experience Questionnaire; Sharing | -0.017 | 0.80 | -0.048 | 0.48 | -0.039 | 0.57 | -0.004 | 0.95 | 0.12 | 0.071 | 0.048 | 0.48 |
| Memory Experience Questionnaire; Vividness | -0.002 | 0.98 | -0.083 | 0.23 | -0.051 | 0.46 | 0.005 | 0.94 | 0.002 | 0.98 | 0.004 | 0.95 |
| Subjective Memory Questionnaire | 0.054 | 0.43 | -0.080 | 0.24 | -0.017 | 0.80 | 0.005 | 0.95 | -0.001 | 0.99 | 0.003 | 0.97 |
| Survey of Autobiographical Memory; Episodic | -0.064 | 0.35 | -0.14 | 0.036 | -0.12 | 0.071 | -0.053 | 0.44 | -0.007 | 0.92 | -0.040 | 0.55 |

**Table S6.**
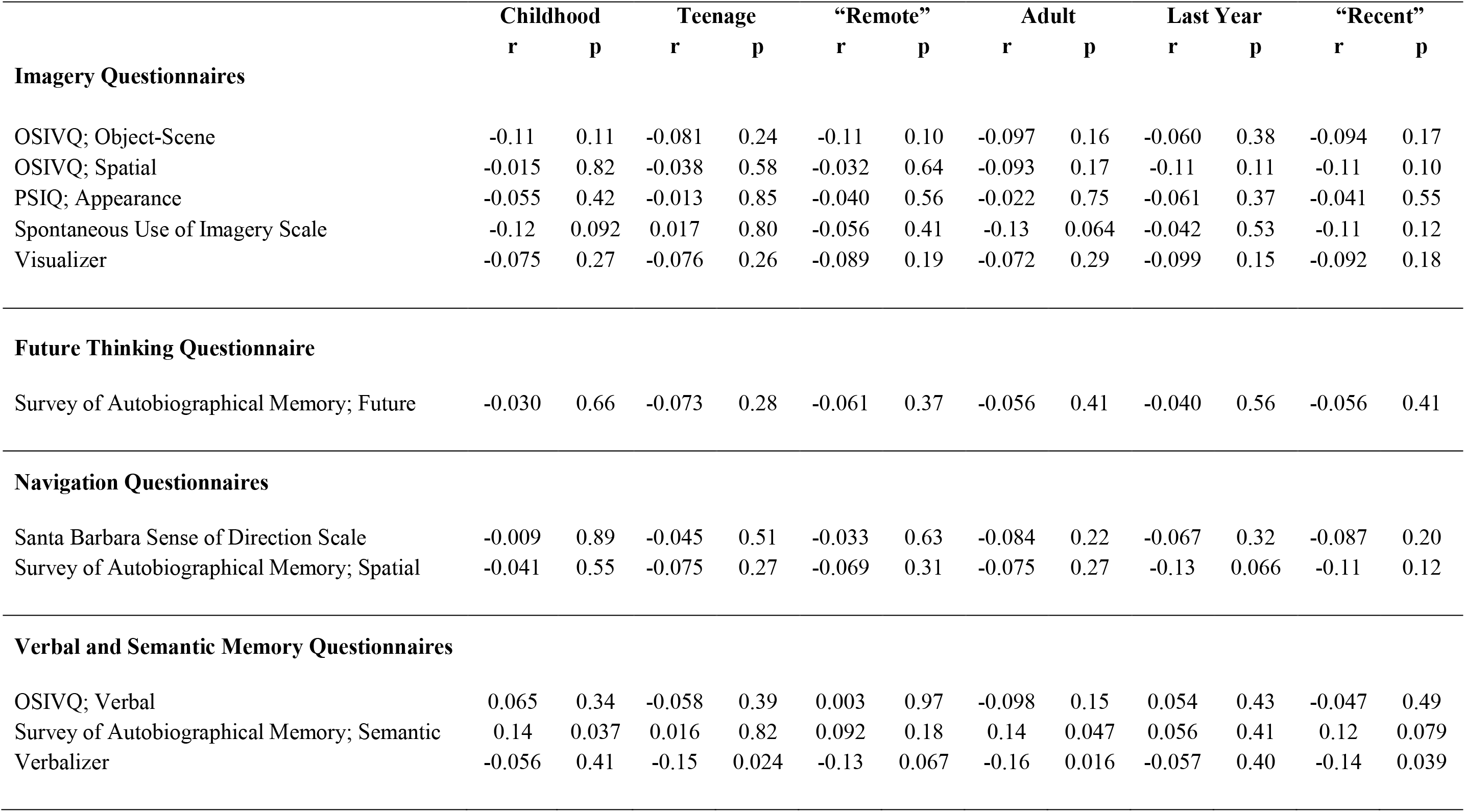
Correlation coefficients of the other group questionnaires with AI internal details as a percentage of total utterances for each memory time point.

|  | Childhood |  | Teenage |  | “Remote” |  | Adult |  | Last Year |  | “Recent” |  |
| --- | --- | --- | --- | --- | --- | --- | --- | --- | --- | --- | --- | --- |
|  | r | p | r | p | r | p | r | p | r | p | r | p |
| **Imagery Questionnaires** |  |  |  |  |  |  |  |  |  |  |  |  |
| OSIVQ; Object-Scene | -0.11 | 0.11 | -0.081 | 0.24 | -0.11 | 0.10 | -0.097 | 0.16 | -0.060 | 0.38 | -0.094 | 0.17 |
| OSIVQ; Spatial | -0.015 | 0.82 | -0.038 | 0.58 | -0.032 | 0.64 | -0.093 | 0.17 | -0.11 | 0.11 | -0.11 | 0.10 |
| PSIQ; Appearance | -0.055 | 0.42 | -0.013 | 0.85 | -0.040 | 0.56 | -0.022 | 0.75 | -0.061 | 0.37 | -0.041 | 0.55 |
| Spontaneous Use of Imagery Scale | -0.12 | 0.092 | 0.017 | 0.80 | -0.056 | 0.41 | -0.13 | 0.064 | -0.042 | 0.53 | -0.11 | 0.12 |
| Visualizer | -0.075 | 0.27 | -0.076 | 0.26 | -0.089 | 0.19 | -0.072 | 0.29 | -0.099 | 0.15 | -0.092 | 0.18 |
| **Future Thinking Questionnaire** |  |  |  |  |  |  |  |  |  |  |  |  |
| Survey of Autobiographical Memory; Future | -0.030 | 0.66 | -0.073 | 0.28 | -0.061 | 0.37 | -0.056 | 0.41 | -0.040 | 0.56 | -0.056 | 0.41 |
| **Navigation Questionnaires** |  |  |  |  |  |  |  |  |  |  |  |  |
| Santa Barbara Sense of Direction Scale | -0.009 | 0.89 | -0.045 | 0.51 | -0.033 | 0.63 | -0.084 | 0.22 | -0.067 | 0.32 | -0.087 | 0.20 |
| Survey of Autobiographical Memory; Spatial | -0.041 | 0.55 | -0.075 | 0.27 | -0.069 | 0.31 | -0.075 | 0.27 | -0.13 | 0.066 | -0.11 | 0.12 |
| **Verbal and Semantic Memory Questionnaires** |  |  |  |  |  |  |  |  |  |  |  |  |
| OSIVQ; Verbal | 0.065 | 0.34 | -0.058 | 0.39 | 0.003 | 0.97 | -0.098 | 0.15 | 0.054 | 0.43 | -0.047 | 0.49 |
| Survey of Autobiographical Memory; Semantic | 0.14 | 0.037 | 0.016 | 0.82 | 0.092 | 0.18 | 0.14 | 0.047 | 0.056 | 0.41 | 0.12 | 0.079 |
| Verbalizer | -0.056 | 0.41 | -0.15 | 0.024 | -0.13 | 0.067 | -0.16 | 0.016 | -0.057 | 0.40 | -0.14 | 0.039 |

**Table S7.**
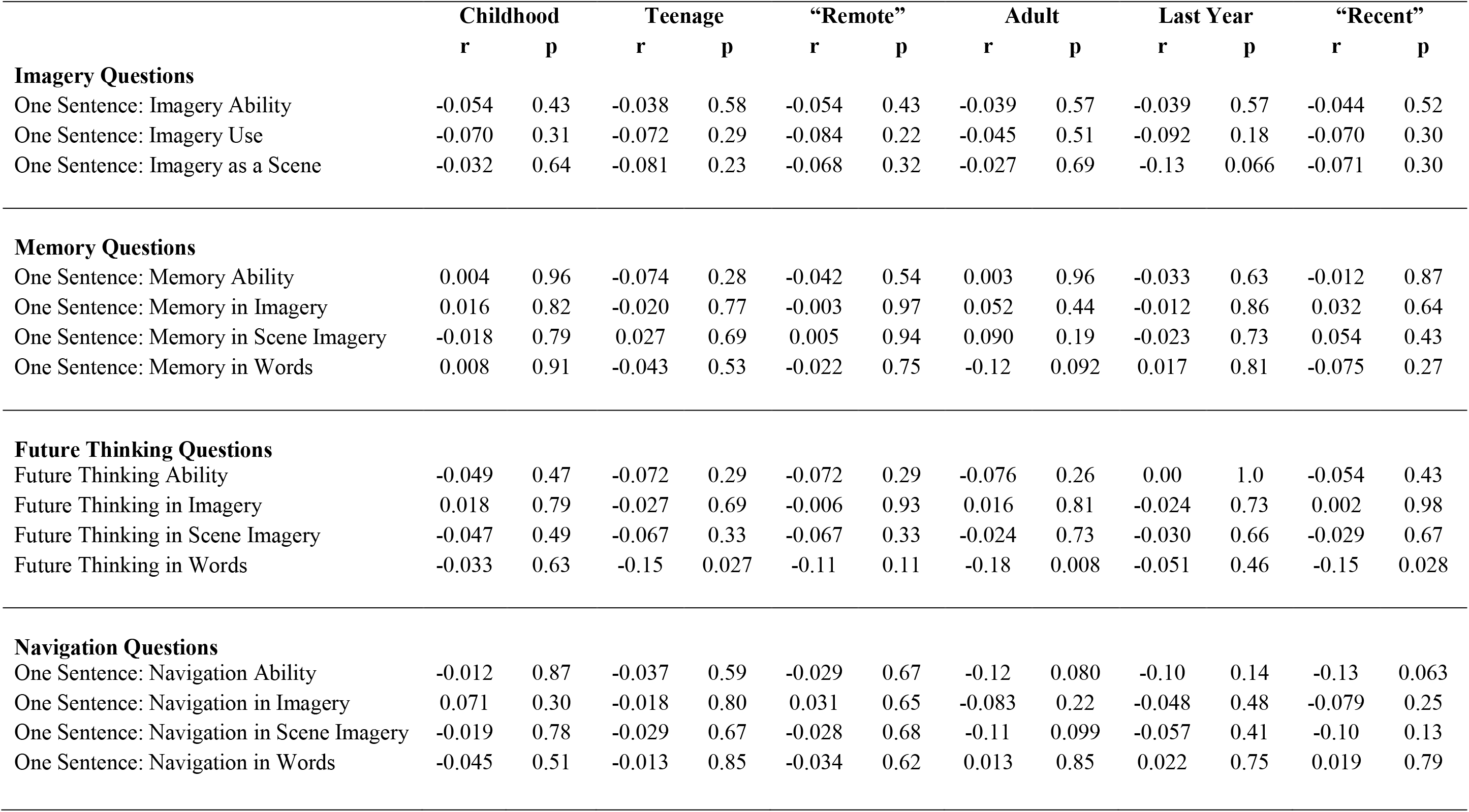
Correlation coefficients of the One Sentence Questionnaire questions with AI internal details as a percentage of total utterances for each memory time point.

|  | Childhood |  | Teenage |  | “Remote” |  | Adult |  | Last Year |  | “Recent” |  |
| --- | --- | --- | --- | --- | --- | --- | --- | --- | --- | --- | --- | --- |
|  | r | p | r | p | r | p | r | p | r | p | r | p |
| **Imagery Questions** |  |  |  |  |  |  |  |  |  |  |  |  |
| One Sentence: Imagery Ability | -0.054 | 0.43 | -0.038 | 0.58 | -0.054 | 0.43 | -0.039 | 0.57 | -0.039 | 0.57 | -0.044 | 0.52 |
| One Sentence: Imagery Use | -0.070 | 0.31 | -0.072 | 0.29 | -0.084 | 0.22 | -0.045 | 0.51 | -0.092 | 0.18 | -0.070 | 0.30 |
| One Sentence: Imagery as a Scene | -0.032 | 0.64 | -0.081 | 0.23 | -0.068 | 0.32 | -0.027 | 0.69 | -0.13 | 0.066 | -0.071 | 0.30 |
| **Memory Questions** |  |  |  |  |  |  |  |  |  |  |  |  |
| One Sentence: Memory Ability | 0.004 | 0.96 | -0.074 | 0.28 | -0.042 | 0.54 | 0.003 | 0.96 | -0.033 | 0.63 | -0.012 | 0.87 |
| One Sentence: Memory in Imagery | 0.016 | 0.82 | -0.020 | 0.77 | -0.003 | 0.97 | 0.052 | 0.44 | -0.012 | 0.86 | 0.032 | 0.64 |
| One Sentence: Memory in Scene Imagery | -0.018 | 0.79 | 0.027 | 0.69 | 0.005 | 0.94 | 0.090 | 0.19 | -0.023 | 0.73 | 0.054 | 0.43 |
| One Sentence: Memory in Words | 0.008 | 0.91 | -0.043 | 0.53 | -0.022 | 0.75 | -0.12 | 0.092 | 0.017 | 0.81 | -0.075 | 0.27 |
| **Future Thinking Questions** |  |  |  |  |  |  |  |  |  |  |  |  |
| Future Thinking Ability | -0.049 | 0.47 | -0.072 | 0.29 | -0.072 | 0.29 | -0.076 | 0.26 | 0.00 | 1.0 | -0.054 | 0.43 |
| Future Thinking in Imagery | 0.018 | 0.79 | -0.027 | 0.69 | -0.006 | 0.93 | 0.016 | 0.81 | -0.024 | 0.73 | 0.002 | 0.98 |
| Future Thinking in Scene Imagery | -0.047 | 0.49 | -0.067 | 0.33 | -0.067 | 0.33 | -0.024 | 0.73 | -0.030 | 0.66 | -0.029 | 0.67 |
| Future Thinking in Words | -0.033 | 0.63 | -0.15 | 0.027 | -0.11 | 0.11 | -0.18 | 0.008 | -0.051 | 0.46 | -0.15 | 0.028 |
| **Navigation Questions** |  |  |  |  |  |  |  |  |  |  |  |  |
| One Sentence: Navigation Ability | -0.012 | 0.87 | -0.037 | 0.59 | -0.029 | 0.67 | -0.12 | 0.080 | -0.10 | 0.14 | -0.13 | 0.063 |
| One Sentence: Navigation in Imagery | 0.071 | 0.30 | -0.018 | 0.80 | 0.031 | 0.65 | -0.083 | 0.22 | -0.048 | 0.48 | -0.079 | 0.25 |
| One Sentence: Navigation in Scene Imagery | -0.019 | 0.78 | -0.029 | 0.67 | -0.028 | 0.68 | -0.11 | 0.099 | -0.057 | 0.41 | -0.10 | 0.13 |
| One Sentence: Navigation in Words | -0.045 | 0.51 | -0.013 | 0.85 | -0.034 | 0.62 | 0.013 | 0.85 | 0.022 | 0.75 | 0.019 | 0.79 |

**Table S8.** Correlation coefficients of the memory questionnaires with the AI internal detail sub-categories as a percentage of total utterances.

|  | AI Events |  | AI Place |  | AI Time |  | AI Perceptual |  | AI Emotion |  |
| --- | --- | --- | --- | --- | --- | --- | --- | --- | --- | --- |
|  | r | p | r | p | r | p | r | p | r | p |
| Memory Experience Questionnaire; Accessibility | 0.071 | 0.30 | -0.12 | 0.085 | 0.098 | 0.15 | 0.013 | 0.85 | 0.023 | 0.74 |
| Memory Experience Questionnaire; Coherence | -0.074 | 0.28 | -0.12 | 0.077 | 0.12 | 0.068 | -0.009 | 0.90 | -0.12 | 0.076 |
| Memory Experience Questionnaire; Sharing | 0.079 | 0.25 | -0.19 | 0.006 | 0.015 | 0.83 | 0.009 | 0.89 | 0.021 | 0.75 |
| Memory Experience Questionnaire; Vividness | 0.025 | 0.71 | -0.16 | 0.017 | 0.073 | 0.29 | -0.044 | 0.52 | 0.056 | 0.41 |
| Subjective Memory Questionnaire | -0.051 | 0.46 | -0.053 | 0.44 | 0.099 | 0.15 | 0.055 | 0.42 | -0.038 | 0.58 |
| Survey of Autobiographical Memory; Episodic | -0.004 | 0.95 | -0.16 | 0.021 | 0.11 | 0.11 | -0.049 | 0.47 | -0.023 | 0.74 |

**Table S9.** Correlation coefficients of the other group questionnaires with the AI internal detail sub-categories as a percentage of total utterances.

|  | AI Events |  | AI Place |  | AI Time |  | AI Perceptual |  | AI Emotion |  |
| --- | --- | --- | --- | --- | --- | --- | --- | --- | --- | --- |
|  | r | p | r | p | r | p | r | p | r | p |
| **Imagery Questionnaires** |  |  |  |  |  |  |  |  |  |  |
| OSIVQ; Object-Scene | -0.015 | 0.83 | -0.15 | 0.024 | 0.078 | 0.25 | -0.084 | 0.22 | -0.003 | 0.97 |
| OSIVQ; Spatial | -0.035 | 0.60 | -0.027 | 0.70 | -0.004 | 0.95 | -0.12 | 0.070 | 0.084 | 0.22 |
| PSIQ; Appearance | 0.014 | 0.84 | -0.16 | 0.016 | 0.11 | 0.11 | -0.012 | 0.86 | -0.050 | 0.47 |
| Spontaneous Use of Imagery Scale | -0.022 | 0.75 | -0.13 | 0.054 | 0.006 | 0.93 | -0.074 | 0.28 | 0.043 | 0.53 |
| Visualizer | -0.074 | 0.28 | -0.015 | 0.83 | -0.033 | 0.63 | -0.12 | 0.078 | 0.13 | 0.054 |
| **Future Thinking Questionnaire** |  |  |  |  |  |  |  |  |  |  |
| Survey of Autobiographical Memory; Future | -0.077 | 0.26 | -0.13 | 0.064 | 0.029 | 0.67 | 0.002 | 0.98 | 0.043 | 0.53 |
| **Navigation Questionnaires** |  |  |  |  |  |  |  |  |  |  |
| Santa Barbara Sense of Direction Scale | -0.085 | 0.21 | -0.052 | 0.45 | 0.12 | 0.091 | 0.040 | 0.56 | -0.089 | 0.19 |
| Survey of Autobiographical Memory; Spatial | -0.075 | 0.27 | -0.092 | 0.18 | 0.099 | 0.15 | -0.008 | 0.91 | -0.062 | 0.36 |
| **Verbal and Semantic Memory Questionnaires** |  |  |  |  |  |  |  |  |  |  |
| OSIVQ; Verbal | -0.14 | 0.047 | -0.046 | 0.50 | 0.066 | 0.33 | 0.11 | 0.12 | -0.040 | 0.56 |
| Survey of Autobiographical Memory; Semantic | 0.070 | 0.31 | -0.14 | 0.035 | 0.13 | 0.049 | 0.077 | 0.26 | -0.013 | 0.84 |
| Verbalizer | -0.24 | < 0.001 | -0.048 | 0.49 | 0.042 | 0.53 | 0.072 | 0.29 | -0.011 | 0.87 |

**Table S10.**
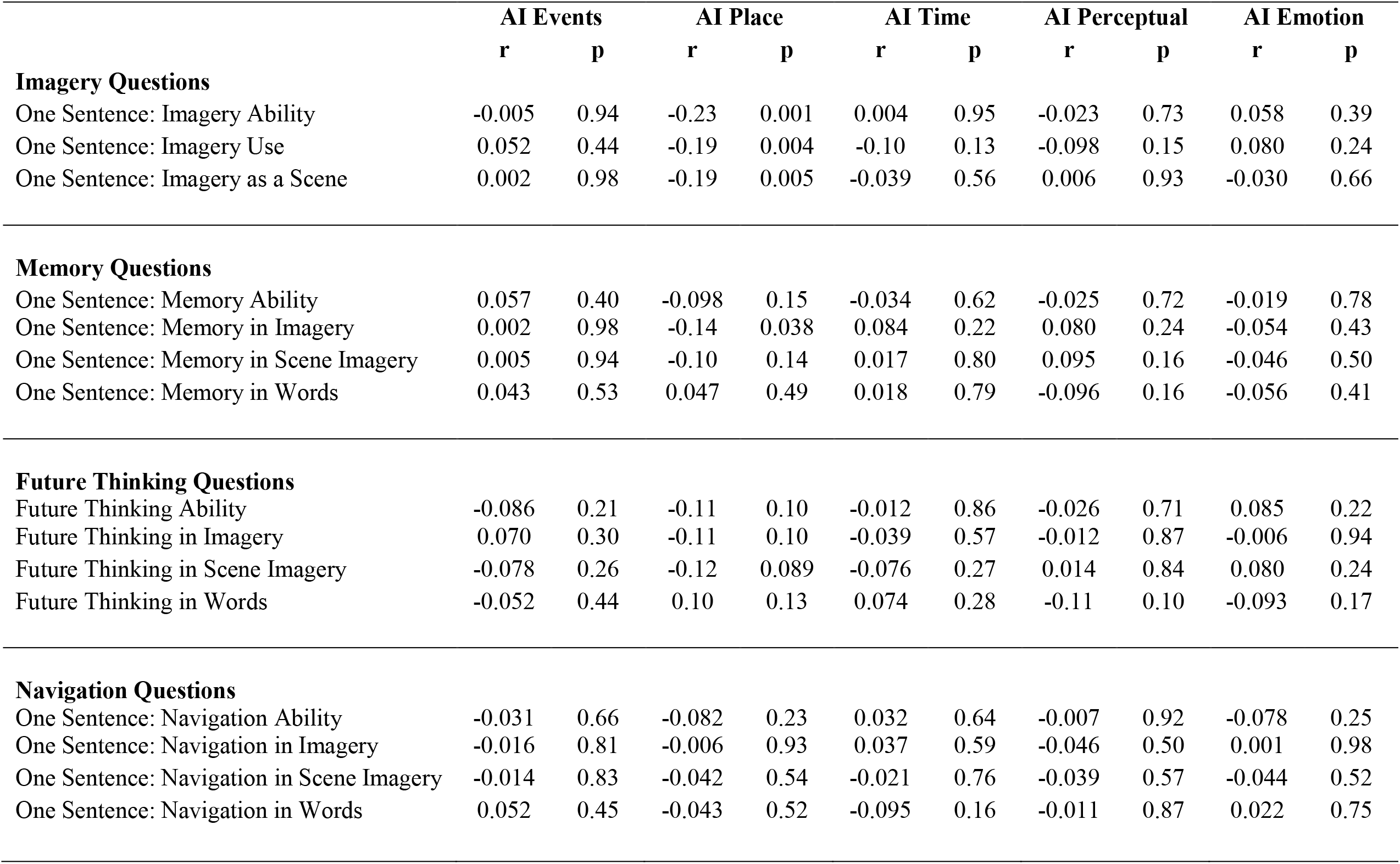
Correlation coefficients of the One Sentence Questionnaire questions with the AI internal detail sub-categories as a percentage of total utterances.

|  | AI Events |  | AI Place |  | AI Time |  | AI Perceptual |  | AI Emotion |  |
| --- | --- | --- | --- | --- | --- | --- | --- | --- | --- | --- |
|  | r | p | r | p | r | p | r | p | r | p |
| **Imagery Questions** |  |  |  |  |  |  |  |  |  |  |
| One Sentence: Imagery Ability | -0.005 | 0.94 | -0.23 | 0.001 | 0.004 | 0.95 | -0.023 | 0.73 | 0.058 | 0.39 |
| One Sentence: Imagery Use | 0.052 | 0.44 | -0.19 | 0.004 | -0.10 | 0.13 | -0.098 | 0.15 | 0.080 | 0.24 |
| One Sentence: Imagery as a Scene | 0.002 | 0.98 | -0.19 | 0.005 | -0.039 | 0.56 | 0.006 | 0.93 | -0.030 | 0.66 |
| **Memory Questions** |  |  |  |  |  |  |  |  |  |  |
| One Sentence: Memory Ability | 0.057 | 0.40 | -0.098 | 0.15 | -0.034 | 0.62 | -0.025 | 0.72 | -0.019 | 0.78 |
| One Sentence: Memory in Imagery | 0.002 | 0.98 | -0.14 | 0.038 | 0.084 | 0.22 | 0.080 | 0.24 | -0.054 | 0.43 |
| One Sentence: Memory in Scene Imagery | 0.005 | 0.94 | -0.10 | 0.14 | 0.017 | 0.80 | 0.095 | 0.16 | -0.046 | 0.50 |
| One Sentence: Memory in Words | 0.043 | 0.53 | 0.047 | 0.49 | 0.018 | 0.79 | -0.096 | 0.16 | -0.056 | 0.41 |
| **Future Thinking Questions** |  |  |  |  |  |  |  |  |  |  |
| Future Thinking Ability | -0.086 | 0.21 | -0.11 | 0.10 | -0.012 | 0.86 | -0.026 | 0.71 | 0.085 | 0.22 |
| Future Thinking in Imagery | 0.070 | 0.30 | -0.11 | 0.10 | -0.039 | 0.57 | -0.012 | 0.87 | -0.006 | 0.94 |
| Future Thinking in Scene Imagery | -0.078 | 0.26 | -0.12 | 0.089 | -0.076 | 0.27 | 0.014 | 0.84 | 0.080 | 0.24 |
| Future Thinking in Words | -0.052 | 0.44 | 0.10 | 0.13 | 0.074 | 0.28 | -0.11 | 0.10 | -0.093 | 0.17 |
| **Navigation Questions** |  |  |  |  |  |  |  |  |  |  |
| One Sentence: Navigation Ability | -0.031 | 0.66 | -0.082 | 0.23 | 0.032 | 0.64 | -0.007 | 0.92 | -0.078 | 0.25 |
| One Sentence: Navigation in Imagery | -0.016 | 0.81 | -0.006 | 0.93 | 0.037 | 0.59 | -0.046 | 0.50 | 0.001 | 0.98 |
| One Sentence: Navigation in Scene Imagery | -0.014 | 0.83 | -0.042 | 0.54 | -0.021 | 0.76 | -0.039 | 0.57 | -0.044 | 0.52 |
| One Sentence: Navigation in Words | 0.052 | 0.45 | -0.043 | 0.52 | -0.095 | 0.16 | -0.011 | 0.87 | 0.022 | 0.75 |

**Table S11.** Correlation coefficients of the memory questionnaires with the number of AI internal details for each memory time point.

|  | Childhood |  | Teenage |  | “Remote” |  | Adult |  | Last Year |  | “Recent” |  |
| --- | --- | --- | --- | --- | --- | --- | --- | --- | --- | --- | --- | --- |
|  | r | p | r | p | r | p | r | p | r | p | r | p |
| Memory Experience Questionnaire; Accessibility | 0.18 | 0.007 | 0.061 | 0.37 | 0.14 | 0.041 | 0.19 | 0.005 | 0.091 | 0.18 | 0.17 | 0.012 |
| Memory Experience Questionnaire; Coherence | 0.14 | 0.045 | 0.010 | 0.89 | 0.083 | 0.22 | 0.089 | 0.19 | -0.016 | 0.81 | 0.053 | 0.44 |
| Memory Experience Questionnaire; Sharing | 0.20 | 0.004 | 0.11 | 0.12 | 0.17 | 0.011 | 0.25 | < 0.001 | 0.20 | 0.003 | 0.26 | < 0.001 |
| Memory Experience Questionnaire; Vividness | 0.16 | 0.018 | 0.069 | 0.31 | 0.13 | 0.054 | 0.15 | 0.026 | 0.062 | 0.36 | 0.13 | 0.055 |
| Subjective Memory Questionnaire | 0.16 | 0.015 | 0.042 | 0.54 | 0.12 | 0.083 | 0.12 | 0.078 | -0.035 | 0.61 | 0.066 | 0.34 |
| Survey of Autobiographical Memory; Episodic | 0.14 | 0.033 | 0.079 | 0.25 | 0.13 | 0.062 | 0.13 | 0.057 | 0.010 | 0.89 | 0.092 | 0.18 |

**Table S12.** Correlation coefficients of the other group questionnaires with the number of AI internal details for each memory time point.

|  | Childhood |  | Teenage |  | “Remote” |  | Adult |  | Last Year |  | “Recent” |  |
| --- | --- | --- | --- | --- | --- | --- | --- | --- | --- | --- | --- | --- |
|  | r | p | r | p | r | p | r | p | r | p | r | p |
| **Imagery Questionnaires** |  |  |  |  |  |  |  |  |  |  |  |  |
| OSIVQ; Object-Scene | 0.14 | 0.041 | 0.013 | 0.85 | 0.087 | 0.20 | 0.067 | 0.33 | -0.066 | 0.33 | 0.016 | 0.82 |
| OSIVQ; Spatial | -0.072 | 0.29 | -0.035 | 0.61 | -0.061 | 0.37 | -0.15 | 0.029 | -0.11 | 0.094 | -0.15 | 0.026 |
| PSIQ; Appearance | 0.079 | 0.25 | 0.055 | 0.42 | 0.076 | 0.26 | 0.079 | 0.24 | -0.028 | 0.68 | 0.041 | 0.55 |
| Spontaneous Use of Imagery Scale | 0.11 | 0.094 | 0.028 | 0.68 | 0.081 | 0.24 | 0.031 | 0.65 | 0.011 | 0.87 | 0.026 | 0.70 |
| Visualizer | -0.036 | 0.60 | -0.069 | 0.31 | -0.059 | 0.39 | -0.14 | 0.038 | -0.13 | 0.055 | -0.15 | 0.023 |
| **Future Thinking Questionnaire** |  |  |  |  |  |  |  |  |  |  |  |  |
| Survey of Autobiographical Memory; Future | 0.070 | 0.30 | 0.058 | 0.39 | 0.073 | 0.28 | -0.018 | 0.79 | -0.098 | 0.15 | -0.056 | 0.41 |
| **Navigation Questionnaires** |  |  |  |  |  |  |  |  |  |  |  |  |
| Santa Barbara Sense of Direction Scale | 0.091 | 0.18 | 0.037 | 0.59 | 0.073 | 0.28 | -0.018 | 0.79 | -0.098 | 0.15 | -0.056 | 0.41 |
| Survey of Autobiographical Memory; Spatial | 0.055 | 0.42 | 0.026 | 0.71 | 0.046 | 0.50 | -0.037 | 0.58 | -0.097 | 0.16 | -0.069 | 0.31 |
| **Verbal and Semantic Memory Questionnaires** |  |  |  |  |  |  |  |  |  |  |  |  |
| OSIVQ; Verbal | 0.18 | 0.009 | 0.18 | 0.007 | 0.20 | 0.003 | 0.18 | 0.010 | 0.054 | 0.43 | 0.14 | 0.035 |
| Survey of Autobiographical Memory; Semantic | 0.15 | 0.025 | 0.092 | 0.18 | 0.14 | 0.041 | 0.14 | 0.034 | 0.013 | 0.85 | 0.10 | 0.13 |
| Verbalizer | 0.055 | 0.42 | 0.019 | 0.78 | 0.042 | 0.54 | 0.10 | 0.13 | 0.013 | 0.85 | 0.075 | 0.27 |

**Table S13.** Correlation coefficients of the One Sentence Questionnaire questions with the number of AI internal details for each memory time point.

|  | Childhood |  | Teenage |  | “Remote” |  | Adult |  | Last Year |  | “Recent” |  |
| --- | --- | --- | --- | --- | --- | --- | --- | --- | --- | --- | --- | --- |
|  | r | p | r | p | r | p | r | p | r | p | r | p |
| **Imagery Questions** |  |  |  |  |  |  |  |  |  |  |  |  |
| One Sentence: Imagery Ability | 0.14 | 0.038 | 0.065 | 0.34 | 0.12 | 0.085 | 0.072 | 0.29 | -0.037 | 0.59 | 0.032 | 0.64 |
| One Sentence: Imagery Use | 0.097 | 0.15 | 0.032 | 0.64 | 0.074 | 0.28 | 0.060 | 0.38 | -0.050 | 0.47 | 0.019 | 0.75 |
| One Sentence: Imagery as a Scene | 0.11 | 0.098 | 0.005 | 0.94 | 0.068 | 0.32 | 0.066 | 0.33 | -0.040 | 0.56 | 0.027 | 0.69 |
| **Memory Questions** |  |  |  |  |  |  |  |  |  |  |  |  |
| One Sentence: Memory Ability | 0.15 | 0.023 | 0.097 | 0.16 | 0.14 | 0.036 | 0.094 | 0.17 | 0.026 | 0.70 | 0.075 | 0.27 |
| One Sentence: Memory in Imagery | 0.18 | 0.007 | 0.074 | 0.28 | 0.15 | 0.032 | 0.082 | 0.23 | 0.068 | 0.32 | 0.086 | 0.21 |
| One Sentence: Memory in Scene Imagery | 0.11 | 0.11 | 0.073 | 0.28 | 0.10 | 0.13 | 0.066 | 0.33 | -0.061 | 0.37 | 0.018 | 0.79 |
| One Sentence: Memory in Words | -0.14 | 0.048 | 0.016 | 0.81 | -0.068 | 0.32 | -0.046 | 0.50 | -0.023 | 0.73 | -0.042 | 0.54 |
| **Future Thinking Questions** |  |  |  |  |  |  |  |  |  |  |  |  |
| Future Thinking Ability | 0.091 | 0.18 | 0.029 | 0.68 | 0.068 | 0.32 | -0.002 | 0.97 | -0.011 | 0.88 | -0.006 | 0.93 |
| Future Thinking in Imagery | 0.11 | 0.12 | 0.013 | 0.85 | 0.069 | 0.32 | 0.053 | 0.44 | 0.019 | 0.78 | 0.044 | 0.52 |
| Future Thinking in Scene Imagery | 0.062 | 0.36 | 0.037 | 0.59 | 0.056 | 0.41 | 0.007 | 0.92 | -0.045 | 0.51 | -0.016 | 0.82 |
| Future Thinking in Words | -0.038 | 0.58 | 0.048 | 0.48 | 0.005 | 0.94 | 0.025 | 0.71 | -0.031 | 0.65 | 0.003 | 0.96 |
| **Navigation Questions** |  |  |  |  |  |  |  |  |  |  |  |  |
| One Sentence: Navigation Ability | 0.13 | 0.060 | 0.062 | 0.36 | 0.11 | 0.11 | -0.048 | 0.48 | -0.034 | 0.62 | -0.048 | 0.48 |
| One Sentence: Navigation in Imagery | 0.078 | 0.25 | 0.055 | 0.42 | 0.076 | 0.27 | 0.018 | 0.79 | -0.040 | 0.55 | -0.006 | 0.93 |
| One Sentence: Navigation in Scene Imagery | 0.062 | 0.36 | 0.11 | 0.10 | 0.098 | 0.15 | 0.082 | 0.23 | 0.031 | 0.65 | 0.070 | 0.31 |
| One Sentence: Navigation in Words | -0.002 | 0.98 | 0.056 | 0.41 | 0.030 | 0.66 | 0.10 | 0.14 | 0.091 | 0.18 | 0.11 | 0.11 |

**Table S14.** Correlation coefficients of the memory questionnaires with the total number of AI internal details for each internal detail sub-category.

|  | AI Internal<br>Details |  | AI Events |  | AI Place |  | AI Time |  | AI<br>Perceptual |  | AI Emotion |  |
| --- | --- | --- | --- | --- | --- | --- | --- | --- | --- | --- | --- | --- |
|  | r | p | r | p | r | p | r | p | r | p | r | p |
| Memory Experience Questionnaire; Accessibility | 0.18 | 0.009 | 0.17 | 0.015 | 0.049 | 0.47 | 0.21 | 0.002 | 0.099 | 0.15 | 0.095 | 0.17 |
| Memory Experience Questionnaire; Coherence | 0.071 | 0.30 | 0.057 | 0.40 | 0.055 | 0.42 | 0.23 | 0.001 | 0.046 | 0.50 | -0.025 | 0.71 |
| Memory Experience Questionnaire; Sharing | 0.26 | < 0.001 | 0.25 | < 0.001 | 0.093 | 0.17 | 0.19 | 0.006 | 0.13 | 0.057 | 0.18 | 0.009 |
| Memory Experience Questionnaire; Vividness | 0.15 | 0.033 | 0.15 | 0.033 | 0.009 | 0.90 | 0.19 | 0.005 | 0.041 | 0.55 | 0.14 | 0.047 |
| Subjective Memory Questionnaire | 0.093 | 0.17 | 0.051 | 0.46 | 0.055 | 0.42 | 0.17 | 0.015 | 0.096 | 0.16 | 0.022 | 0.75 |
| Survey of Autobiographical Memory; Episodic | 0.12 | 0.088 | 0.12 | 0.087 | -0.01 | 0.89 | 0.22 | 0.001 | 0.045 | 0.51 | 0.063 | 0.35 |

**Table S15.** Correlation coefficients of the other group questionnaires with the total number of AI internal details for each internal detail sub-category.

|  | AI Internal<br>Details |  | AI Events |  | AI Place |  | AI Time |  | AI Perceptual |  | AI Emotion |  |
| --- | --- | --- | --- | --- | --- | --- | --- | --- | --- | --- | --- | --- |
|  | r | p | r | p | r | p | r | p | r | p | r | p |
| **Imagery Questionnaires** |  |  |  |  |  |  |  |  |  |  |  |  |
| OSIVQ; Object-Scene | 0.045 | 0.51 | 0.062 | 0.36 | -0.025 | 0.72 | 0.16 | 0.019 | -0.029 | 0.67 | 0.050 | 0.47 |
| OSIVQ; Spatial | -0.13 | 0.051 | -0.11 | 0.12 | -0.15 | 0.032 | -0.053 | 0.44 | -0.14 | 0.035 | 0.007 | 0.91 |
| PSIQ; Appearance | 0.060 | 0.38 | 0.062 | 0.37 | -0.084 | 0.22 | 0.16 | 0.021 | 0.039 | 0.57 | 0.011 | 0.87 |
| Spontaneous Use of Imagery Scale | 0.051 | 0.46 | 0.057 | 0.40 | -0.020 | 0.77 | 0.086 | 0.21 | -0.010 | 0.89 | 0.077 | 0.26 |
| Visualizer | -0.13 | 0.049 | -0.11 | 0.10 | -0.12 | 0.068 | -0.062 | 0.36 | -0.15 | 0.024 | 0.031 | 0.65 |
| **Future Thinking Questionnaire** |  |  |  |  |  |  |  |  |  |  |  |  |
| Survey of Autobiographical Memory; Future | 0.029 | 0.67 | 0.011 | 0.87 | -0.055 | 0.42 | 0.084 | 0.22 | 0.018 | 0.79 | 0.051 | 0.45 |
| **Navigation Questionnaires** |  |  |  |  |  |  |  |  |  |  |  |  |
| Santa Barbara Sense of Direction Scale | -0.012 | 0.86 | -0.034 | 0.62 | -0.054 | 0.43 | 0.13 | 0.053 | 0.038 | 0.57 | -0.076 | 0.26 |
| Survey of Autobiographical Memory; Spatial | -0.031 | 0.65 | -0.036 | 0.60 | -0.086 | 0.21 | 0.14 | 0.036 | -0.002 | 0.98 | -0.075 | 0.27 |
| **Verbal and Semantic Memory Questionnaires** |  |  |  |  |  |  |  |  |  |  |  |  |
| OSIVQ; Verbal | 0.18 | 0.007 | 0.089 | 0.19 | 0.12 | 0.079 | 0.18 | 0.007 | 0.20 | 0.003 | 0.093 | 0.17 |
| Survey of Autobiographical Memory; Semantic | 0.13 | 0.058 | 0.12 | 0.089 | -0.088 | 0.20 | 0.19 | 0.004 | 0.11 | 0.11 | 0.041 | 0.55 |
| Verbalizer | 0.071 | 0.30 | -0.032 | 0.64 | 0.066 | 0.34 | 0.14 | 0.047 | 0.12 | 0.089 | 0.086 | 0.21 |

**Table S16.** Correlation coefficients of the One Sentence Questionnaire questions with the total number of AI internal details for each internal detail sub-category.

|  | AI Internal<br>Details |  | AI Events |  | AI Place |  | AI Time |  | AI Perceptual |  | AI Emotion |  |
| --- | --- | --- | --- | --- | --- | --- | --- | --- | --- | --- | --- | --- |
|  | r | p | r | p | r | p | r | p | r | p | r | p |
| **Imagery Questions** |  |  |  |  |  |  |  |  |  |  |  |  |
| One Sentence: Imagery Ability | 0.069 | 0.31 | 0.066 | 0.33 | -0.099 | 0.15 | 0.081 | 0.24 | 0.021 | 0.76 | 0.11 | 0.10 |
| One Sentence: Imagery Use | 0.042 | 0.53 | 0.098 | 0.15 | -0.10 | 0.13 | -0.034 | 0.62 | -0.045 | 0.51 | 0.091 | 0.18 |
| One Sentence: Imagery as a Scene | 0.046 | 0.50 | 0.072 | 0.29 | -0.074 | 0.28 | 0.032 | 0.64 | 0.021 | 0.76 | 0.009 | 0.89 |
| **Memory Questions** |  |  |  |  |  |  |  |  |  |  |  |  |
| One Sentence: Memory Ability | 0.11 | 0.11 | 0.13 | 0.054 | 0.044 | 0.52 | 0.070 | 0.31 | 0.052 | 0.45 | 0.032 | 0.64 |
| One Sentence: Memory in Imagery | 0.12 | 0.081 | 0.083 | 0.22 | 0.010 | 0.88 | 0.18 | 0.009 | 0.12 | 0.085 | 0.029 | 0.67 |
| One Sentence: Memory in Scene Imagery | 0.053 | 0.44 | 0.034 | 0.62 | -0.038 | 0.58 | 0.068 | 0.32 | 0.089 | 0.19 | -0.026 | 0.70 |
| One Sentence: Memory in Words | -0.057 | 0.41 | -0.013 | 0.85 | 0.008 | 0.91 | -0.005 | 0.94 | -0.090 | 0.19 | -0.051 | 0.46 |
| **Future Thinking Questions** |  |  |  |  |  |  |  |  |  |  |  |  |
| Future Thinking Ability | 0.022 | 0.75 | 0.003 | 0.96 | -0.061 | 0.37 | 0.035 | 0.61 | 0.006 | 0.93 | 0.086 | 0.21 |
| Future Thinking in Imagery | 0.059 | 0.39 | 0.10 | 0.14 | -0.023 | 0.74 | 0.012 | 0.86 | 0.005 | 0.94 | 0.016 | 0.81 |
| Future Thinking in Scene Imagery | 0.011 | 0.88 | -0.006 | 0.93 | -0.061 | 0.37 | -0.035 | 0.61 | 0.004 | 0.95 | 0.089 | 0.19 |
| Future Thinking in Words | 0.004 | 0.95 | 0.016 | 0.82 | 0.16 | 0.023 | 0.085 | 0.21 | -0.048 | 0.49 | -0.023 | 0.74 |
| **Navigation Questions** |  |  |  |  |  |  |  |  |  |  |  |  |
| One Sentence: Navigation Ability | 0.008 | 0.91 | 0.011 | 0.88 | -0.013 | 0.85 | 0.076 | 0.27 | 0.013 | 0.85 | -0.040 | 0.56 |
| One Sentence: Navigation in Imagery | 0.026 | 0.71 | 0.019 | 0.78 | 0.071 | 0.30 | 0.079 | 0.25 | -0.015 | 0.83 | 0.035 | 0.61 |
| One Sentence: Navigation in Scene Imagery | 0.089 | 0.19 | 0.10 | 0.14 | 0.096 | 0.16 | 0.064 | 0.35 | 0.028 | 0.68 | 0.037 | 0.59 |
| One Sentence: Navigation in Words | 0.090 | 0.19 | 0.12 | 0.070 | 0.023 | 0.74 | -0.078 | 0.25 | 0.022 | 0.75 | 0.087 | 0.20 |

## One Sentence Questionnaire

1. Please rate your ability to construct a mental image (circle one) 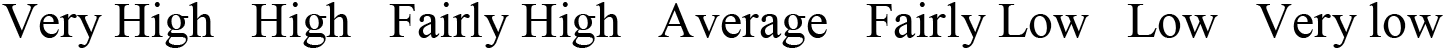
2. In everyday life, how much do you think in images (e.g. thinking in pictures in your mind) 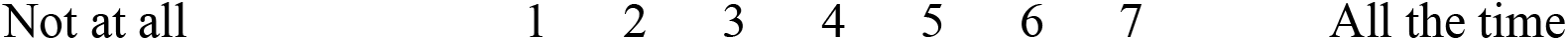
3. If you think in images, to what extent does this involve spatially coherent scenes (e.g. scenes that you could step into or operate within) compared to single objects? 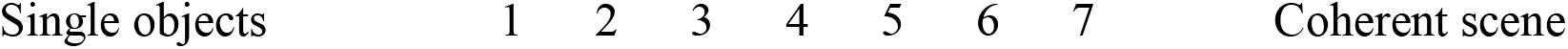
4. Please rate your ability to remember your personal past (circle one) 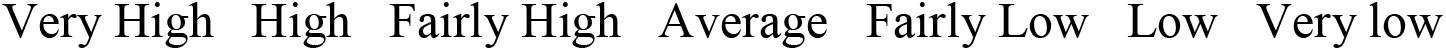
5. When recalling the past, to what extent do you think in images 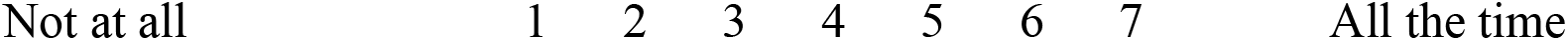
6. If you think in images, when recalling the past to what extent do you evoke spatially coherent scenes in your mind’s eye, compared to imagining single objects 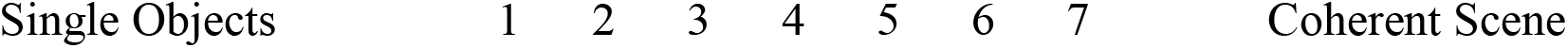
7. When recalling the past, how much do you think verbally (e.g. thinking in words and sentences) 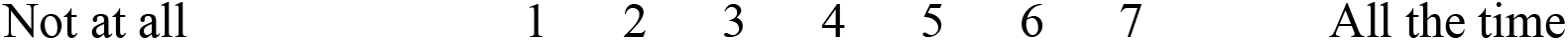
8. Please rate your ability to imagine future events (circle one) 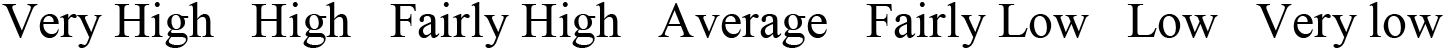
9. When imagining the future, to what extent do you think in images 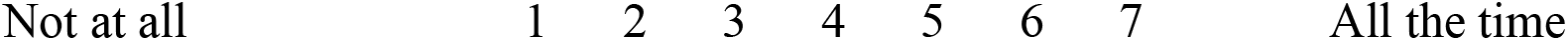
10. If you think in images, when imagining the future to what extent do you evoke spatially coherent scenes in your mind’s eye, compared to imagining single objects 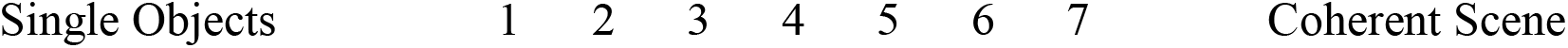
11. When imagining the future, how much do you think verbally (e.g. thinking in words and sentences) 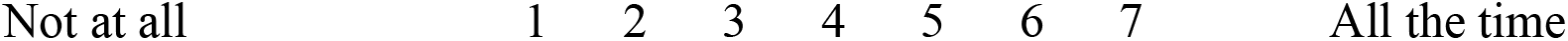
12. Please rate your navigational ability (circle one) 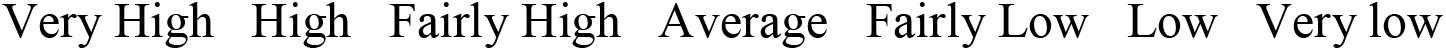
13. When you navigate, to what extent do you think in images 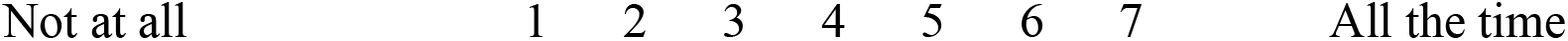
14. If you think in images, when navigating to what extent do you evoke spatially coherent scenes in your mind’s eye, compared to imagining single objects 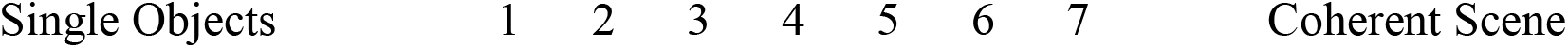
15. When navigating, how much do you think verbally (e.g. thinking in words and sentences) Not at all 1 2 3 4 5 6 7 All the time 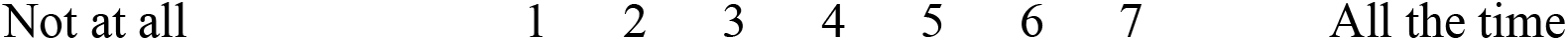

## Highlights

- Imagery and navigation questionnaires reflect their purported cognitive functions
- Memory questionnaires reflect autobiographical memory vividness
- Episodic details and memory questionnaires measure different aspects of memory
- Imagery questionnaires also correlated with memory vividness and future thinking
- Single questions modelled performance comparably to established questionnaires

